# Presymptomatic neuroanatomical and cognitive biomarkers of alpha-synuclein propagation in a mouse model of synucleinopathy

**DOI:** 10.1101/2022.10.12.511820

**Authors:** Stephanie Tullo, Aline S Miranda, Esther del Cid-Pellitero, Mei Peng Lim, Daniel Gallino, Anoosha Attaran, Raihaan Patel, Vladislav Novikov, Megan Park, Flavio H. Beraldo, Wen Luo, Irina Shlaifer, Thomas M. Durcan, Timothy J. Bussey, Lisa M. Saksida, Edward A. Fon, Vania F. Prado, Marco A.M. Prado, M. Mallar Chakravarty

**Affiliations:** Integrated Program in Neuroscience, McGill University, Montreal, Quebec, Canada; Computational Brain Anatomy (CoBrA) Laboratory, Cerebral Imaging Center, Douglas Mental Health University Institute, McGill University, Verdun, Quebec, Canada; Robarts Research Institute, Schulich School of Medicine, The University of Western Ontario, Ontario, Canada; Departamento de Morfologia, Instituto de Ciencias Biologicas, Universidade Fedearal de Minas Gerais, Brazil; McGill Parkinson Program, Neurodegenerative Diseases Group, Department of Neurology and Neurosurgery, Montreal Neurological Institute-Hospital, McGill University, Montreal, Québec, Canada; Department of Biological & Biomedical Engineering, McGill University, Montreal, Quebec, Canada; Early Drug Discovery Unit, Montreal Neurological Institute, McGill University, Montreal, Quebec, Canada; Department of Physiology and Pharmacology, Schulich School of Medicine, The University of Western Ontario, Ontario, Canada; Department of Anatomy & Cell Biology, Schulich School of Medicine, The University of Western Ontario, Ontario, Canada; Department of Psychiatry, McGill University, Montreal, Quebec, Canada

**Keywords:** alpha-synuclein, Parkinson’s disease, mouse model, neuroimaging, cognition, prion, prion-like, Lewy-body

## Abstract

There is significant evidence suggesting aggregated misfolded alpha-synuclein, a major component of Lewy bodies, propagates in a prion-like manner contributing to disease progression in Parkinson’s disease (PD) and other synucleinopathies. Animal models are essential for understanding and developing treatments for these diseases. However, despite modelling human pathology, most endpoints studied in mice do not translate to humans. Furthermore, the progression by which alpha-synuclein misfolding affects human-relevant measures such as brain volume and underlying subtle, high-level cognitive deficits is poorly understood. Here we used a mouse model of synucleinopathy; hemizygous M83 human A53T alpha-synuclein transgenic mice inoculated with recombinant human alpha-synuclein preformed fibrils (PFF) injected in the right striatum to initiate alpha-synuclein misfolding and aggregation. We examined alpha-synuclein-induced atrophy at 90 days post-injection using *ex vivo* magnetic resonance imaging as well as high-level cognition and motor function, as biomarkers of alpha-synuclein toxicity. We observed widespread atrophy in bilateral regions that project to or receive input from the injection site, highlighting a network of regions that are consistent with structural changes observed in humans with PD. Moreover, we detected early deficits in reversal learning with touchscreen testing in PFF-injected mice prior to motor dysfunction, consistent with the pathology observed in cortical-striatal and thalamic loops. We show, using translational approaches in mice, that progression of prion-like spreading of alpha-synuclein causes selective atrophy via connected brain regions leading to high-level cognitive deficits. We propose that precise imaging and cognitive biomarkers can provide a more direct and human-relevant measurement of alpha-synuclein-induced toxicity in pre-clinical testing.

**Significance Statement:** The work described in this manuscript showcases the utility of state-of-the-art methodologies (magnetic resonance imaging and touchscreen behavioural tasks) to examine endophenotypes, both in terms of symptomatology and neuroanatomy, of alpha-synuclein propagation in a mouse model of synucleinopathy. Our work further validates the M83-Hu-PFF mouse model of synucleinopathy-associated pathogenesis of neurodegenerative diseases while highlighting precise imaging and cognitive biomarkers of protein misfolding toxicity. Specifically, we identified rapid and translational biomarkers that can serve as a proxy for the direct examination of cellular levels for pathology. We anticipate that these biomarkers can measure progression of toxicity, specifically in the early phases, and may be more reliable than end stage pathology and more useful as endpoints in the examination of novel therapeutics.

## 1. Introduction

Parkinson’s disease (PD) is a progressive neurodegenerative disorder characterised by motor and non-motor features, such as resting tremor, bradykinesia, muscular rigidity, autonomic dysfunction, cognitive deficits, and late-stage dementia (1,2). The neuropathological staging of disease progression and diagnosis is largely dependent on the presence and location of alpha-synuclein (aSyn)-immunoreactive Lewy neurites (LNs) and Lewy bodies (LBs) (3). While the precise role of aSyn in PD pathogenesis remains unresolved, several studies have provided support for a role for prion-like transmission of aSyn toxicity, which suggests neurodegeneration is mediated by cell-to-cell spreading of pathogenic proteins. This hypothesis, which may generalize to other neurodegenerative conditions, postulates that certain proteins that misfold act as templates to induce further maladaptive conformational changes in their normal counterparts, increasing their propensity to become toxic and aggregated (4–13). Several lines of evidence support the aSyn spreading hypothesis in the context of synucleinopathies such as PD, Lewy-Body dementia, and multiple-system atrophy. These include human post-mortem studies reporting endogenous aSyn aggregates appearing in transplanted fetal dopaminergic cells years after transplantation (14–15), as well as pathology spreading in cellular (7, 16–19) and animal models (21–28). Taken together these studies strongly support the hypothesis that aSyn propagates in a prion-like manner through the central nervous system via neuronal connections. However, whether the progressive spreading of misfolded aSyn is related to brain-region specific atrophy that underlies cognitive dysfunction is still unknown.

Clinical research in PD supports the hypothesis of prion-like spreading as indexed by the progression of atrophy (29–34) and mathematical models that use multi-modal integration to predict brain atrophy patterns (35–37). In this context, the use of advanced small animal magnetic resonance imaging (MRI) techniques allows for whole brain image assays that have homology with the clinic (38–44). Further, recent advances in animal behavioural testing use fully automated and standardized touchscreen tasks that are similar to human tests to investigate executive function in animal models (45–47)

Here, we seeded M83 hemizygous transgenic mice expressing human A53T aSyn mutant with human aSyn preformed fibrils (PFF) in the dorsal striatum and identified brain-region atrophy and connected network-like changes due to aSyn-induced spreading that are strikingly similar to brain-region atrophy in humans. Given the brain regions affected, we examined reversal learning in mice injected with PFFs and found progressive cognitive deficits that preceded motor dysfunction by weeks. These results provide evidence for robust and progressive cognitive and imaging changes due to aSyn misfolding and spreading in connected brain networks.

## 2. Results

We sought to evaluate the spreading of toxic aSyn concurrently using MRI, immunohistochemistry, and to determine if these signatures were related to motor behaviours and/or cognitive behaviours. To evaluate the toxic misfolded aSyn, we used 3-4-month-old hemizygous M83 aSynA53T transgenic mice injected with either human aSyn preformed fibrils (PFF), or phosphate buffered saline (PBS) in the right striatum. To this end, we examined MRI-derived atrophy (section 2.1) (n=15), cellular markers of pathology using immunohistochemistry (section 2.2) (n=14), and motor and cognitive behaviours (n = ~30) (in three separate cohorts of mice). Final numbers per group per experimental condition are presented in Table 1. Electron microscope characterization of the fibrils in terms of distribution per length can be seen in Supplementary Figure 1.

**Table 1.**
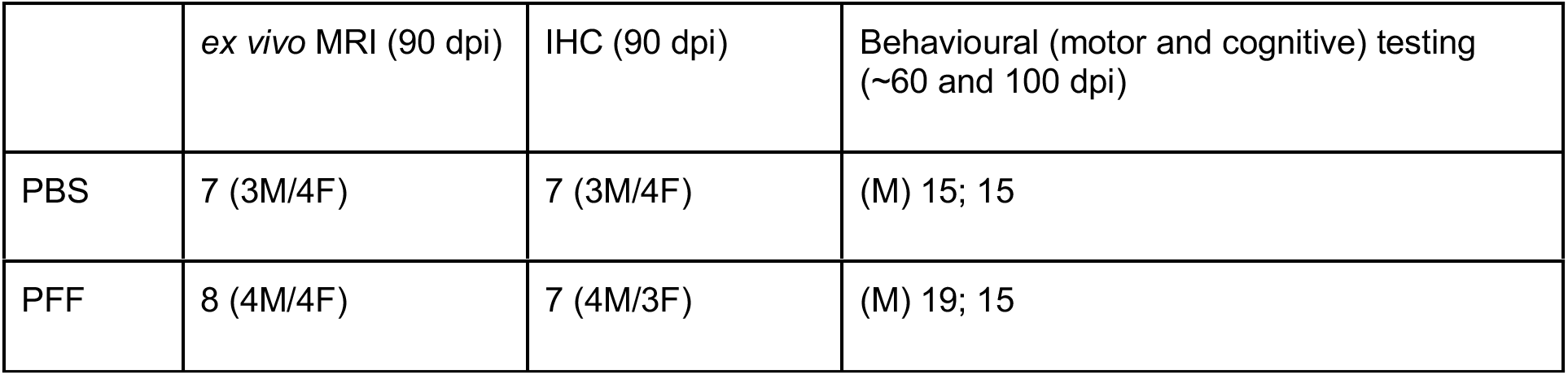
Sample per experimental condition. Phosphate buffered saline (PBS); human aSyn preformed fibrils (PFF); magnetic resonance imaging (MRI); immunohistochemistry (IHC); male (M); female (F).

|  | *ex vivo* MRI (90 dpi) | IHC (90 dpi) | Behavioural (motor and cognitive) testing (~60 and 100 dpi) |
| --- | --- | --- | --- |
| PBS | 7 (3M/4F) | 7 (3M/4F) | (M) 15; 15 |
| PFF | 8 (4M/4F) | 7 (4M/3F) | (M) 19; 15 |

### 2.1 MRI findings

To parallel human studies (48–49,34), we used high-resolution T1-weighted MRI images (70 μm^3^ isotropic voxels; Bruker 7T) acquired *ex vivo* at 90 days post-injection (dpi). We evaluated differences in brain anatomy using region-of-interest brain volumes automatically estimated through the Multiple Automatically Generated Templates (MAGeT)-Brain segmentation pipeline (50) using definitions from the Allen Brain Atlas (ABA) (51–52). Using a group-wise registration (deformation-based morphometry; DBM) technique to derive maps of the Jacobian determinants that were used to assess group differences in voxel-level volumetric expansion and compression (53–54; https://github.com/CoBrALab/twolevel_ants_dbm).

#### 2.1.1 Volumetric analysis

PFF-injected mice had smaller volumes across several regions with predominantly bilateral effect in comparison to the PBS-injected mice (control group) (Figure 1B). Differences were mainly observed in the key structures implicated in PD, such as the injection site (right striatum) (Figure 1C), right nucleus accumbens, the right substantia nigra pars compacta (Figure 1D), bilateral somatomotor cortices (Figure 1E), and right subthalamic nucleus. The volumetric differences were evaluated using a general linear model and corrected for multiple comparisons using a somewhat lenient threshold of 10% False Discovery Rate [FDR] (55). aSyn-induced MRI-derived atrophy was also observed in other regions, such as the thalamus (Figure 1F), amygdala, hypothalamus, hippocampus, in addition to the frontal and a few regions in the parietal cortex. Full volumetric results are provided in Supplementary Table 1 for each structure in the ABA atlas and includes standardised beta-, t-, p- and q-values.

**Figure 1.**
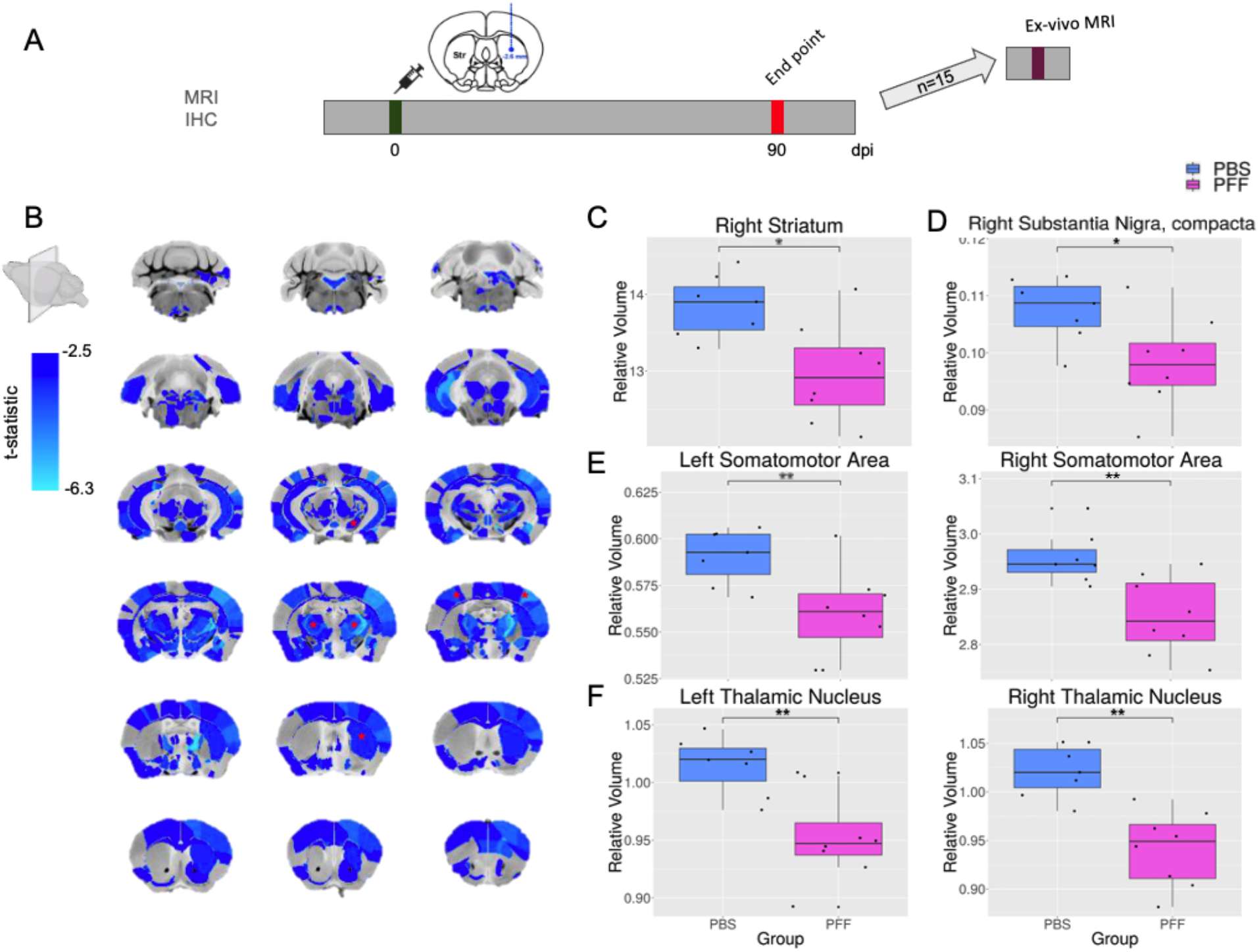
Structure volume differences between PFF- and PBS-injected mice displaying widespread bilateral volumetric decreases in PFF-injected mice. [A] Experimental timeline for mice that underwent *ex vivo* MRI imaging (n=15). [B] Coronal slices of a mouse brain average displayed from posterior to anterior slices with statically significant volumetric differences highlighted. Colourmap denotes t-statistic map of group volumetric effects thresholded at FDR 10% (ranging from the t-value at FDR 10% threshold (bottom; t =-2.48) to the maximum t-value obtained from the linear model (top; t =−6.30)). Red asterisks denote regions of interest plotted below [C-F]. Smaller volumes for the PFF-injected mice were observed for structures in the nigrostriatal pathway such as [C] the injection site (right caudoputamen) (t=−3.45; p=1.22E-13; q=0.0511), [D] right substantia nigra, pars compacta (t=−3.04; p=1.65E-13; q=0.0696), [E] the left and right frontal cortex (the primary motor (left: t=−3.06; p=1.64E-13; q=0.0499; right: t= −3.73; p= 9.03E-14; q=0.0499) and primary somatosensory cortex (left: t=-4.02; p=8.44E-14; q=0.0499; right: t=-5.01; p=1.05E-13; q=0.0499)), [F] the left and right thalamus (e.g. bilateral reticular nucleus) (left: t=-4.22; p=7.24E-14; q=0.0499; right: t= −5.96; p=7.01E-14; q=0.0359).

When examining voxel-wise measures of atrophy using the absolute Jacobian determinants (56), we similarly observed widespread yet subtle atrophy attributable to the PFF injection (surviving FDR 20%). These findings roughly follow our volumetric results and are further detailed in the supplementary materials (section 5.1; Supplementary Figure 2).

#### 2.1.2 Whole brain structural covariance patterns of atrophy

Although our investigation of volumetric differences demonstrated significant volumetric decreases in areas that project to, or receive input from the injection site, we sought to investigate patterns of voxels that preferentially atrophy together. To achieve this, we were inspired by previous evidence of altered volumetric covariance structure that reflects a human PD atrophy pattern (34). Here we used orthogonal projective non-negative matrix factorization (OPNMF) as recently detailed by our group (57–58) and others (59–60). While conceptually similar to other matrix decomposition techniques, such as principal or independent components analysis, this method provides interpretable orthogonal components with positive weights. OPNMF was used to decompose normalised values of voxel-wise volume measures (indexed by Jacobian determinants) into two matrices, a components matrix, which describes groupings of voxels sharing a covariance pattern, and a subject weights matrix, which describes how each subject loads onto each pattern. With the orthogonality constraint, the contribution of each voxel is restricted to only one component, however there may be voxels from the same structure that are present across multiple components.

The number of components, k=4 was chosen by investigating the reconstruction error at different values of k=3-10 (as described in Patel et al. (2020) & Robert et al. (2021) (57–58)). The spatial pattern of voxels for component 1 consists of frontal cortical, posterior thalamic, pons and cerebellar voxels, component 2 consists of ventral cortical and medullary voxels, component 3 consists of mainly subcortical regions such as the anterior striatum and hippocampus, with some brainstem voxels, and component 4 consists of mainly a striatal-nigral-hypothalamic pattern (Figure 2; Supplementary Figure 3).

**Figure 2.**
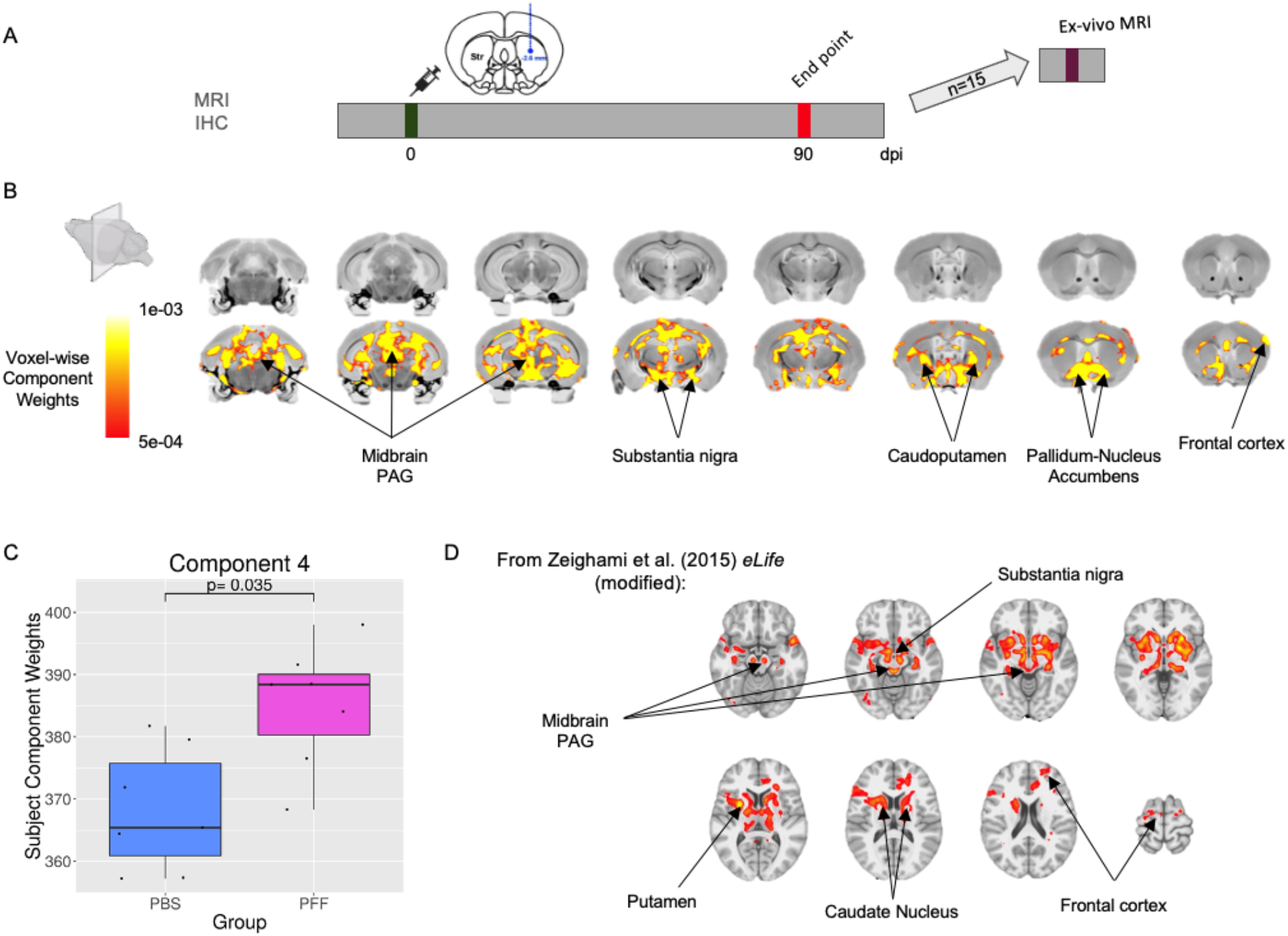
Structural covariance pattern of aSyn-induced voxel-wise atrophy. [A] Experimental timeline for mice that underwent *ex vivo* MRI imaging (n=15). [B] Orthogonal projective nonnegative matrix factorization (OPNMF) decomposition of voxel-based deformation measures in M83 mice that received PFF or PBS injection in the right dorsal striatum. The spatial pattern of voxel scores for component 4 is displayed on the mouse brain average, displaying a network of voxels sharing a similar variance pattern. The colourmap denotes the component weights for each voxel such that higher weights (hotter) represent a stronger covariance with the pattern whereas lower weights (darker red) describe voxels with a lower covariance with the overall pattern. Component 4 voxel-wise weights revealed a striatal-nigral-hypothalamic pattern. [C] Group differences of OPNMF component 4 subject weightings, which describe how each subject loads onto the identified atrophy pattern, were assessed using a general linear model and covarying for sex. Box plot shows significant group effects for component subject weights between the PFF and PBS injected mice, denoting a Hu-PFF specific spatial patterns of atrophy. [D] Distribution of atrophy in humans with Parkinson’s disease, obtained using a similar decomposition technique (34). The eLife article is distributed under the terms of a Creative Commons Attribution License that permits unrestricted use and redistribution provided that the original author and source are credited (creativecommons.org/licenses/by/4.0/). Permission obtained from the first author to use their figure for mouse to human brain atrophy pattern comparisons. Our findings suggest that atrophy patterns from mice replicate human PD atrophy patterns.

A general linear model to examine group differences in subject component weights (with sex in the model) revealed a pattern of aSyn PFF-induced network level atrophy for component 4 (p=0.035). This spatial covariance pattern of component 4 recapitulates a human PD atrophy covariance pattern previously described in Zeighami et al. (2015) (34), using data from humans with PD derived using similar techniques, such as T1w images, DBM, and a multivariate decomposition technique, independent component analysis (Figure 2).

### 2.2 Examining cellular pathology in MRI-derived regions atrophy using immunostaining

To better understand the cellular underpinning of the observed MRI-derived atrophy patterns, we examined four markers in the brain using individual IHC: 1) a proxy for aSyn inclusions (pS129Syn), 2) gliosis (GFAP), 3) neuroinflammation (Iba-1), and 4) dopaminergic cell loss (TH). The regions were selected based on structure-specific volume or network-level local morphological volume differences observed using MAGeT-Brain and OPNMF respectively (n=14 subjects; 28 regions; total of 93 stain-region pairs; Supplementary Table 2). DAB (3,3-Diaminobenzidine)-positive cells (for GFAP, Iba-1, and TH markers) and DAB-positive particles (for pS129Syn marker; given that pS129Syn can be found in the cell body or the axonal and dendritic processes) (normalized by tissue area) was assessed using linear models to examine the injection group by sex interaction for each region-stain pair, and the results were corrected at 5% FDR.

As described in previous studies (61–63), it is well established that misfolded aSyn induces increases in pS129Syn, as well as number of microglia and astrocytes. Similarly, here, we observed significant increases in pS129Syn, Iba-1, and GFAP in PFF-injected compared with PBS-injected mice across various regions of the brain identified in the MRI-derived patterns of atrophy from our network-level analysis; such as the injection site (right striatum), right primary motor cortex, right thalamus, right hypothalamus, left and right nucleus accumbens, left primary motor cortex and left primary somatosensory area periaqueductal gray and medulla left and (Figure 3). Taken together, these findings provide evidence of aSyn-spreading that is associated with atrophy signatures.

**Figure 3.**
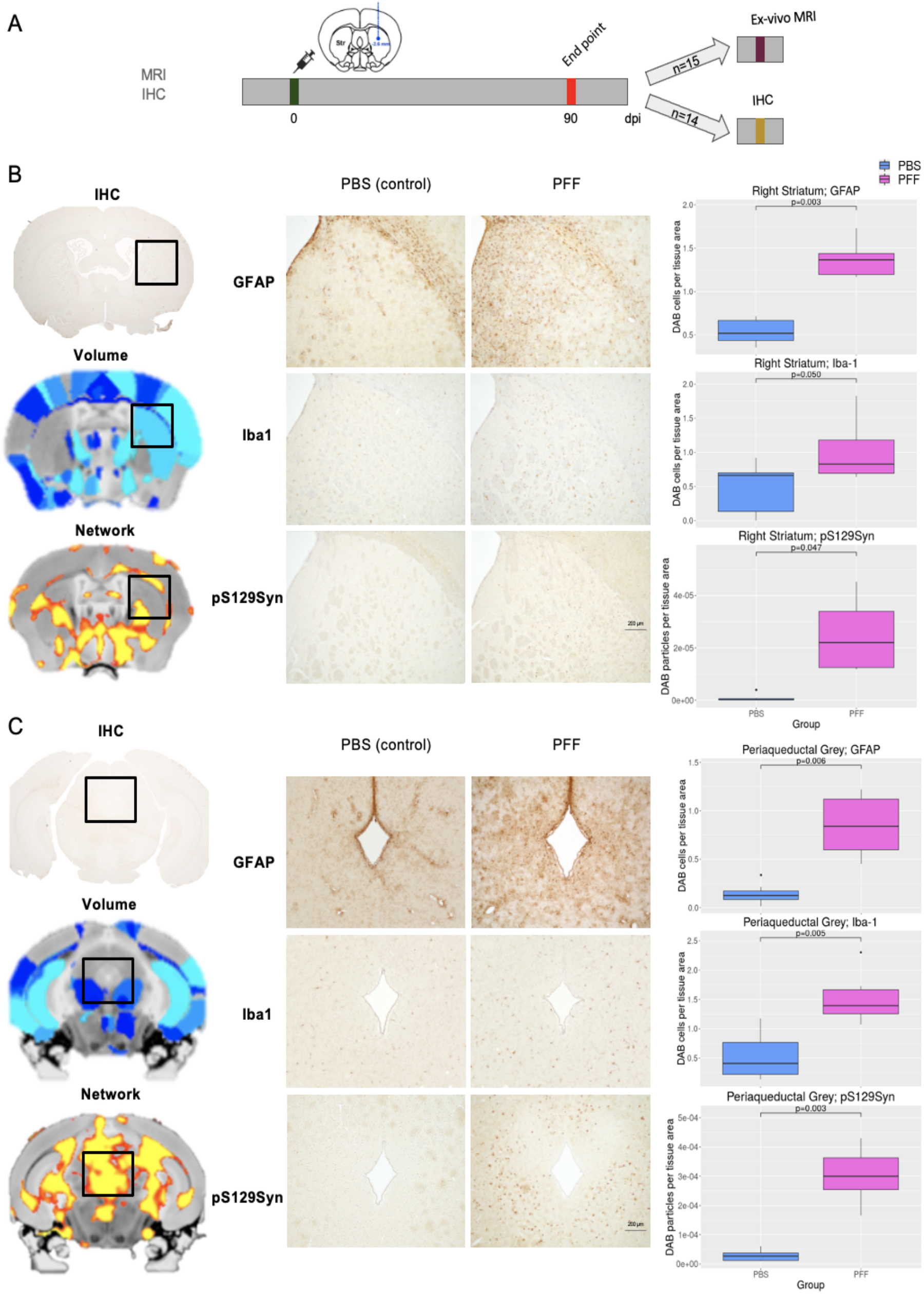
MRI-derived atrophy and IHC cellular underpinnings for M83 PFF-versus PBS-injected mice. [A] Experimental timeline for mice that underwent *ex vivo* MRI imaging (n=15). [B-C] Coronal brain slices *ex vivo* IHC brain (top), MRI brain average slice overlaid with volume (MAGeT) (middle) and network (OPNMF) (bottom) atrophy results shown on the left-hand side. Microscope images of immunostained brain slices zoomed in (10x) onto the region of interest: [B] injection site (right striatum) and [C] periaqueductal gray for each of the three markers (astrocyte marker; GFAP, microglia marker; Iba-1, and aggregate aSyn marker; pS129Syn) for a PBS- and PFF-injected mouse. Group-wise differences in terms of DAB positive markers per tissue area for each marker shown on the right-hand side for both the injection site [B] (GFAP: t=8.21; p=0.003, Iba-1: t=2.52; p=0.05, pS129Syn: t=2.32; p=0.047) and the periaqueductal gray [C] (GFAP: t=5.74; p=0.006, Iba-1: t=4.53; p=0.005, pS129Syn: t=7.41; p=0.003).

Downstream cellular players involved in atrophy, such as astrocytes and microglia were more abundant in regions closely connected to the injection site and part of our observed network for the PFF-injected mice. The regions include the right substantia nigra and right midbrain reticular nucleus, as well as an increase in astrocytes in the right primary somatosensory cortex, left substantia nigra and left striatum; regions that are a part of the nigrostriatal pathway.

### 2.3 Assessing pre-symptomatic cognitive and motor deficits

Given the brain regions affected by atrophy in PFF-injected mice (cortical-striatal and thalamic networks), we anticipated that executive function, and in particular behavioural flexibility and reversal learning (64–66), could provide a cognitive correlate that reflects aSyn misfolding and spreading. Indeed, recent work suggest that aSyn toxicity in M83 homozygous mice triggers reversal learning, but not attentional deficits (67). Since we observed no sex differences in our imaging findings, we chose to focus this experiment on male mice, given the predominance of PD in men and because of the higher statistical power needed to study sex as a biological variable in behaviour performance (45, 67). We tested mice longitudinally (49±7 and 84±7 dpi) using pairwise visual discrimination and reversal learning (PVD-R, 45) in two independent cohorts of PFF- or PBS-injected M83 hemizygous mice (Figure 4A-K) using two different sets of images; one set in which the images are more easily differentiable (Figure 4F-H) and a second set in which the images are harder to differentiate (Figure 4I-K). These same mice were also tested in a series of motor tasks (open field, wire hang, grip force, and rotarod) following completion of the touchscreen tests (at 63±7 and 98±7 dpi; the motor task data from the two cohorts were pooled as there was no differences between cohorts, Figure 5A).

**Figure 4.**
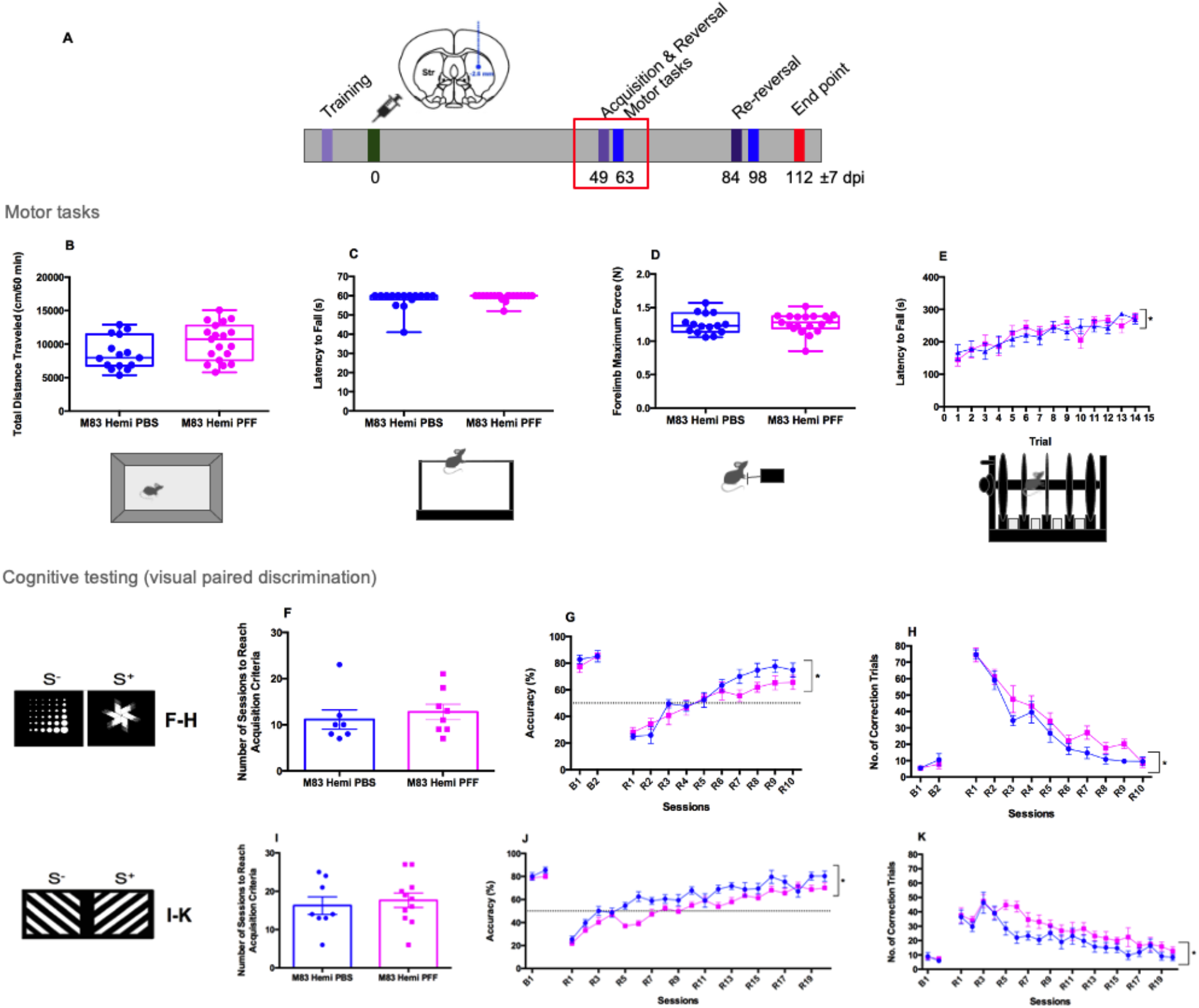
Cognitive deficits appear before motor deficits in the M83 +/-PFF model. [A] Experimental design comparing M83+/-(Hemi) injected with PBS or PFFs (wpi, weeks post injection); [B] Total distance (cm) traveled in the open field during 60 minutes (Unpaired t test: t=1.865 df=32 p=0.0714); [C] Latency to fall (s) in the wire-hang test (Mann Whitney non-parametric test: p= 0.4387); [D] Grip force (N) (Unpaired t test: t=0.1657 df=32 p= 0.8695); [E] Latency to fall (s) in the rotarod test (Repeated measures two-way ANOVA: Session: F (13, 416) = 8.118, p < 0.0001; Genotype: F (1, 32) = 0.02405 p = 0.8777; Interaction: F (13, 416) = 0.8244, p = 0.6342) of PBS-injected M83 mice and PFFs-injected M83 mice (N=15 for PBS and 19 for PFFs using two independent PFF preparations). [F] Number of sessions to reach acquisition criteria in the PVD test using Marble (S-) and Fan (S+) stimuli during the acquisition phase (Unpaired t test: t=0.6051 df=13 p=0.5555); [G] Mice were baselined for 2 days and reversal learning tested Accuracy (%) (Repeated measures two-way ANOVA: Session: F (5.531, 69.89) = 35.24, p < 0.0001; Genotype: F (1, 13) = 2.385 p = 0.1465; Interaction: F (11, 139) = 1.335, p = 0.2115); [H] Number of correction trials performed (Repeated measures two-way ANOVA: Session: F (4.521, 57.54) = 56.17, p < 0.0001; Genotype: F (1, 13) = 4.789 p = 0.0475; Interaction: F (11, 140) = 0.8458, p = 0.5948). PBS-injected M83 mice (N=7) and PFFs-injected M83 mice (N=8). Parameters were measured across baseline days 1 and 2 (B1, B2) and reversal days 1 to 10 (R1-R10). [I] Number of sessions to reach acquisition criteria in the PVD test using diagonal lines facing down (S-) and diagonal lines facing up (S+) stimuli during the acquisition phase (Unpaired t test: t=0.4690 df=17 p=0.6450); [J] Mice were baselined for 2 days and reversal learning tested Accuracy (%)(Repeated measures two-way ANOVA: Session: F (7.549, 128.3) = 40.36 p < 0.0001; Genotype: F (1, 17) = 9.098 p = 0.0078; Interaction: F (21, 357) = 1.881 p =0.0114); [K] Number of correction trials performed(Repeated measures two-way ANOVA: Session: F (5.824, 99.01) = 18.14 p < 0.0001; Genotype: F (1, 17) = 4.178 p = 0.0568; Interaction: F (21,357) = 1.231 p = 0.2212). PBS-injected M83 mice (N=8) and PFFs-injected M83 mice (N=11). Parameters were measured across baseline days 1 and 2 (B1, B2) and reversal days 1 to 20 (R1-R20). Results are expressed as mean ± SEM, Unpaired t test (B,D,F,I); Mann Whitney test (C) and Repeated measures two-way ANOVA (E,G,H,J,K). Asterisks indicate statistical differences between groups: *p<0.05.

**Figure 5.**
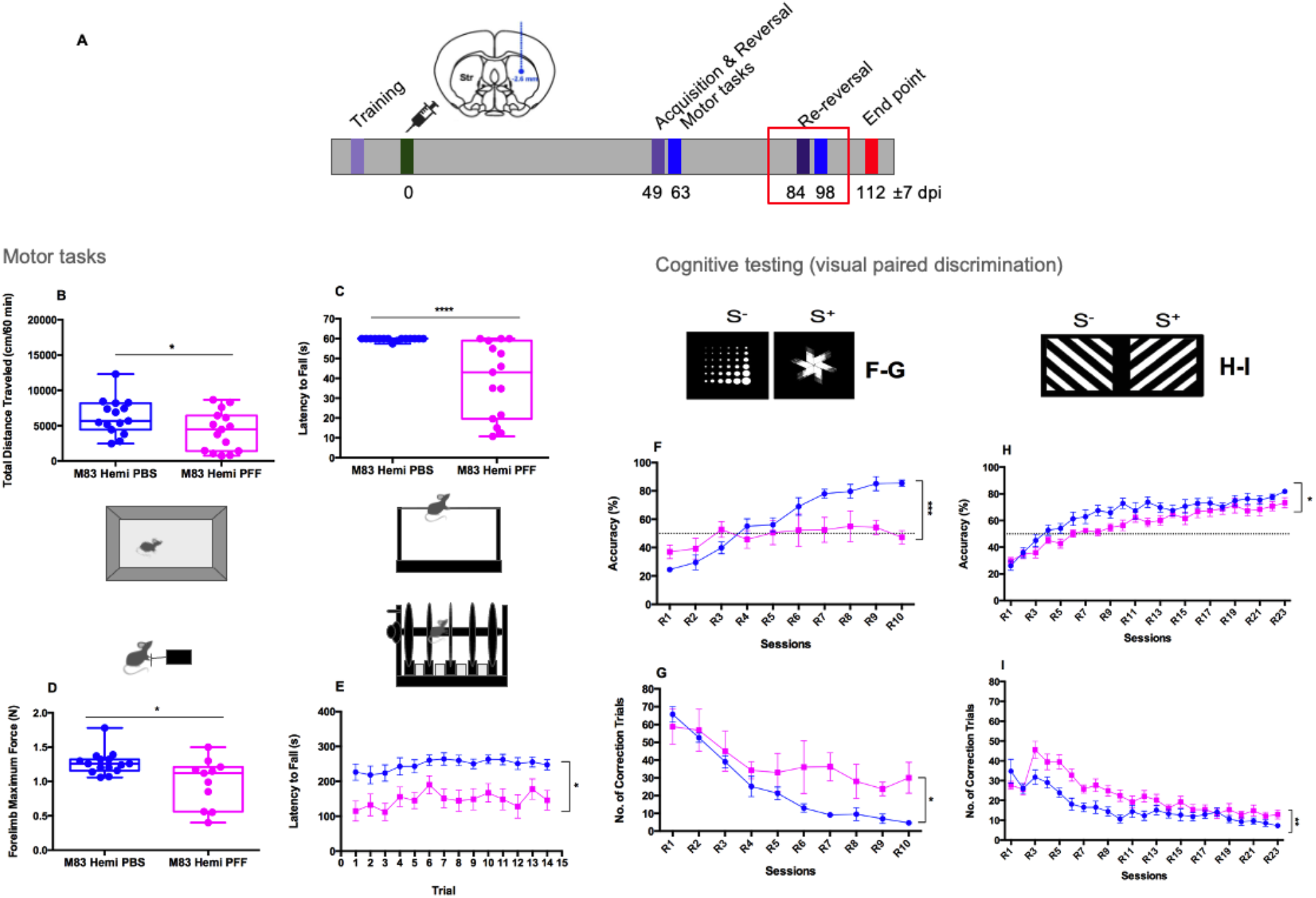
Progressive cognitive and motor deficits in the M83 +/-PFF model. [A] Experimental design comparing M83+/-(Hemi) injected with PBS or PFFs (wpi, weeks post injection); [B] Total distance (cm) traveled in the open field during 60 minutes (Unpaired t test: t=2.083 df=28 p=0.0465); [C] Latency to fall (s) in the wire-hang test (Mann Whitney non-parametric test: p < 0.0001); [D] Grip force (N) (Mann Whitney non-parametric test: p = 0.0211); [E] Latency to fall (s) in the rotarod test (Repeated measures two-way ANOVA: Session: F (6.124, 167.2) = 1.569, p = 0.1577; Genotype: F (1,28) = 17.97 p = 0.0002; Interaction: F (13, 355) = 0.3815, p = 0.9752) of PBS-injected M83 mice and PFFs-injected M83 mice (N=15 for PBS and 15 for PFFs using two independent PFF preparations. Of note, 4 PFFs-injected mice succumbed from the disease and a total of 11 PFFs mice performed the grip force task). [F] Re-reversal performance using marble and fan stimuli Accuracy (%) (Repeated measures two-way ANOVA: Session: F (3.105, 27.95) = 14.23, p < 0.0001; Genotype: F (1, 9) = 6.970, p = 0.0269; Interaction: F (9, 81) = 6.418, p < 0.0001); [G] Number of correction trials (Repeated measures two-way ANOVA: Session: F (2.893, 26.04) = 22.05, p < 0.0001; Genotype: F (1,9) = 6.419, p = 0.0320; Interaction: F (9, 81) = 2.412, p = 0.0177) performed by PBS-injected M83 mice (n=7) and PFFs-injected M83 mice (n=4, 4 mice had become symptomatic and had to be collected prior to the test) in the re-reversal test. Parameters were measured across re-reversal days 1 to 10 (R1-R10). [H] Re-reversal for PVD-R Accuracy (%) (Repeated measures two-way ANOVA: Session: F (6.632, 111.2) = 29.67, p < 0.0001; Genotype: F (1, 17) = 7.365, p = 0.0147; Interaction: F (22, 369) = 1.068, p = 0.3805); I) Number of correction trials (Repeated measures two-way ANOVA: Session: F (6.367, 106.5) = 19.60, p < 0.0001; Genotype: F (1, 17) = 12.34, p = 0.0027; Interaction: F (22, 368) = 2.227, p = 0.0014) performed by PBS-injected M83 mice (n=8) and PFFs-injected M83 mice (n=11). Diagonal lines facing down (S-) and diagonal lines facing up (S+) stimuli were used during the PVD-R re-reversal phase. Parameters were measured across re-reversal days 1 to 23 (R1-R23). Results are expressed as mean ± SEM, Unpaired t test (B); Mann Whitney test (C,D) and Repeated measures two-way ANOVA (E,F,G,H,I). Asterisks indicate statistical differences between groups: *p<0.05.

At the early time point (around 60 dpi), we found no differences in motor performance between PBS and PFF-injected M83 hemizygous mice in locomotor activity (Figure 4B), wirehang test (Figure 4C), grip force (Figure 4D) or rotarod (Figure 4E), suggesting that at this time point motor behaviour was unimpaired. Body weight and vertical activity were also not changed (Supplementary Figure 4).

On the touchscreen tests, at the 49 dpi time point, PBS and PFF-injected M83 mice performed identically in their ability to recognize the easy set of images (Figure 4F). In contrast, when we reversed the contingency with this easy set of images, PFF-injected mice performed significantly worse than PBS-injected mice (Figure 4G), had increased correction errors (Figure 4H), showed slightly delayed reward collection latency, and slightly lower number of trials completed. The deficits in reversal learning appeared mainly at the last stages of the task, suggesting the mice made more errors in learning the new contingency. An independent cohort of mice tested with the more difficult images showed similar deficits, increasing errors after the initial 3-day reversal and in the later phases of reversal learning (Figure 4I-K). Reward correct latencies and trials completed were also altered in PFF-injected mice compared to controls (Supplemental Figure 5). These results suggest that PFF-injected mice present a selective deficit in reversal learning. M83 mice injected with PBS performed identically to control wild-type mice (not shown).

We repeated the reversal learning (re-reversal, Fan stimulus rewarded) at 84±7 dpi, and again investigated motor performance in mice after the touchscreen testing (98±7 dpi). PFF-injected mice presented profound motor deficits at this age, congruent with the starting of symptomatic disease phase (Figure 5A-E). During re-reversal mice demonstrated a strong deficit in reversal learning compared to PBS-injected mice (Figure 5F, G). Re-reversal using the more difficult set of images also revealed a behavioural deficit, with decreased accuracy and increased correction errors (Figure 5H-I). The results with the more difficult set of images, in which mice take more than twice as much to reach criteria for proper acquisition of the task, suggest again a deficit in relearning the initial contingency for PFF-injected mice during reversal. Moreover, correct touch latency and reward collection latency were also affected in PFF-injected mice (Supplementary Figure 5), likely reflecting motor deficits that become apparent at this stage. Immunofluorescence analysis confirmed increased aSyn S129 phosphorylation and increased microglial staining in these mice in brain regions whereby we observed MRI-derived atrophy (Supplemental Figure 7-8).

## 3. Discussion

Using translational methodologies (MRI and cognitive touchscreen technology), our results provide evidence for the prion-like spreading of aSyn from a disease epicenter (the injection site: right dorsal striatum), resulting in brain atrophy, cellular pathology, and behavioural deficits. We observed widespread effects on brain morphology at 90 days post-injection in PFF-injected M83 hemizygous mice in bilateral regions that project to or receive input from the injection site, highlighting a network of regions implicated in PD-related behavioural deficits. While high-level cognition (reversal learning) and motor impairment coincided with the emergence of these atrophy patterns, there is significant evidence that cognitive impairments emerge earlier. This suggests that the toxic spreading of misfolded aSyn from the dorsal striatum to connected regions preferentially affects high-level cognition potentially arising from the patterns of cortical-striatal-thalamic atrophy we observed. The pattern that we observed using MRI data, a methodology translatable across species, strongly suggests that the observed pattern of PFF-induced atrophy recapitulates patterns observed in human patients with PD (34). Similarly, the behavioural testing we used examines cognitive domains that are affected in PD and other synucleinopathies (68–71). Accordingly, there were global increases in pS129Syn, microglia, and astrocytes staining in PFF-injected compared with saline-injected mice, indicating that PFFs induce alpha-synuclein aggregate formation, neuroinflammation, and gliosis underlying MRI-observable pathology.

### 3.1 The case for minimally invasive translational methodologies

The work described in this manuscript further validates the M83-Hu-PFF mouse model of synucleinopathy-associated pathogenesis of neurodegenerative diseases by demonstrating the utility of state-of-the-art methodologies to examine endophenotypes in the mice that replicate those in human patients with the disease. Here, we identified biomarkers that are rapid and translational that align with traditional cellular correlates of pathology, with regards to changes in anatomy and cognitive functioning. Specifically, we show that neuroanatomical pathology can be detected distal to the inoculation site prior to onset of motor impairments and coincides with cognitive deficits observable using touchscreens. These findings are suggestive of a set of biomarkers that can serve as a proxy for the direct examination of cellular levels for pathology. In fact, we anticipate that these biomarkers can measure progression of toxicity, specifically in the early phases of protein misfolding and that they can be more reliable than end stage pathology.

The signatures of atrophy detected using our MRI-based measures further demonstrate that regions that are interconnected with the injection site displayed the most severe pathology. For example, given the significant connectivity between basal ganglia regions and the thalamus, there is a reasonable expectation of aSyn propagation resulting in downstream atrophy. Further, the atrophy of commissural tracts such as the corpus callosum strongly suggests wide-spread propagation across hemispheres (72). Our observations of the increased presence of phosphorylated aSyn, astrocytes, and microglia coincident with atrophy covariation due to PFF injection suggest that atrophy patterns and downstream inflammation are primary drivers of the MRI-observable patterns. Specifically, our data suggest that the effects of pathology (in terms of MRI-derived atrophy) may be more pronounced for downstream players of said atrophy, specifically in terms of microglia and astrocyte infiltration, as opposed to the direct effect of pathological aSyn deposition in regions known to be connected to the PFF injection site. Interestingly, the increased presence of astrocyte and/or microglia markers in these regions were not accompanied by deposition in phosphorylated aSyn in the PFF group. These findings further suggest an infiltration of atrophy- and inflammation-mediating cells prior to severe pS129Syn deposition. We did not observe any significant differences in TH between the two injection groups, suggesting minimal impact of the PFF spreading on the dopaminergic system; however, dopaminergic cell loss is variably seen across the literature in this model and if so, is commonly observed in later stages of the disease progression (>90 days post-injection) (21,24). These data add important information that supports the previously reported findings describing pathophysiology reminiscent of Lewy body inclusions in regions proximal to the injection site of the PFFs (20,21,73,22–28). The homology between our MRI findings and the atrophy patterns observed in clinical PD populations described in Zeighami et al. (2015) (34) further support the translational relevance of this model. Taken together, our results call for further investigation of the longitudinal and progressive nature of the prion-like spreading of aSyn *in vivo* as a means of achieving careful characterization of a PD-like pathophysiology that can be used to test novel therapeutic interventions without having to only rely on end stage pathology.

In addition to the translatability of our MRI-derived atrophy findings, our motor and cognitive symptomatology results may have translational relevance in the context of human synucleinopathies, such as PD. Several studies examining motor performance in PFF-injected mice using traditional motor paradigms have described impaired performance on the rotarod test and other traditional behaviours (22,24,25). Whether some component of this dysfunction relates to accumulation of synuclein in the spinal cord is not fully understood. Moreover, these deficits are not consistently observed. Some studies report no significant differences (25) or even observe improved performance post-injection (74–75). Here, we only observe significant decline in motor performance for PFF-injected mice at 98 ±7 dpi (past the 90 dpi MRI timepoint). Taken together, these findings suggest the need to develop tests that are more sensitive than motor impairments to early toxicity of proteinopathies.

Typically, in the prodromal PD phase, the spread of protein aggregates resulting in the appearance of motor symptomatology manifests only after extensive pathology (3). Accordingly, non-motor symptoms have been extensively reported among patients later diagnosed with PD (76–78), appearing long before the characteristic motor impairments (79), thus pointing to early biomarker potential of non-motor phenotypes. While prodromal symptoms in patients with PD are most commonly autonomic (constipation, loss of sense of smell, etc …) and sleep disturbances, these symptoms are difficult to examine in mice. Furthermore, the prodromal period is significantly shorter in these mice compared to humans, likely due to the nucleation phase, which is skipped with the injection of the fibrils, thereby dramatically accelerating the disease process. Thus, deficits beyond motor impairment are rarely examined, other than the occasional use of cognitive paradigms such as the Y-maze test, Morris Water maze or object recognition (80,22,24). An alternate mouse model of PD (6-OHDA injected C57BL/6N mice) was tested in touchscreens and found to have attention deficits, but not reversal learning deficits (81). Conversely, with the M83 model, recent experiments by our group suggest that toxic aSyn in homozygous M83 mice (not injected with PFFs, which spontaneous develop disease around a year of age) leads also to reversal learning deficits but not attention deficits (67). Moreover, in this work, reducing the load of aSyn misfolding (with the introduction of a chaperone transgene) mitigated these behavioural deficits and resulted in milder patterns of MRI-derived atrophy in the brain. Although these experiments (in M83 homozygous mice) lack the timed initiation of conversion and prion-like spreading of misfolded aSyn to connected networks, in which can be captured in the PFF-injected M83 hemizygous model, taken together with our current findings, these data suggest that spreading of toxic aSyn misfolding is critical to cause atrophy and cognitive deficits. Mechanistically, recent experiments have suggested a role for thalamic projecting globus pallidus parvalbumin-expressing neurons in reversal learning associated with PD (82). Our results support the notion that global volumetric changes observed in cortico-striatal-thalamic networks after spreading of toxicity may contribute to the cognitive deficits. Impairment in early phases of reversal learning is usually thought to reflect behavioural flexibility and are associated with cortical-striatal dysfunctions (83). In contrast, late deficits in reversal learning may be associated with basal ganglia dysfunction, and some of recent work suggest that dysfunctional dopamine-acetylcholine balance affects reversal learning but shifting goal-directed to habitual behaviour (66). Future studies using more direct tests of striatal function may provide even more sensitive cognitive biomarkers to follow the spreading and toxicity of misfolded aSyn.

### 3.2 Limitations of the study

Although the use of PFF injection to reproduce spreading of synucleinoipathies is widespread, it likely does not reproduce the true spreading observed in human synucleinpathies, which do not start in the striatum (3,84). Hence, although our experiments show a pattern of brain atrophy and cognitive deficits that resemble PD changes in humans, they likely reflect toxicity of aSyn spreading from connected regions after striatal injection of PFFs. While small animal MRI imaging is minimally invasive, whereby we can acquire *in vivo* images of the brains of anaesthetised mice, and track brain anatomy changes over time (an essential part of our future studies), here we acquired *ex vivo* images of the mouse brains due to nature of the multi-site experimental design. Nonetheless, our data speak to the utility of this translational tool. Despite the limitation of cross-sectional data, multivariate techniques such as NMF (59) used here allow for the investigation of whole brain-wide aSyn-induced atrophy patterns. While this technique does not model brain change, it does support a network-like spreading model of neurodegeneration. Nonetheless, we infer connectivity based on covariance structure in our MRI data. However, the addition of retro- and antero-grade tracers, diffusion-weighted imaging, and resting-state functional MRI would complement connectivity studies and would be greatly beneficial as these data provide direct or complimentary information about brain connections. Another important limitation of the current exploratory study is the focus on male mice for cognitive testing, despite the observation that both male and female mice present similar changes in brain volume in critical brain regions. Future experiments should investigate whether female mice present similar behavioural deficits.

### 3.3 Conclusions

It is remarkable that brain areas that show consistent atrophy also contribute to high-level cognitive deficits observed in this study. Our experiments also suggest that cognitive effects appear to arise earlier than motor deficits in this model, which to the best of our knowledge has been tested for the first time using touchscreen tasks in the mouse PFF model. The work presented here provides a detailed validation of the PFF model for the examination of synucleinopathy-associated pathogenesis. We find that changes in MRI imaging biomarkers and high-level cognition resemble those involved in human synucleinopathies, such as PD. This suggests that the combination of PFF injection to spread toxicity and aSyn pathology combined with the use of these two early biomarkers of toxicity may be more sensitive and afford more faithful translation for testing new disease-modifying compounds in PD and other synucleinopathies.

## 4. Material and Methods

### 4.1 Animals

We used transgenic hemizygous M83 mice, expressing one copy of human alpha-synuclein bearing the familial PD-related A53T mutation under the control of the mouse prion protein promoter (85), in addition to the endogenous mouse alpha-synuclein, maintained on a C57BL/C3H background. The M83 hemizygous mice used here present no phenotype until they are 22 and 28 months of age, on average (85), unless aSyn PFFs are injected to trigger accumulation of misfolded toxic aSyn.

#### 4.1.1 MRI and IHC experiments

Hemizygous TgM83+/-mice (B6; C3H-Tg[SNCA]83Vle/J) were purchased from The Jackson Laboratory (004479, Bar Harbor, ME). All mice were housed at the Montreal Neurological Institute’s (MNI) Animal Facility (McGill University, Montreal, QC, Canada) under standard housing conditions with food and water *ad libitum*. Mice were originally housed in groups of four upon arrival from JAX, however due to fighting occurring between males, all mice were individually housed at the start of the experiment to prevent further fighting. Mice were housed with a 12/12-hour light/dark cycle, with lights on at 08:00. All study procedures were performed in accordance with the Canadian Council on Animal Care and approved by the McGill University Animal Care Committee.

The experimental timeline with all the procedures performed at each time point is depicted in Figure 1. Prior to any intervention, the mice underwent weighing prior to injection time point and prior to the experimental endpoint.

#### 4.1.2 Behaviour and IF experiments

All procedures followed guidelines of Canadian Council for animal care and were approved by the local animal care committee at the University of Western Ontario (Protocol number 2020-163) Hemizygous TgM83+/-mice (B6; C3H-Tg[SNCA]83Vle/J) were purchased from The Jackson Laboratory (004479, Bar Harbor, ME) and bred at the University of Western Ontario so littermate control mice (M83+/-injected with PBS or wild-type mice) could be used in the experiments.

All mice that underwent behavioural tasks were single housed (because of aggressiveness of the strain) in a temperature and pressure-controlled room with a 12:12 light-dark cycles (7am:7pm) at the University of Western Ontario. The room was maintained at 22-24°C and 40%-60% of air humidity level. Environmental enrichment was not provided to the mice and cages were changed weekly. All tasks in the touchscreen battery are motivated by strawberry milkshake (Nielson-Saputo Dairy Products) reward. Thus, to ensure adequate motivation to work for food rewards, mice (10-12 weeks or older) were food-restricted at least two weeks prior to the start of behavioural testing and were maintained at 85% of free feeding body weight until they were euthanized. All mice were weighed, and food pellets ranging from 1.5-3 grams (3.35kcal/gram) were delivered to animals upon return to respective home cage after daily testing. Food pellets are commercially available at Bio-Serv in Flemington, New Jersey. Drinking water was provided ad libitum. All behavioural tests were conducted during the light phase.

#### 4.1.3 Injection material and Stereotaxic injections

To investigate toxic aggregated alpha-synuclein spreading, we injected healthy 3-4-month-old hemizygous M83 mice with preformed fibrils (PFFs) of human alpha-synuclein or phosphate-buffered saline (PBS) in the right dorsal striatum to have a known disease epicentre from which to map the spreading of alpha-synuclein and the consequences for cognition and motor behaviour. Injections were done according to published protocols for PFF generation and preparation (23). The PFFs were made and characterized in-house by the Early Drug Discovery Unit (EDDU) according to established SOPs (production: https://zenodo.org/record/3738335#.Yp46fXbMKUl; characterization: https://zenodo.org/record/3738340#.YzIjBezML0p) at the Montreal Neurological Institute following the protocol described in Volpicelli-Daley et al. (2014). PFFs were sonicated and DLS analysis was completed to ensure the average diameter of PFFs was <100nm. PFFs were added to 200 mesh copper carbon grid (3520C-FA, SPI Supplies), fixed with 4% PFA for 1minute and stained with 2% acetate uranyl (22400-2, EMS) for 1 minute. PFFs were characterized using a negative staining protocol (https://zenodo.org/record/3738340#.Yp46xnbMKUl). PFFs were visualized using a transmission electron microscope (Tecnai G2 Spirit Twin 120 kV TEM) coupled to a AMT XR80C CCD Camera, and analyzed with Fiji-ImageJ 1.5 and GraphPad Prism 9 software. Electron microscope characterization of the fibrils in terms of distribution per length can be seen in Supplementary Figure 1.

Briefly, the mice were anaesthetized with isoflurane (5% induction, 2% maintenance), given an injection of carprofen (0.1cc/10 grams) and 250 mg/ml bupivacaine subcutaneously for pain relief (for the mice part of the imaging experiments) or Metacam (10mg/kg for the mice part of the behavioural experiments), and positioned into a stereotaxic platform. Recombinant human alpha-synuclein fibrils (total volume: 2.5 μL; 5 mg/mL; total protein concentration, 12.5 μg per brain) were stereotaxically injected in the right dorsal striatum (co-ordinates: +0.2 mm to +0.3 mm relative to Bregma, +2.0 mm from midline, +2.6 mm beneath the dura) (Figure 1). The control condition was M83 mice receiving sterile PBS in the same injection site. Injections were performed using a 5 μL syringe (Hamilton; 33 gauge) at a rate of 0.25 μL per min with the needle in place for about 5 min prior to and after infusion of the inoculum. Different syringes were used for each type of inoculum to prevent any contamination. Post-injection, the mice were placed on a heating pad for recovery before being returned to their home cage. Two groups of mice were injected: one for the imaging experiment and the other for the behavioural experiment. A total of 8 mice per injection group per sex were injected, however after surgery, the final numbers were 6 PBS-injected males, 8 PBS-injected females, 8 PFF-injected males and 7 PFF-injected females, for a total of 29 mice for the MRI and IHC analyses. For behaviour, two separate sets of male mice (n=34) were injected, with two independent PFF preparations. All mice performed the Pairwise visual discrimination task with reversal (PVD-R) to evaluate reversal learning using the touchscreen technology at approximately 49 and 84 ±7 days after PFF injection. Motor tasks including open field, wire hang, grip-force and rota-rod were also performed after each of the touchscreen tests. The first cohort with 7 PBS-injected mice and 8 PFF-injected mice underwent PVD-R with the easy set of images, Marble and Fan. The second cohort with 8 PBS-injected mice and 11 PFF-injected mice was used to confirm the data with a more difficult set of images, diagonal lines (sloping upward or downward).

For one set of mice, after inoculation, the mice were monitored weekly for health and neurological signs such as reduced grooming, kyphosis, and/or decreased motor functioning (including reduced ambulation, tail rigidity, paraparesis), whereby the frequency of monitoring increased upon the onset of the symptomatology up until the predefined endpoint time point. At 90 dpi, the mice were perfused, and the brains were prepped for either *ex-vivo* MRI or immunohistochemistry to examine the spreading properties of aSyn in the earliest stages of the disease.

*A* second set of mice were tested for behaviour, starting at 49±7 dpi and then again at 84±7 dpi. These mice were followed longer and were typically perfused for immunofluorescence analysis between 98-112±7 dpi. Two mice injected with PFFs in the first cohort (easy set of images) did not present robust pathology and therefore were removed from the analysis.

### 4.2 Magnetic Resonance Imaging (MRI) acquisition

Prior to *ex vivo* MRI, 15 mice (~7-8 mice per injection group; ~3-4 per sex) were anaesthetised with isoflurane and transcardially perfused with 0.9% (wt/vol) PBS with 0.4% ProHance (gadoteridol, a gadolinium-based MRI contrast agent and 0.1% heparin (an anticoagulant), followed by 10% formalin withT 0.4% ProHance. Prior to scanning, the brains were stored in sodium azide 0.1 M for about 3-4 weeks, until the brains were scanned. These steps are in accordance with recommendations detailed by Cahill et al. (2012) (86) to prevent fixation artifacts and improve the quality of the MR images acquired.

MRI acquisition was performed on 7.0-T Bruker Biospec (70/30 USR) 30-cm inner bore diameter; AVANCE electronics) at the Douglas Research Centre (Montreal, QC, Canada). High-resolution *ex vivo* T1-weighted images (FLASH; Fast Low Angle SHot) were acquired for each subject (TR=21.5 ms, TE=5.1ms, 70 μm isotropic voxels, 2 averages, scan time= 23:50 min, matrix size=260×158×210, flip angle=20°). Brains were scanned in-skull to improve the accuracy of automated image registration methods (84) for improved MRI-derived measures.

#### 4.2.1 Image processing

##### 4.2.1.1 Pre-processing

All brain images were exported as DICOM from the scanner and converted to the MINC (Medical Imaging NetCDF) file format. Image processing was performed using the MINC suite of software tools (http://bic-mni.github.io). Next, the images were stripped of their native coordinate system, left-right flipped to compensate for Bruker’s native radiological coordinate system, denoised using patch-based adaptive non-local means algorithm (87), and affinely registered to an average mouse template (the Dorr–Steadman–Ullman atlas; 88-90) to produce a rough brain mask. Next, a bias field correction was performed, and intensity inhomogeneity was corrected using N4ITK (91) at a minimum distance of 5 mm. MRI workflow post-pre-processing described in Figure 6. Pre-processed MRI data is publicly available (doi: 10.5281/zenodo.7186777).

**Figure 6.**
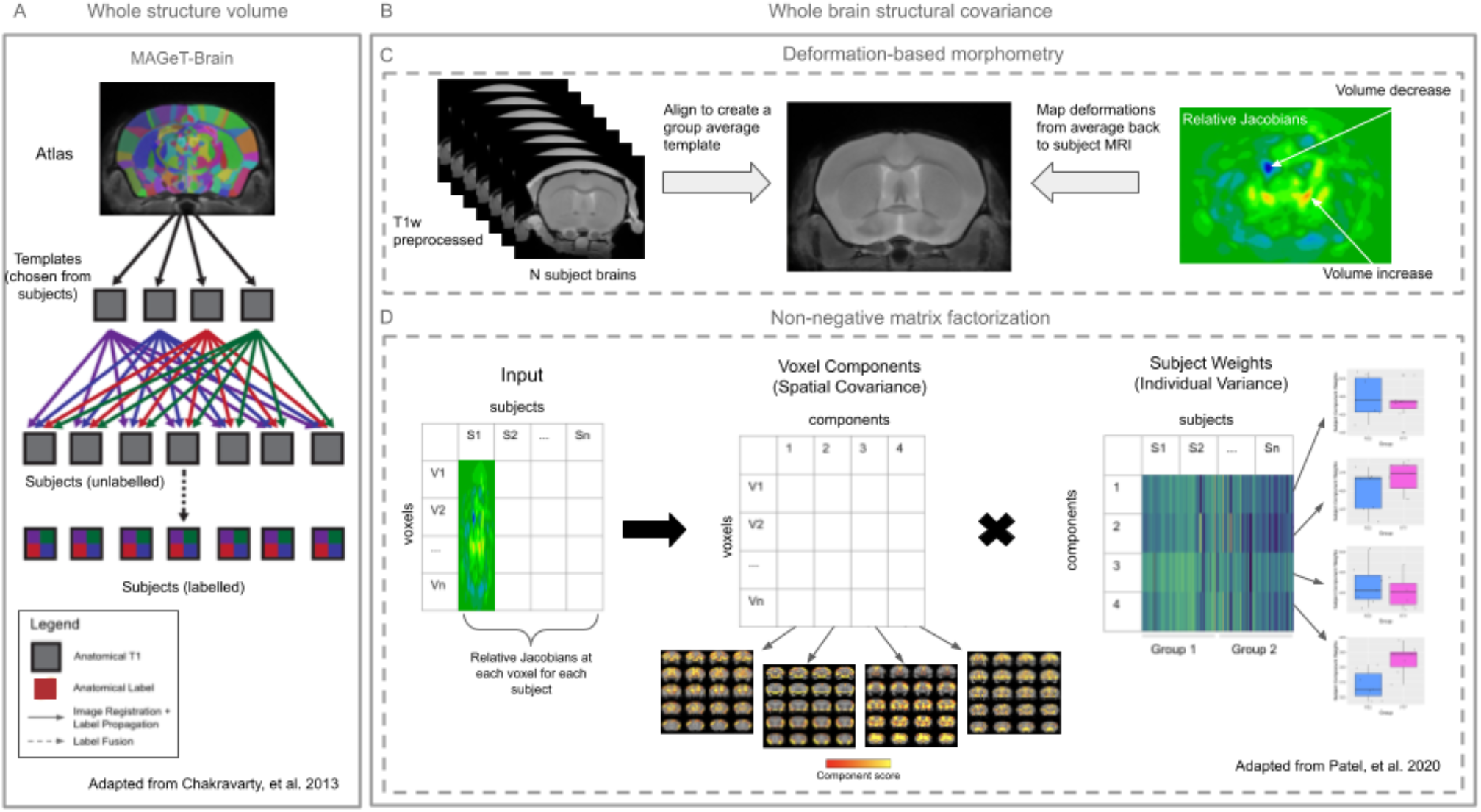
MRI Methods Workflow for the volumetric and network-like analysis. Two analyses were performed using the MRI data: [A] whole structure volumes were examined using the MAGeT-Brain segmentation and [B] whole brain structural covariance (network-like) analysis was performed using two techniques; [C] deformation-based morphometry and [D] non-negative matrix factorization. [A] MAGeT-Brain segmentation pipeline: a priori definitions of structure MRI delineations, provided by a downsampled version of the Allen mouse brain atlas, are propagated onto subject MRI images using image registration techniques (nonlinear registrations). Templates are a subset of the subject images whereby the atlas is first propagated onto these subjects as a way of artificially inflating the number of atlases. [C] Deformation-based morphometry (DBM): Voxel-wise measures of volume (Jacobian determinants) are generated by registering all subject images, creating a group average, such that by mapping each subject image from the average back to its individual space, we obtain information about voxel increase and decreases. [D] The voxel-wise Jacobian values obtained from DBM for each subject were used as input (voxels by subject) and decomposed using orthogonal projective non-negative matrix factorization (OPNMF) into two matrices; 1) voxel components matrix, which describes clusters of voxels sharing a spatial covariance pattern (voxels by components), such that for each component, a voxel’s weight describes how strongly that voxel’s volume covaries with every other voxel in the brain, and 2) subject weights matrix, which describes how each subject loads onto each pattern (components by subjects), whereby for each component, group difference in subject weights (how strongly each subject relates to a spatial covariance pattern) can be assessed using general linear models.

##### 4.2.1.2 MAGeT volumetric segmentation

Before examining voxel-wise covariance patterns of atrophy using OPNMF, we examined structure volume differences between the two injection groups to assess the effect of PFF on global neuroanatomy of the brain. To obtain structure volumes, the pre-processed MRI images were segmented into 355 structures per hemisphere (with 53 interhemispheric regions), for a total of 763 regions using the Multiple Automatically Generated Templates (MAGeT) Brain segmentation algorithm (48), and a down-sampled version of the Allen Mouse Brain Atlas (49). Given the difference between the resolution of the Allen Institute Mouse Brain Atlas (49) and the MRI data acquired, a hierarchically grouped version of the Allen Institute Mouse Brain Atlas was used where select regions were hierarchically grouped into coarser-grained labels due to such limitations of the MRI image resolution and contrast. The segmentations were first propagated from the atlas to a subset of the subject data, known as the template library. From there, the segmentations from the template library were propagated to each subject image via a pairwise nonlinear image registration with a version of Automatic Normalization Tools (ANTS) compatible with the minc-toolkit (https://github.com/vfonov/mincANTS). This step yields multiple segmentations per subject, which were then fused using majority-vote label fusion to generate a final segmentation (92). From these final segmentations, volumes of brain structures were computed and analysed for group differences.

##### 4.2.1.3 Deformation-based morphometry

The denoised and inhomogeneity-corrected images were used as input for the model building image registration tool component of the Pydpiper toolkit (54). In order to obtain a voxelwise measure of neuroanatomy, mice brains were registered together through a series of linear and nonlinear registration steps with iterative template refinement to create a group-wise average using the ANTS registration algorithm (93). These steps result in a study-specific template which serves to create a population-specific common space that allows for group-level comparisons. Next, the transform mapping from the common space group average to native image of each subject was calculated to map the minimum deformation required at a voxel-level to map each subject to the average neuroanatomy of the group. The log-transformed Jacobian determinant were estimated at each voxel in the deformation fields and blurred with Gaussian smoothing using 0.2 mm full width at half maximum kernel to better conform to Gaussian assumptions for statistical testing. This method has been described in Chakravarty et al. (2016) (94) and Allemang-Grand et al. (2017) (95) for cross-sectional analysis. The relative Jacobian determinants of the deformation fields were used to measure local anatomical differences. These alterations could either be expansions or contractions and are dependent on the magnitude of the deformation at each voxel.

#### 4.2.2 Quality control

Prior to performing analyses, the images were manually inspected for artifacts (hardware, software, motion artifacts or tissue heterogeneity and foreign bodies) and images with said artifacts were excluded prior to and after pre-processing. Images that passed both levels of quality control were included in the analyses https://github.com/CoBrALab/documentation/wiki/Mouse-QC-Manual-(Structural)). Furthermore, the outputs from the processing techniques were also inspected to assess quality control of the registration by visually assessing the resultant images of the registrations for a proper orientation, size, and sensible group average, and to assess the quality control of the segmentations ensuring that the labels matched the underlying anatomy.

### 4.3 Immunohistochemistry

At 90 dpi, 14 mice (7 mice per injection group; 7 mice per sex) were anaesthetised with isoflurane and transcardially perfused with ice cold 0.9% (wt/vol) PBS followed by ice cold 10% formalin neutral buffer solution for immunohistochemistry. The brains were extracted from the skull and were fixed overnight with 10% formalin then transferred to 70% ethanol solution until the brains were blocked and processed for immunohistochemistry.

Brains of PBS- and formalin neutral buffer solution-perfused mice were sliced in 1 mm thick coronal slices, processed by the Histology Core Facility at the McGill Goodman Cancer Research Centre (http://mcgillgcrc.com/research/facilities/histology), and subsequently the brain slices were embedded in paraffin. Immunohistochemistry was performed on 5 μ-thick serial sections, mounted on glass slides, deparaffinized in xylene baths for 15 minutes and rehydrated through a series of graded ethanol baths (100%, 95%, and 70% ethanol for 10, 10, and 5 minutes respectively). For antigen retrieval, slides were incubated and boiled in a citrate buffer (pH 6.0) for 10 min using a pressure cooker. After cooling, slides were washed with Tris Buffered Saline with Tween 20 (TBS-T) for 5 minutes, then incubated in a 3% hydrogen peroxide solution for 15 minutes to inhibit endogenous peroxidases and washed again with TBS-T for 10 minutes. Slides were blocked at room temperature for 30 minutes with diluted normal goat serum (10% serum in TBS-T), and then incubated with a primary antibody diluted at 1:500 in 5% normal goat serum in TBS-T overnight at 4°C. The primary antibodies used in this study were glial fibrillary acidic protein (GFAP; Source: Dako; Host: Rabbit; https://www.agilent.com/store/productDetail.jsp?catalogId=Z033429-2&catId=SubCat3ECS_86547), ionised calcium-binding adapter molecule 1 (Iba-1; Source: Wako; Host: Rabbit; https://labchem-wako.fujifilm.com/us/product/detail/W01W0101-1974.html), tyrosine hydroxylase (TH; Source: Pel-Freez; Host: Rabbit; https://www.pel-freez.com/p40101-150-ab-rabbit-tyrosine-hydroxylase-p40101-150), and phosphorylated Serine129 alpha-synuclein (pS129Syn; Source: Abcam (ab184674); Host: Mouse; https://www.abcam.com/alpha-synuclein-phospho-s129-antibody-p-syn81a-ab184674.html). A negative control slide was used to test for false positives for each staining process whereby no primary antibody was used. After washing thrice with TBS-T for 5 minutes each time, sections were incubated with a (horseradish peroxidase) HRP-conjugated secondary antibody (1:500 in blocking buffer; 5% serum) for 30 minutes at room temperature. Peroxidase-positive structures were visualised by incubation with DAB (3,3’-diaminobenzidine) (Vector Laboratories; https://vectorlabs.com/products/substrates/dab-hrp-substrate#:~:text=Vector%20DAB%20Substrate%20(3%2C3,a%20gray%2Dblack%20reaction%2 0product.) for a maximum of 10 minutes (on average 3 minutes). After counterstaining with haematoxylin, the slides were dehydrated (reserve of the steps used to deparaffinize and rehydrate the brain slices) and cover slipped. TIFF format images of the cover slipped microscope slides were acquired using a bright field microscope (Olympus DP-21SAL coupled to a digital camera DP21/DP26).

Quantification of DAB positive markers normalised by tissue area (mm^2^) was outsourced and performed by Visikol Inc. (https://visikol.com/). Briefly, images underwent standardised preprocessing to normalise intensity differences between images and a colour deconvolution technique (to identify the brown DAB precipitate (positive particles) and haematoxylin blue particles (cell nuclei)), with noise reduction, was applied to extract the signal of interest for each marker, to determine the total number of positive cells for the markers of interest. Additionally, images were also segmented to determine total tissue pixel area for each ROI region such that the counts for each ROI were normalised to its respective total ROI tissue area (converted from pixel area to millimetres squared).

Quantification of cell markers such as GFAP, Iba-1 and TH yielded higher accuracy (intraclass correlation; ICC values >0.85) counts when compared to manual counts. An in-house validation was performed on all four markers for a small subset of images (n=7 subjects) within two ROIs (bilateral striatum), with ICC > 0.9 for a manual count accuracy measure between two independent raters; S.T. and M.P.). However, for the pS129Syn marker, given that the phosphorylated protein can be in either the cell nucleus or the cell processes (dendrites or axon), such constraints and with the slice thickness, the quantification of positive cell markers were best identified using mainly the intensity feature, whereby we examined DAB positive pixels normalized by tissue area. Specifically, for the pS129Syn-ROI stain-region pairs, the quantification of positive pixels was performed using an in-house ImageJ python script, coded by Dr. Thomas Durcan’s group (https://github.com/neuroeddu/HistQ). This technique yielded a higher ICC accuracy value, ICC=~0.7, compared with an ICC of 0.5 in which was obtained using the Visikol quantification. Similarly, a colour convolution technique was used to determine total number of positive particles and total tissue area (both of which was measured using intensity thresholds, set by the user, which was based on an average of the intensities manually set by the user for each subject image belonging to a stain-region pair). Again, the outputs measured used for analyses were DAB-positive particles normalised per tissue area for the pS129Syn stained images.

### 4.4 Immunofluorescence

Animals were anaesthetised on a lethal overdose of ketamine (100mg/kg) and xylazine (20 mg/kg) in 9% sterile saline. They were cardiac perfused with ice cold 1X PBS followed by cold 4% paraformaldehyde (PFA) (96). Whole brains were stored in 4% PFA for 24 h, then in 15% sucrose in 1X PBS for 24 h, followed by 30% sucrose for an additional 48 h. Finally, brain tissue was frozen in Cryomatrix embedding media (Cat# 67-690-06, Thermo Scientific: Shandon) on dry ice and stored at −80 °C for further use. Brain tissue was cut into 20 μm sections via Leica CM1950 Cryostat and stored free-floating in 1X PBS + 0.02% azide (PBS-N) at 4 °C.

For phospho-S129 aSyn (pS129Syn) and Iba1 immunofluorescent labelling, 20 μm sections were mounted onto Superfrost Plus slides and allowed to dry. Brain sections were washed two times with 1X TBS for 5 minutes each, followed by boiling in 10 mM sodium citrate and 0.02% Tween (pH 6.1) antigen retrieval buffer at 95°C for 20 minutes, next cooled (in a bucket of ice) to room temperature for 30-40 minutes in the same buffer. Then, sections were washed once with 1X TBS, and permeabilized by washing them three times in TBS + 0.2% Triton, 5 minutes each. Nonspecific binding was prevented by incubating sections with 5% donkey serum, 2% normal goat serum in TBS + 0.2% triton at room temperature for 1.5 hours. Sections were then processed for immunostaining by overnight incubation at 4°C in the primary antibodies of anti-Human phospho alpha-synuclein (Ser129) (1:2000, Cat# ab51253, Abcam, RRID:AB_869973), anti-Iba-1 (1:500, Cat# 234 006, SYSY, RRID: AB_2884925) diluted in the blocking buffer. After rinsing three times with TBS + 0.2% Triton sections were incubated with secondary antibodies donkey anti-rabbit Alexa Fluor 647 (1:500, Cat# A-31573, ThermoFisher, RRID:AB_2536183) and goat anti-chicken Alexa Fluor 488 (1:500 Cat# A-11039, Thermofisher, RRID:AB_2534069) for 2 hours at room temperature. Brain sections were washed with TBS, three times, then stained with Hoechst 33342 (1:1000, Cat#62249, ThermoFisher) for 10 minutes, followed by rinsing with TBS two times. Then autofluorescence was quenched using TrueBlack Lipofuscin Autoflursence Quencher (Cat# 23007). Whole section and high-resolution colocalization images were captured using Leica DM6B Thunder Imager with 20X (N.A. 0.8) and 40X (N.A. 0.95) objectives, respectively.

### 4.5 Pairwise visual discrimination reversal and motor behaviour

Details regarding animal care during behavioural tasks such as environmental quality control, food and drink access are described above in the 4.1.2 Behaviour and IF experiments section.

#### 4.5.1 Pairwise visual discrimination task with reversal (PVD-R)

Pairwise visual discrimination task with reversal (PVD-R) was used to evaluate reversal learning as previously described (45, 97). The PVD-R was conducted using the automated Bussey-Saksida touchscreen system for mice (model 80614; Lafayette Instrument, Lafayette, Indiana) and the data collected using ABET II Touch software Vesion 2.20 (Lafayette Instrument, Lafayette, Indiana). Each mouse was scheduled for only one session at about the same time daily. The general procedure, including habituation and the pre-training program for PVD-R in a touch-screen-based automated operant system for mice is described in detail elsewhere (45).

Two separate cohorts of M83 hemizygous mice injected with either saline or aSyn PFFs were evaluated in the PVD-R at two distinct time-points, 49±7 and 84±7 dpi. The first cohort with 7 PBS-injected mice and 8 PFF-injected mice underwent PVD-R with an easy set of images, Marble and Fan. The second cohort with 8 PBS-injected mice and 11 PFF-injected mice performed the PVD-R with the more difficult set of images, diagonal lines. The mice were first habituated and pre-trained in the PVD task (45) before undergoing the stereotaxic surgery for PBS or aSyn PFFs injection. Following the surgery, the mice were given a recovery period of 10 days on free food before food restriction started again.

The evaluation of reversal learning in the touchscreen PVD-R is based on two phases: acquisition and reversal (45). In the acquisition phase, mice were required to choose between a rewarded (S+, fan image) and unrewarded (S-, marble image) stimulus displayed in a mask with two windows placed in front of the touchscreen. A set of more difficult images was also used with a second cohort of mice to confirm the aSyn-associated cognitive impairment. In this experiment, 315-degree diagonal lines were the unrewarded (S-) and 45-degree diagonal lines were the rewarded stimuli (S+) during the acquisition phase. The location of the S+ and S-stimuli was pseudorandomly assigned at either the left or right window and the same stimulus arrangement was not presented more than 3 times. When the mouse touched the S+ (correct), the stimuli were removed, and the strawberry milkshake reward was delivered along with illumination of the magazine light and a tone. An incorrect response (touching the S-image) resulted in a 5 second timeout with the house light on. After an incorrect response, the mouse had to start a correction trial by entering the magazine. Correction trials preserved the left/right arrangement of the S+/S-images from the incorrect trial until a correct choice was made. The results of correction trials did not contribute to the overall trial count or correct/incorrect responses. Each session ended once the mouse completed 30 trials or reached 60 min of testing. The mouse had to achieve at least 80% correct responses (24/30 trials correct) for 2 out of 3 consecutive days in order to reach the acquisition criterion. Once all animals in a cohort had reached the acquisition criterion, they were put back on the task and received 2 further task sessions that served as baseline performance (B1 and B2 in the graphs). Baseline sessions were identical to the PVD acquisition. For the first experiment with the easy set of images, following the baseline sessions, mice were tested on the reversal learning task for 30 trials per session, for 10 sessions. In the reversal phase, the S+ and S-contingencies were reversed, i.e., S+ was the marble image and the S-the fan image. For the second experiment with the more difficult set of images, following the baseline sessions, the mice performed the reversal learning according to the following steps: 3 consecutive days of 10 trials per session (pooled together as the first session); 2 consecutive days of 15 trials (pulled together as the second session); and then 30 trials per session until the controls reach around 80% accuracy. In the reversal phase, the S+ and S-contingencies were reversed, i.e., the S+ was the 315-degree diagonal line and the S-was the 45-degree diagonal line images. Trial initiation and correction trials happened in the same fashion as during acquisition and there were no criteria required for the reversal phase. The session also ended either after completion of 30 trials or in 60 minutes. At approximately 49±7 dpi, the mice were trained on acquisition and then submitted to reversal as described above. The mice were then tested once a week (maintenance) using the same image rewards as the reversal until the second time-point at 84±7 dpi. The reversal maintenance required no criteria to pass. At 84±7 dpi, mice were probed on re-reversal (the previous S+ stimulus in reversal now became the S-stimulus). Number of sessions to reach acquisition criteria, the percentage of accuracy, number of correction trials, correct touch latency (s), reward collection latency and trials completed were recorded. The measurements were compared between PBS-injected M83 mice and PFFs-injected M83 mice.

#### 4.5.2 Motor behavioural tasks

All mice that underwent PVD-R also performed motor behavioural tasks (open-field, wirehang, grip-force and rotarod). The motor behavioural tasks were conducted under food restriction and approximately 63±7 and 98±7 dpi, immediately after the touchscreen testing.

##### 4.5.2.1 Open Field

Locomotor activity in the open field was assessed as previously described (98). Mice were first habituated to the testing room for at least 30 minutes prior to the motor assessment.

Mice were placed in the centre of the open field arena, which was a 20 cm x 20 cm platform surrounded by 30 cm high walls. Mice were allowed to freely explore the open field arena for one hour and the total distance travelled, and the vertical activity were automatically recorded (Omnitech Electronics Inc., Columbus, USA).

##### 4.5.2.2 Wire-Hang

The wire-hang test was conducted to assess motor strength and coordination as described elsewhere (99). Briefly, the mice were placed on a metal grid 45 cm above a large tub filled to the top with wood-chip bedding. Mice were prompted to grip the wires with forepaws by gently vibrating the cage lid. The cage lid was then inverted and the latency to fall from the cage lid was recorded with a 60-second cut-off time. Each mouse was subjected to five trials, spaced at least 5 min apart. The average time to fall was recorded and used for analyses.

##### 4.5.2.3 Grip-force

Grip-force in the forelimbs was measured using Columbus Instruments Grip Strength Meter as previously described (100–101). Mice were allowed to grasp the smooth and triangular pull bar with forelimbs by holding their tails. Mice were then pulled backward in the horizontal plane and the peak force (N) applied to the bar was recorded. Three trials were performed per mouse within the same session and the highest measurement from the three trials was recorded.

##### 4.5.2.4 Rotarod

To investigate motor coordination and balance, mice were tested on the rota-rod as previously described (99). Mice were placed on the rota-rod (San Diego Instruments; San Diego, CA, USA) and rotation was accelerated linearly from 5 to 50 rpm over 5 min with no reverse. Each mouse was tested for ten trials on the first day and four trials on the second day. Mice were returned to home cage and given at least 10-min breaks between trials. Latency to fall was recorded automatically.

### 4.6 Statistical analyses

Statistical analyses were carried out using R software (3.5.0) and the RMINC package (https://wiki.phenogenomics.ca/display/MICePub/RMINC), and all results were corrected for multiple comparisons using the False Discovery Rate (FDR) (55).

#### 4.6.1 Volumetric analyses

Volumetric analyses were performed to examine differences in both whole structures and voxel-wise volumes. We performed univariate analyses to assess group-level differences in neuroanatomy for structure volumes, and voxel-wise differences from the DBM measures. These analyses were performed using linear models to examine the injection group by sex interaction at each region using the volume outputs or at each voxel using the absolute Jacobian outputs.

#### 4.6.2 Whole brain structural covariance analysis

##### 4.6.2.1 Orthogonal projective non-negative matrix factorization

Alpha-synuclein PFF-induced brain atrophy was also examined using orthogonal projective non-negative matrix factorization (OPNMF) (57–60) to examine structural covariance patterns of anatomical variation. OPNMF decomposes an input matrix into two matrices. The input matrix has dimensions m x n, where m is the number of voxels and n the number of subjects. Here, the input matrix was composed of the inverted relative Jacobian determinants z-scored at each voxel for each subject, loaded as columns. The first output matrix is the component matrix W (m x k), where k (defined a priori by the user) represents the number of components/spatial patterns of covariance. This output matrix is composed of the component scores for each voxel, which describes the groupings of voxels sharing a covariance pattern. The second output matrix is the weight matrix H (k x n), which contains the weightings of each subject for each component, thereby describing how each subject loads onto each pattern. The component and weight matrices are constructed such that their multiplication reconstructs the input data as best as possible by minimising the reconstruction error between the original input and the reconstructed input. With the orthogonality constraint, this variant of NMF ensures output spatial components are non-overlapping, and that each output component represents a distinct pattern of voxels. Thus, a specific voxel can only be part of one component, however the voxels within a region can be included in more than one component. Using the component weighting output matrix, the spatial pattern of voxel scores for each component can be plotted onto the mouse brain average. Injection group differences in subject weights for each component were examined using general linear models, with sex modelled as a covariate.

To select the optimal number of components (k) to analyse, we emulated the stability analyses performed in Patel et al. (2020) (57). Two measures are commonly examined when assessing stability and determining k. First, accuracy is measured by observing reconstruction error of a decomposition, defined as the Frobenius norm of the element-wise difference between the original and reconstructed inputs. We plot the gradient in reconstruction error, enabling quantification of the gain in accuracy provided by increasing the number of components from one granularity to the next. Second, stability of a decomposition is measured by assessing the similarity of output spatial components across varying splits of subjects. To track stability, various OPNMF is performed on various subsets of subjects and the spatial similarity of the resulting outputs describes how consistent results are across the sample. However, given the relatively small number of subjects in our study, assessing stability of OPNMF across different subsets of subjects was not feasible and thus bypassed. Reconstruction error generally increases with k given the increased degrees of freedom and greater number of covariance patterns detected. However, a higher k is also likely to lead to overfitting of patterns specific to a single or few subjects. Thus, we select k by choosing the smallest k that can capture the major patterns of interest. Here, k=4 was selected (see section 5.2 in supplementary material; Supplementary Figure 6).

#### 4.6.3 Immunohistochemistry analysis

Regions of interest were selected majorly based on where MRI-derived atrophy was observed (as determined by volumetric (MAGeT) and/or network analyses (OPNMF)), as well as in a small number of regions where we did not observe any MRI-derived differences as a result of the inoculum received (namely, the anterior hippocampus and superior medulla). Analyses were performed separately for each of the four stains (GFAP, Iba-1, pS129Syn, and TH), for each of the following brain regions: primary motor cortex, primary sensory cortex, hippocampus, hypothalamus, medulla (superior and inferior), midbrain reticular nucleus, nucleus accumbens, paraventricular nucleus of hypothalamus, periaqueductal gray, pons (anterior and posterior), substantia nigra, striatum (level of injection), thalamus (anterior and posterior).

The data for some stain-region pairs did not satisfy the assumption of normal distribution for parametric analyses, even after cubed root transformation of the data. Most of the data distributions were right-skewed and peaking at the values approaching zero. For these data, nonparametric linear models were performed, whereas parametric linear models were performed for the stain-region pairs that did satisfy normal distribution prior to or after cubed root transformation. For all analyses, DAB-positive markers (normalized by tissue area) were assessed using a linear model with respect to injection group and sex. Moreover, analyses were not performed for the left side of the pons for pS129Syn, and the left side of the posterior thalamus for Iba-1 as there was not enough data available due to common artifacts (such as tissue ripping, lost, and folding onto itself occurring during the staining process).

#### 4.6.4 Cognitive and behavioural analysis

In the PVD-R, the number of sessions to reach acquisition criteria was analysed by unpaired two-tailed t test; the percentage of accuracy, number of correction trials, correct touch latency (s), reward collection latency and trials completed were analysed by repeated measures two-way ANOVA. Data normality was evaluated using Shapiro-Wilk test. Comparisons between groups in the motor tasks were made by the Student’s t-test, when the data were determined to follow a normal distribution, or the Mann-Whitney test when variables did not follow a normal distribution. Data were analysed using GraphPad Prism v8 software and expressed as mean ± SEM.

## Acknowledgments

We would like to thank a few members of the Early Drug Discovery Unit at McGill; Emmanuelle Nguyen-Renou for her help with PFF production, and Vincent Soubannier, Frederique Larroquette, Rhalena Amber Thomas, and Eddie Cai for their support with immunohistochemistry quantification.

S.T. was awarded a Healthy Brains for Healthy Lives fellowship and was awarded a Fonds de Recherche du Quebec en Sante (FRQS) doctoral training scholarship. E.A.F. is supported by a CIHR Foundation grant (FDN-154301) and by a Canada Research Chair 1315 (Tier 1) in Parkinson’s disease. L.M.S. and T.J.B. received support from the Canadian Institutes of Health Research, the Natural Science and Engineering Research Council of Canada and the BrainsCAN Canada First Research Excellence Fund. L.M.S. is a Tier I Canada Research Chair in Translational Cognitive Neuroscience and a CIFAR Fellow in the Brain, Mind and Consciousness program. T.J.B. is a Western Research Chair. A.S.M. received a BrainsCAN Postdoctoral fellowship and a Return Home fellowship from the International Society for Neurochemistry. A.A. received a Parkinson’s Society Post-Doctoral fellowship. M.A.M.P. and V.F.P. received support from the Canadian Institutes of Health Research (CIHR, PJT 162431, PJT 159781, PJ2 179755), Natural Science and Engineering Research Council of Canada (NSERC; 06577-2018 RGPIN; 03592-2021 RGPIN), CFI/ORF, The Tanenbaum Open Science Institute (TOSI), a BrainsCAN/Healthy Brains for Healthy Lives (HBHL; Canada First Research Excellence Fund to Western/McGill University) Initiative for Translational Neuroscience Award, and BrainsCAN Research Excellence Award Awards, as well as support for the Rodent Cognitive and Innovation Core for behaviour experiments. M.A.M.P. is a Tier I Canada Research Chair in Neurochemistry of Dementia. M.M.C receives salary support from the FRQS (Chercheur Boursier Junior 2), NSERC, CIHR, and BrainsCAN/HBHL Initiative for Translational Neuroscience Award, and HBHL.

