## Supplementary Information for "Presymptomatic neuroanatomical and cognitive biomarkers of alpha-synuclein propagation in a mouse model of synucleinopathy"

### **Supporting Information for**
### Presymptomatic neuroanatomical and cognitive biomarkers of alpha-synuclein propagation in a mouse model of synucleinopathy

Stephanie Tullo, Aline S Miranda, Esther del Cid-Pellitero, Mei Peng, Daniel Gallino, Anoosha Attaran, Raihaan Patel, Vladislav Novikov, Megan Park, Flavio Beraldo, Timothy J. Bussey, Lisa M. Saksida, Edward A. Fon, Vania F. Prado, Marco A.M. Prado, M. Mallar Chakravarty

Stephanie Tullo, M.Mallar Chakravarty. *Cerebral Imaging Center,* Douglas Mental Health University Institute, Verdun, Canada, H4H 1R3. 514-761-6131 ext 4781. Marco A.M. Prado, *Robarts Research Institute*, London, Canada, N6A 5B7. 519-931-5777 ext 24888


#### **Supporting Information Text**

##### 5.1 Voxel-based deformation analysis

For voxel-wise measures of atrophy, we observed subtle yet widespread effects between the two groups. At FDR 20%, we observed voxels in the left and right striatum, right primary and secondary motor cortex, bilateral anterior cingulate cortex, bilateral primary somatosensory cortex, bilateral thalami (encapsulating almost the entirety of the thalamus; however more focused on the anterior and lateral parts), bilateral hippocampi (primarily in the CA1 and dentate gyrus subregions), the periaqueductal gray, bilateral substantia nigra, bilateral midbrain reticular nuclei (albeit more of the right hemisphere), as well as subregions in the pons and medulla (Supplementary Figure 2).

##### 5.2 OPNMF stability analysis

To select the optimal number of components, k, to analyse, we assessed accuracy at various granularities (from k=3 to k=10) (1), whereby accuracy was measured by observing the gradient in reconstruction error. While there was a gain in accuracy by increasing the number of components from one granularity to the next, to reduce the likelihood of overfitting, k=4 was selected using regression models to track which granularity best discriminates the two injection groups (PBS vs PFF-injected mice), given the data can be separated into 4 groups (2 injection groups and 2 sexes). The gain in accuracy from k=3 to k=4 (plot A) is also much larger than gain in accuracy for further increasing k. Thus, a choice of k=4 also balances accuracy (lower error) vs. complexity (more components).

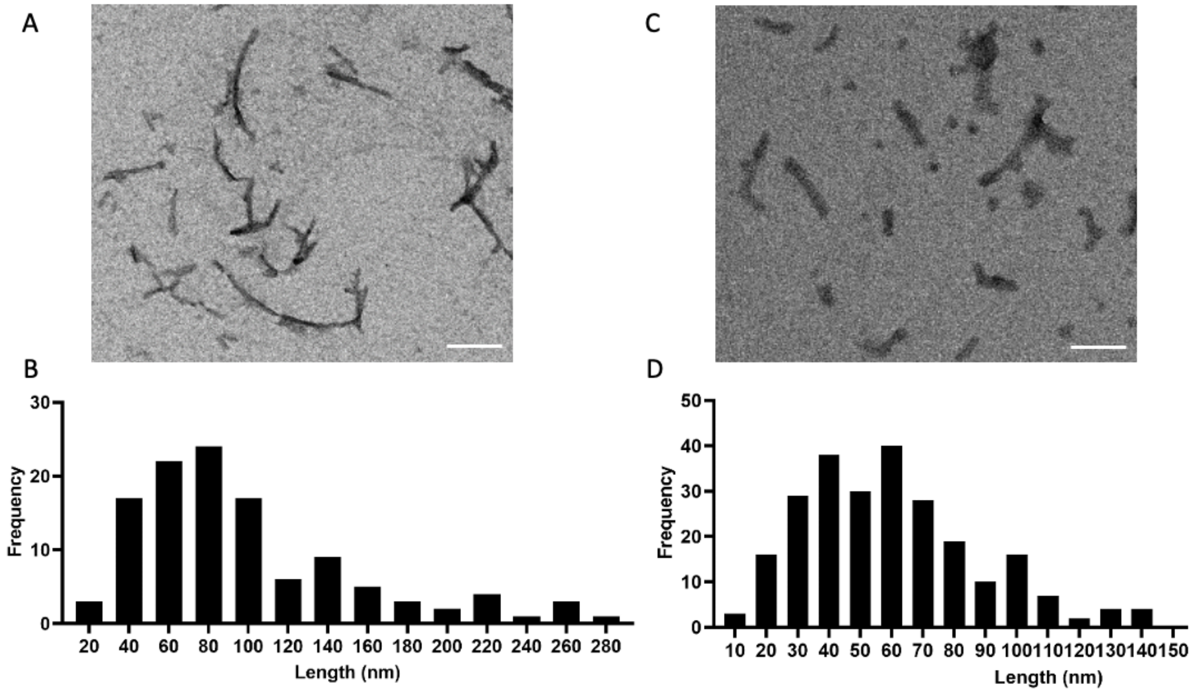


**Supplementary Figure 1.** Human alpha-synuclein- preformed fibrils (syn-PFFs) characterization. [A,C] Representative photomicrographs of human syn-PFFs non-sonicated [A] and 30 sec sonicated [B] staining by negative staining and visualized using Tecnai 12 BioTwin 120 kV electron microscope. [B, D] Histograms showed the syn- PFFs length distribution measured using ImageJ software and their distribution plotted using GraphPad Prism software. [B] Human syn-PFFs non-sonicated (n=117, length average= 99.20 nm, median length= 81.08 nm, minimal length = 26.94 nm, maximal length= 273.63 nm). [D] Human syn- PFFs sonicated for 30 seconds (n= 252, length average= 62.50 nm, median length= 57.46nm, minimal length= 10.91 nm, maximal length= 207.89 nm). Bar scale for [A] is 200 nm, [C] is 100 nm.

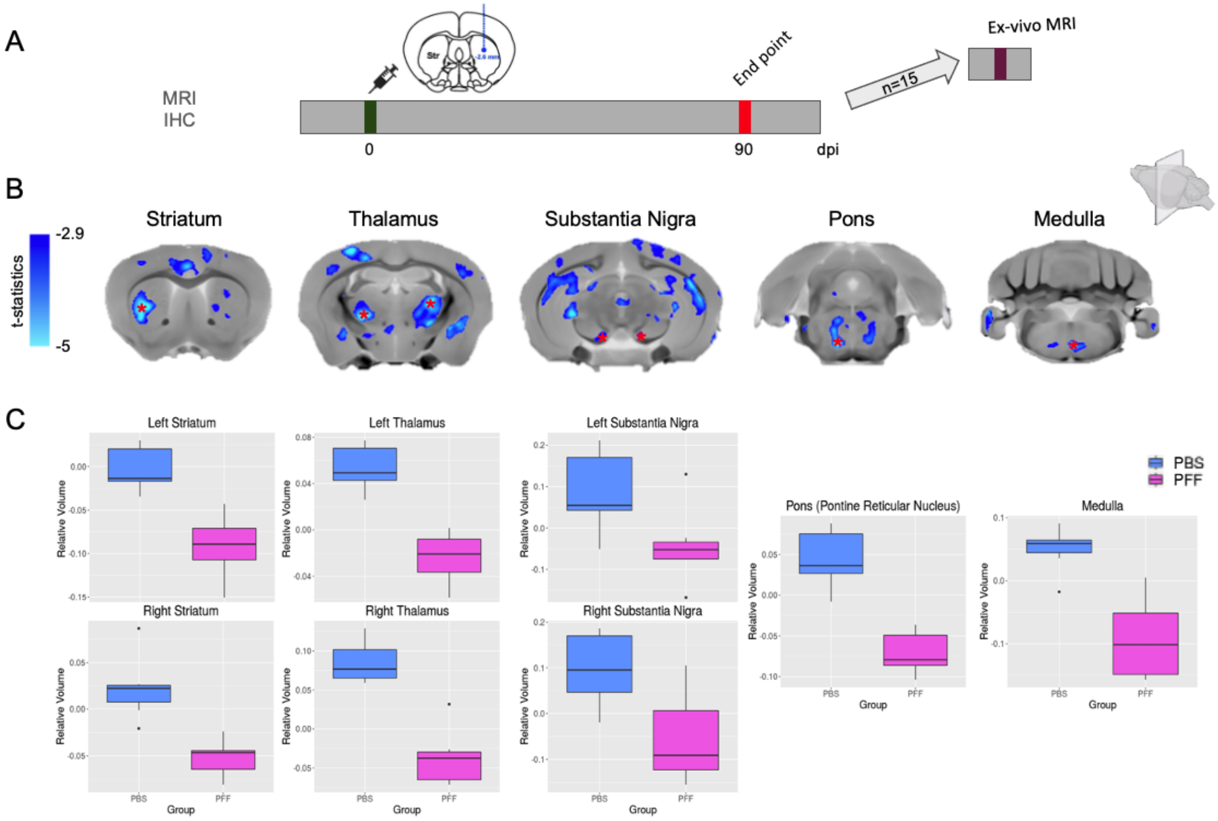


**Supplementary Figure 2. Voxel-wise volumetric differences between PFF- and PBS-injected mice.** [A] Experimental timeline for mice that underwent *ex vivo* MRI imaging (n=15). [B] At FDR 20%, we observed subtle effects when examining voxel-wise volume differences between the two groups. Smaller voxels for the PFF-injected mice were observed in some key structures known to project to or receive input from the injection site (right striatum) (regions in B). [C] Box plots representing voxel-wise differences between the PBS and PFF injected mice in the injection site, the contralateral striatum (left), the left and right thalamus, bilateral substantia nigra, as well as regions in the pons and medulla.

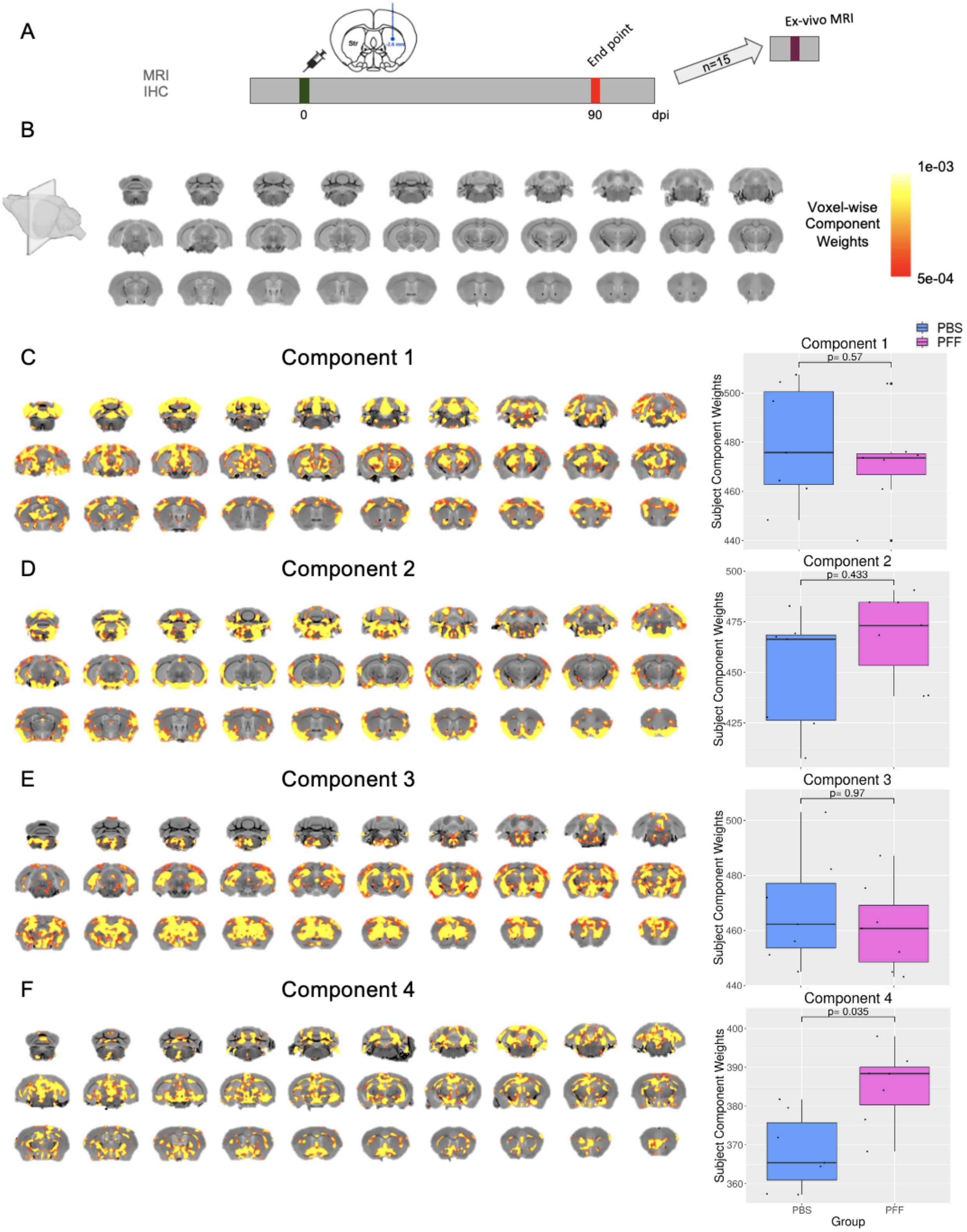


**Supplementary Figure 3. Results of the 4 components OPNMF run.** [A] Experimental timeline for mice that underwent *ex vivo* MRI imaging (n=15). [B] Coronal slices of a mouse brain average displayed from posterior to anterior slices [C-F] OPNMF decomposition of voxel-based deformation in M83 mice for all 4 components. Colourmap denotes voxel-wise component weights. For each component, the spatial pattern of voxel component scores plotted onto coronal slices of an average mouse brain (posterior to anterior slices), depicts the networks of voxels sharing a similar variance pattern (left) and group differences of subject component weightings, describing how each subject loads onto the identified atrophy pattern were assessed using general linear models (right). [C] Component 1 revealed a thalamo-cerebellar pattern with frontal cortex involvement. [D] Component 2 revealed a ventral cortex-pons-medulla pattern. [E] Component 3 revealed a subcortical pattern mainly composed of striatum and hippocampal voxels. [F] Component 4 revealed a striatal-nigral-hypothalamic pattern. Component 4 was the only component with significant group differences in the subject-wise component weights, such that the pattern was driven by PFF-injected mice ($p=0.035$).

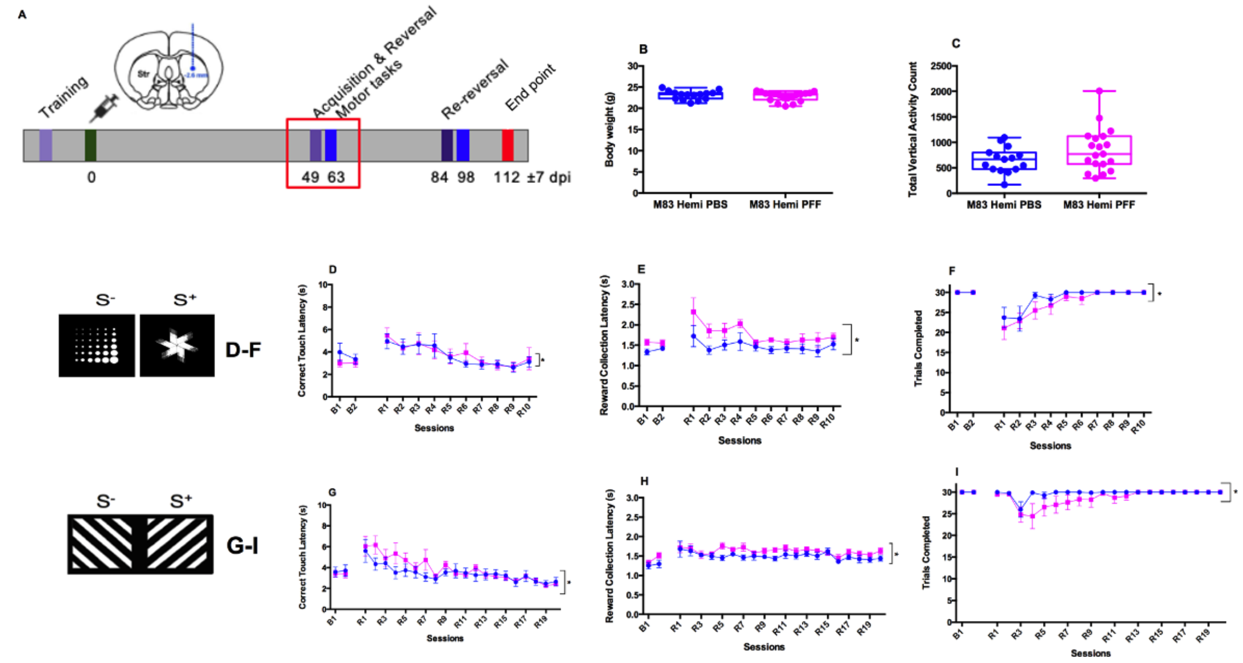


**Supplementary Figure 4. Additional parameters for motor and cognitive behaviour at 49±7 and 63±7 days post injection.** [A] Experimental design; [B] Body weight (g) (Unpaired t test: t=0.8363 df=32 p=0.4092); [C] Total vertical activity count in the open field task for 60 minutes (Unpaired t test: t=1.636 df=32 p=0.1116). PBS-injected M83 mice (n=15) or PFFs-injected M83 mice (n=19). [D] Correct touch latency (s) (Repeated measures two-way ANOVA: Session: F (4.063, 51.34) = 4.932, p = 0.0018; Genotype: F (1, 13) = 0.01394, p = 0.9078; Interaction: F (11, 139) = 0.5417, p = 0.8719); [E] Reward collection latency (s) (Repeated measures two-way ANOVA: Session: F (3.466, 43.49) = 3.063, p = 0.0317; Genotype: F (1, 13) = 8.788, p = 0.0110; Interaction: F (11, 138) = 0.5742 = 0.5417, p = 0.8472); [F] Number of trials completed (Repeated measures two-way ANOVA: Session: F (2.390, 30.42) = 8.578, p = 0.0006; Genotype: F (1, 13) = 1.304, p = 0.2740; Interaction: F (11, 140) = 0.4391, p = 0.9359) by PBS-injected M83 mice (n=7) or PFFs-injected M83 mice (n=8) in the PVD reversal learning (PVD-R) task. Marble (S-) and fan (S+) stimuli were used during the PVD-R acquisition phase. Parameters were measured across baseline days 1 and 2 (B1, B2) and reversal days 1 to 10 (R1-R10). [G] Correct touch latency (s) (Repeated measures two-way ANOVA: Session: F (3.693, 62.60) = 7.272, p = 0.0001; Genotype: F (1, 17) = 0.5530, p = 0.4672; Interaction: F (21, 356) = 1.051, p = 0.4010); [H] Reward collection latency (s) (Repeated measures two-way ANOVA: Session: F (6.277, 105.2) = 3.351, p = 0.0041; Genotype: F (1, 17) = 3.736, p = 0.0701; Interaction: F (21, 352) = 0.6577, p = 0.8734); [I] Number of trials completed (Repeated measures two-way ANOVA: Session: F (1.894, 32.19) = 3.498, p = 0.0445; Genotype: F (1, 17) = 1.470, p = 0.2419; Interaction: F (21, 357) = 1.249, p = 0.2072) by PBS-injected M83 mice (n=8) or PFFs-injected M83 mice (n=11) in the PVD-R learning task. Diagonal lines facing down (S-) and diagonal lines facing up (S+) stimuli were used during the PVD-R acquisition phase. Same stimuli were reversed with diagonal lines facing down shown as the (S+) and diagonal lines facing up as the (S-) during reversal (G-I). Parameters were measured across baseline days 1 and 2 (B1, B2) and reversal days 1 to 20 (R1-R20). Results are expressed as mean ± SEM, Unpaired t test (B,C) and Repeated measures two-way ANOVA (D-I). Asterisks indicate statistical differences between groups: *p<0.05.

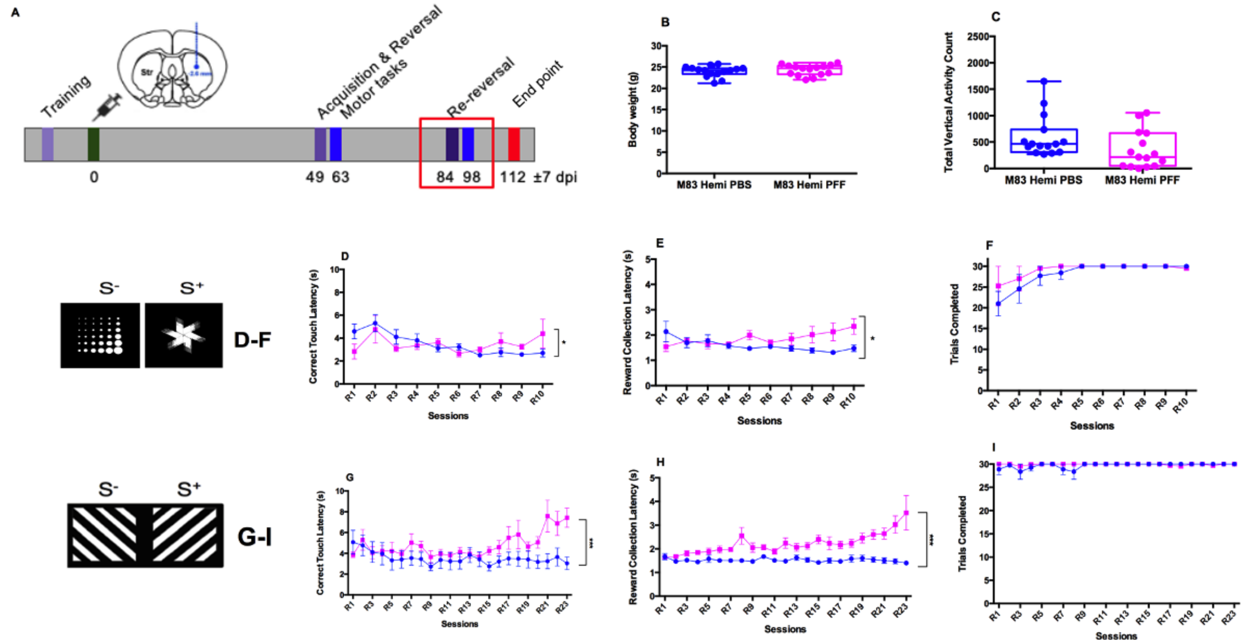


**Supplementary Figure 5. Additional parameters for motor and cognitive behaviour at 84±7 and 98±7 days post injection.** [A] Experimental design; [B] Body weight (g) (Unpaired t test: t=0.7332 df=28 p=0.4695); [C] Total vertical activity count in the open field task for 60 minutes (Unpaired t test: t=1.849 df=28 p=0.0751) of PBS-injected M83 mice (n=15) and PFFs-injected M83 mice (n=15). [D] Correct touch latency (s) (Repeated measures two-way ANOVA: Session: F (2.574, 22.88) = 4.109, p = 0.0221; Genotype: F (1, 9) = 0.001007, p = 0.9754; Interaction: F (9, 80) = 2.408, p = 0.0180); [E] Reward collection latency (s) (Repeated measures two-way ANOVA: Session: F (3.068, 26.93) = 0.6341, p = 0.6029; Genotype: F (1, 9) = 2.737, p = 0.1324; Interaction: F (9, 79) = 3.356, p = 0.0016); [F] Number of trials completed (Repeated measures two-way ANOVA: Session: F (1.520, 13.68) = 3.990, p = 0.0524; Genotype: F (1, 9) = 0.4738, p = 0.5086; Interaction: F (9, 81) = 0.4197, p = 0.9211) by PBS-injected M83 mice (n=7) and PFFs-injected M83 mice (n=4) in the PVD reversal learning (PVD-R) task. Parameters were measured across re-reversal days 1 to 10 (R1-R10). [G] Correct touch latency (s) (Repeated measures two-way ANOVA: Session: F (3.747, 62.85) = 3.450, p = 0.0148; Genotype: F (1, 17) = 2.349, p = 0.1438; Interaction: F (22, 369) = 4.025, p<0.0001); [H] Reward collection latency (s) (Repeated measures two-way ANOVA: Session: F (3.025, 50.74) = 3.092, p = 0.0347; Genotype: F (1, 17) = 19.14, p = 0.0004; Interaction: F (22, 369) = 3.918, p<0.0001); [I] Number of trials completed (Repeated measures two-way ANOVA: Session: F (1.716, 28.70) = 1.266, p = 0.2927; Genotype: F (1, 17) = 1.166, p = 0.2952; Interaction: F (22, 368) = 1.059, p=0.3907) by PBS-injected M83 mice (n=8) and PFFs-injected M83 mice (n=11) in the PVD-R task. Parameters were measured across re-reversal days 1 to 23 (R1-R23). Results are expressed as mean ± SEM, Unpaired t test (B,C) and Repeated measures two-way ANOVA (D-I). Asterisks indicate statistical differences between groups: *p<0.05, ***p<0.0001.

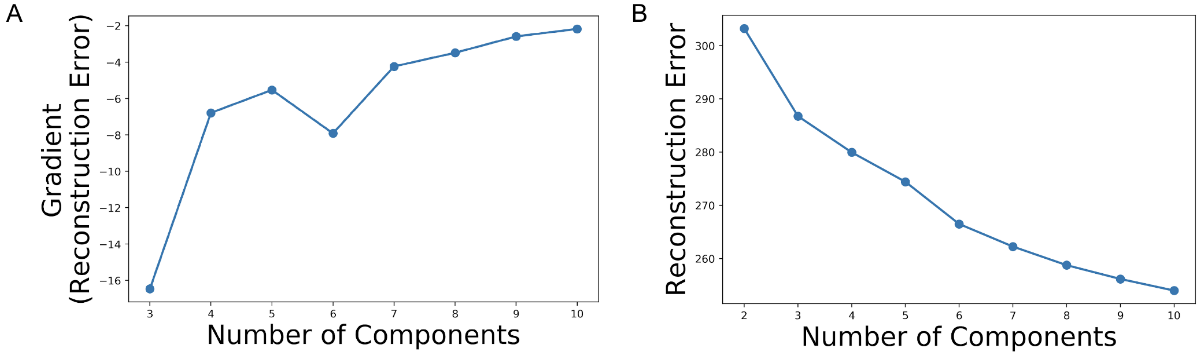


**Supplementary Figure 6. Accuracy measures for k=3 to 10 OPNMF component runs.** [A] Gradient in reconstruction error is the quantification of the gain in accuracy provided by increasing the number of components from one granularity to the next. The biggest gain in accuracy is observed when moving from k=3 to k=4 components. [B] The reconstruction error trends such that the higher the k, the greater accuracy. While the maximum number of components is the number of columns minus one, however at the maximum number of components there is a greater propensity of overfitting and therefore specificity to the current data (thereby less generalizability).

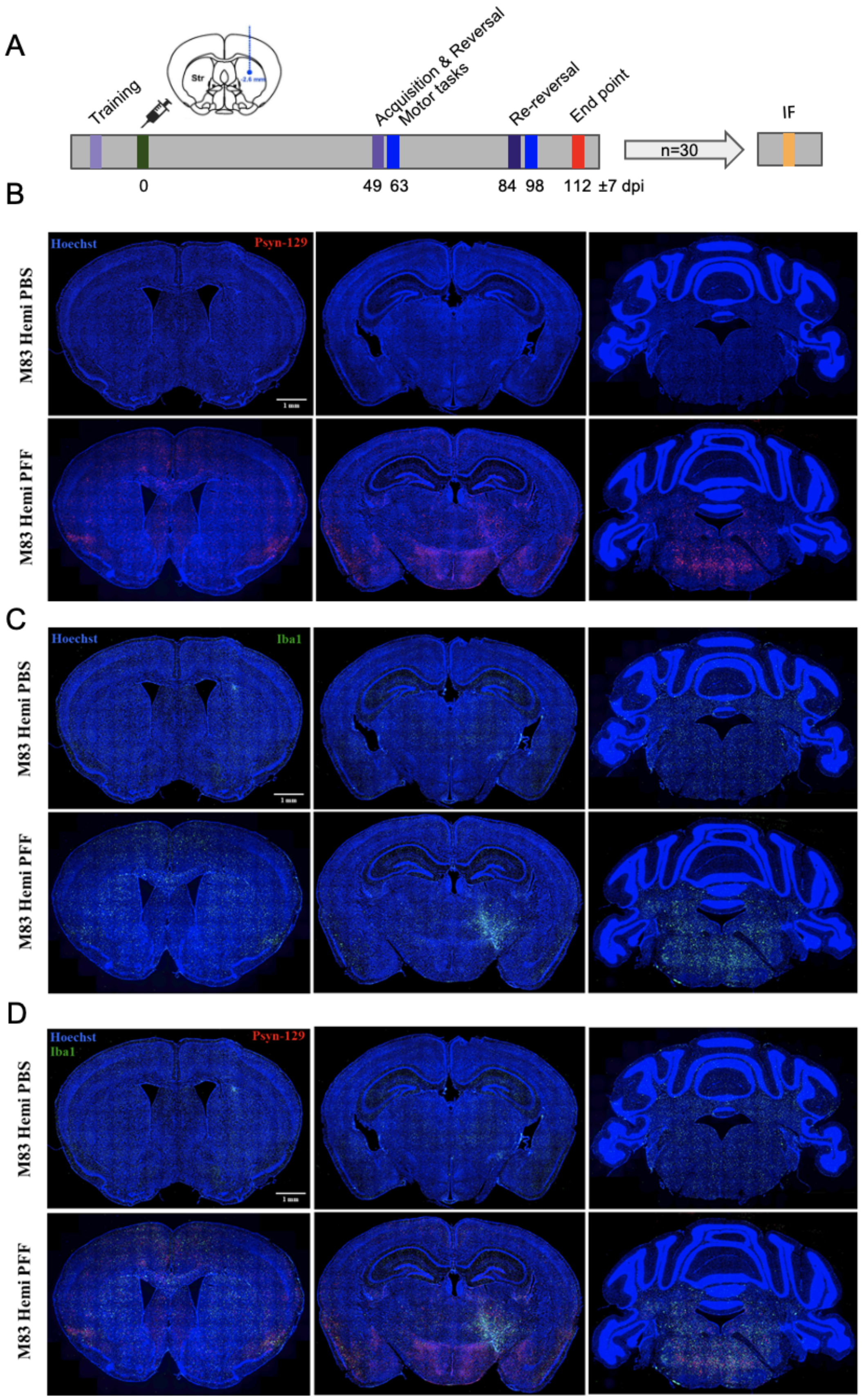
A
Training
Str
-2.6 mm
Acquisition & Reversal
Motor tasks
Re-reversal
End point
IF
n=30
0
49 63
84 98
112 ±7 dpi
B
Hoechst
Psyn-129
M83 Hemi PBS
M83 Hemi PFF
1 mm
C
Hoechst
Iba1
M83 Hemi PBS
M83 Hemi PFF
1 mm
D
Hoechst
Iba1
Psyn-129
M83 Hemi PBS
M83 Hemi PFF
1 mm

**Supplementary Figure 7. Spreading of alpha-synuclein and microglial reaction in brain regions of M83 mice after behavioural assessment.** [A] Experimental timeline. [B-D] Representative images of [C] pSyn129 (red) and [B] Iba-1 (microglia, green) immunolabelling with DAPI nuclear marker (blue) in coronal sections from PBS-injected M83 mice or PFFs-injected M83 mice. [D] Merged image with psyn129 (red) and Iba-1 (green) immunolabelling with Hoechst nuclear marker (blue). Images shown here are near the coordinates of injection. Ipsilateral and contralateral hemispheres are labelled as respective to right hemisphere intrastriatal injection. Tiling images of the whole section taken at 20X. Scale bar is 1 mm.

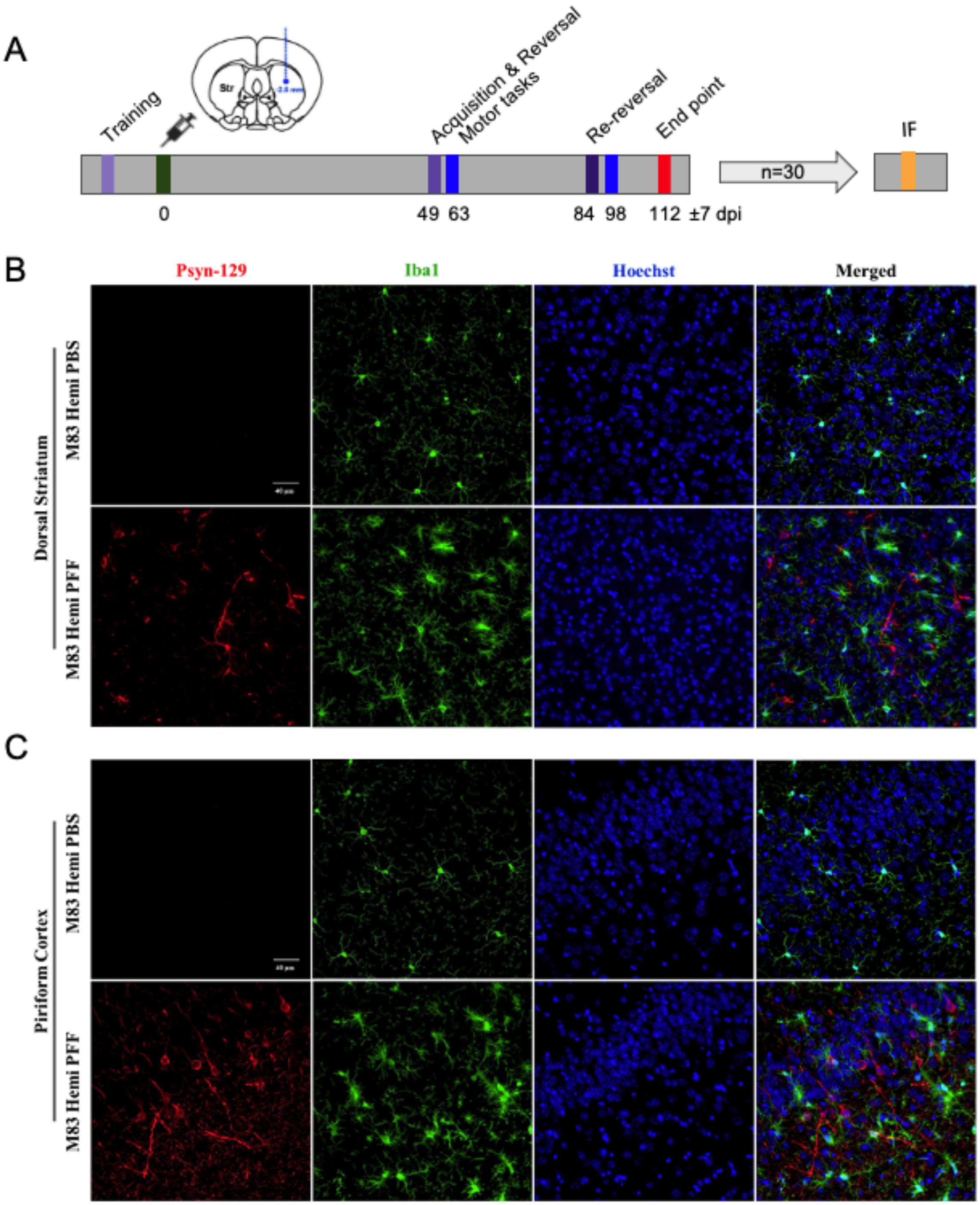


**Supplementary Figure 8. Immunoreactivity of psyn129 and Iba-1 in the brain of M83 mice.** [A] Experimental timeline. [B-C] Representative high-resolution microscopy immunofluorescence images in the [B] ipsilateral dorsal striatum (region of injection) and in the [C] piriform cortex of PBS-injected M83 mice or PFFs-injected M83 mice labelling with pS129Syn (alpha-synuclein, red), Iba-1 (microglia, green) and Hoechst (nuclear mark, blue). Images taken at 40X. Scale bar is 40 µm.

**Supplementary Table 1.** Volumetric statistical results for each structure in the Allen Mouse Brain Atlas in terms of standardised betas, t-, p- and q- values.

| region | beta-groupPFF | tvalue-groupPFF | pvalue-groupPFF | qvalue-tvalue-groupPFF |
|---|---|---|---|---|
| right medial forebrain bundle | -0.0038 | -6.2156 | 6.97E-14 | 0.0359 |
| right Reticular nucleus of the thalamus | -0.0993 | -5.9643 | 7.01E-14 | 0.0359 |
| right Basolateral amygdalar nucleus, posterior part | -0.0227 | -5.2727 | 7.10E-14 | 0.0438 |
| right Ventral posterolateral nucleus of the thalamus | -0.0695 | -5.2295 | 7.15E-14 | 0.0438 |
| left Prosubiculum | -0.0895 | -5.2184 | 7.06E-14 | 0.0438 |
| right Primary somatosensory area, barrel field | -0.1713 | -5.0072 | 1.05E-13 | 0.0499 |
| right Rostrolateral area | -0.0390 | -4.8418 | 1.07E-13 | 0.0499 |
| left Cortical amygdalar area, posterior part, medial zone | -0.0653 | -4.6971 | 7.84E-14 | 0.0499 |
| right Primary auditory area | -0.0876 | -4.6596 | 1.00E-13 | 0.0499 |
| right auditory radiation | -0.0125 | -4.5608 | 1.09E-13 | 0.0499 |
| right Basomedial amygdalar nucleus, posterior part | -0.0326 | -4.5194 | 9.09E-14 | 0.0499 |
| right Orbital area, ventrolateral part | -0.1237 | -4.5090 | 1.04E-13 | 0.0499 |
| right Ventral anterior-lateral complex of the thalamus | -0.0564 | -4.4569 | 9.67E-14 | 0.0499 |
| right Central amygdalar nucleus | -0.0504 | -4.3918 | 9.48E-14 | 0.0499 |
| right Supplemental somatosensory area | -0.3622 | -4.3379 | 1.01E-13 | 0.0499 |
| right Ventral auditory area | -0.0742 | -4.2561 | 1.04E-13 | 0.0499 |
| left Medial geniculate complex, dorsal part | -0.0079 | -4.2531 | 8.33E-14 | 0.0499 |
| left Reticular nucleus of the thalamus | -0.0803 | -4.2194 | 7.24E-14 | 0.0499 |
| right Secondary motor area | -0.6425 | -4.2188 | 9.74E-14 | 0.0499 |
| left Ventral anterior-lateral complex of the thalamus | -0.0563 | -4.1939 | 7.74E-14 | 0.0499 |
| right Orbital area, medial part | -0.0831 | -4.1257 | 9.35E-14 | 0.0499 |
| right internal capsule | -0.1016 | -4.0632 | 8.79E-14 | 0.0499 |
| left Rostrolateral area | -0.0528 | -4.0498 | 8.55E-14 | 0.0499 |
| right Dorsal auditory area | -0.0403 | -4.0248 | 9.41E-14 | 0.0499 |
| left Primary somatosensory area, unassigned | -0.0497 | -4.0230 | 8.44E-14 | 0.0499 |

| | | | | |
|---|---|---|---|---|
| right Ventral medial nucleus of the thalamus | -0.0825 | -4.0063 | 9.81E-14 | 0.0499 |
| right Orbital area, lateral part | -0.1628 | -3.9963 | 9.22E-14 | 0.0499 |
| right Subthalamic nucleus | -0.0119 | -3.9915 | 9.28E-14 | 0.0499 |
| right Parasubthalamic nucleus | -0.0109 | -3.9434 | 9.16E-14 | 0.0499 |
| left Nucleus sagulum | -0.0044 | -3.8999 | 7.29E-14 | 0.0499 |
| right Ventral posteromedial nucleus of the thalamus | -0.1499 | -3.8975 | 9.95E-14 | 0.0499 |
| Nucleus raphe pontis | -0.0087 | -3.8911 | 8.73E-14 | 0.0499 |
| left Subparaventricular zone | -0.0060 | -3.8828 | 7.39E-14 | 0.0499 |
| left Subiculum | -0.1287 | -3.8783 | 7.58E-14 | 0.0499 |
| left Posterior hypothalamic nucleus | -0.0325 | -3.8736 | 8.16E-14 | 0.0499 |
| left Ventral posterolateral nucleus of the thalamus | -0.0719 | -3.8574 | 8.00E-14 | 0.0499 |
| left Basolateral amygdalar nucleus, ventral part | -0.0169 | -3.8222 | 7.54E-14 | 0.0499 |
| right Posterior hypothalamic nucleus | -0.0394 | -3.8183 | 1.03E-13 | 0.0499 |
| left Infralimbic area | -0.0461 | -3.8168 | 7.95E-14 | 0.0499 |
| left Central amygdalar nucleus | -0.0628 | -3.8166 | 7.64E-14 | 0.0499 |
| right Temporal association areas | -0.1120 | -3.8108 | 8.85E-14 | 0.0499 |
| right Prosubiculum | -0.0615 | -3.8075 | 1.08E-13 | 0.0499 |
| left Intertrigeminal nucleus | -0.0022 | -3.7913 | 8.61E-14 | 0.0499 |
| right Visceral area | -0.0882 | -3.7648 | 1.02E-13 | 0.0499 |
| right Lateral preoptic area | -0.0200 | -3.7408 | 8.91E-14 | 0.0499 |
| right Primary somatosensory area, mouth | -0.2160 | -3.7361 | 1.02E-13 | 0.0499 |
| left Ventromedial hypothalamic nucleus | -0.0216 | -3.7351 | 7.89E-14 | 0.0499 |
| right Primary motor area | -0.4224 | -3.7332 | 9.03E-14 | 0.0499 |
| left Anterior area | -0.0724 | -3.7265 | 8.50E-14 | 0.0499 |
| left Hippocampo-amygdalar transition area | -0.0261 | -3.7126 | 8.67E-14 | 0.0499 |
| left Primary somatosensory area, barrel field | -0.2570 | -3.7101 | 8.27E-14 | 0.0499 |
| left Dorsomedial nucleus of the hypothalamus | -0.0178 | -3.6943 | 8.11E-14 | 0.0499 |
| left external capsule | -0.0366 | -3.6730 | 7.69E-14 | 0.0499 |
| right Infralimbic area | -0.0466 | -3.6722 | 9.88E-14 | 0.0499 |

| | | | | |
|---|---|---|---|---|
| right Anteromedial visual area | -0.0466 | -3.6542 | 8.97E-14 | 0.0499 |
| left cuneate fascicle | -0.0053 | -3.6320 | 7.49E-14 | 0.0499 |
| left Basomedial amygdalar nucleus, posterior part | -0.0274 | -3.6210 | 7.34E-14 | 0.0499 |
| left Cortical amygdalar area, posterior part, lateral zone | -0.0501 | -3.6095 | 7.79E-14 | 0.0499 |
| left Posterior amygdalar nucleus | -0.0521 | -3.6089 | 8.05E-14 | 0.0499 |
| left Endopiriform nucleus, dorsal part | -0.0616 | -3.5981 | 8.21E-14 | 0.0499 |
| right Pedunculopontine nucleus | -0.0581 | -3.5944 | 1.06E-13 | 0.0499 |
| right Primary somatosensory area, nose | -0.0850 | -3.5917 | 9.54E-14 | 0.0499 |
| right external capsule | -0.0409 | -3.5857 | 9.61E-14 | 0.0499 |
| left Lateral hypothalamic area | -0.0807 | -3.5803 | 7.20E-14 | 0.0499 |
| left Parasubthalamic nucleus | -0.0110 | -3.5767 | 7.44E-14 | 0.0499 |
| right Anteromedial nucleus, ventral part | -0.0082 | -3.5764 | 1.07E-13 | 0.0499 |
| left Entorhinal area, lateral part | -0.2239 | -3.5737 | 8.38E-14 | 0.0499 |
| right Lateral hypothalamic area | -0.0964 | -3.5635 | 1.10E-13 | 0.0500 |
| right Subiculum | -0.0898 | -3.5546 | 1.11E-13 | 0.0501 |
| right Field CA1 | -0.3198 | -3.5292 | 1.14E-13 | 0.0507 |
| right Laterodorsal tegmental nucleus | -0.0103 | -3.5105 | 1.13E-13 | 0.0507 |
| right Agranular insular area, posterior part | -0.1123 | -3.5049 | 1.12E-13 | 0.0507 |
| left Ventral part of the lateral geniculate complex | -0.0284 | -3.5047 | 1.11E-13 | 0.0507 |
| right Cortical amygdalar area, posterior part, medial zone | -0.0701 | -3.5046 | 1.15E-13 | 0.0507 |
| right Zona incerta | -0.0943 | -3.5003 | 1.15E-13 | 0.0507 |
| left principal mammillary tract | -0.0032 | -3.4719 | 1.19E-13 | 0.0511 |
| right Caudoputamen | -1.2259 | -3.4462 | 1.22E-13 | 0.0511 |
| right Medial geniculate complex, medial part | -0.0132 | -3.4460 | 1.24E-13 | 0.0511 |
| left Ventral posteromedial nucleus of the thalamus | -0.1329 | -3.4411 | 1.18E-13 | 0.0511 |
| right Posterior complex of the thalamus | -0.0890 | -3.4370 | 1.23E-13 | 0.0511 |
| left Lateral preoptic area | -0.0227 | -3.4365 | 1.16E-13 | 0.0511 |
| Nucleus raphe pallidus | -0.0117 | -3.4356 | 1.21E-13 | 0.0511 |

| | | | | |
|---|---|---|---|---|
| left rubrospinal tract | -0.0210 | -3.4221 | 1.20E-13 | 0.0511 |
| right Peritrigeminal zone | -0.0190 | -3.4210 | 1.25E-13 | 0.0511 |
| left Endopiriform nucleus, ventral part | -0.0337 | -3.4186 | 1.20E-13 | 0.0511 |
| left Magnocellular reticular nucleus | -0.0329 | -3.4174 | 1.17E-13 | 0.0511 |
| right Parabrachial nucleus | -0.0566 | -3.3983 | 1.26E-13 | 0.0523 |
| left Peripeduncular nucleus | -0.0039 | -3.3751 | 1.27E-13 | 0.0539 |
| left Basomedial amygdalar nucleus, anterior part | -0.0393 | -3.3681 | 1.29E-13 | 0.0539 |
| left medial forebrain bundle | -0.0033 | -3.3619 | 1.28E-13 | 0.0539 |
| right Anterior area | -0.0649 | -3.3471 | 1.30E-13 | 0.0547 |
| right Dorsal part of the lateral geniculate complex | -0.0423 | -3.3377 | 1.33E-13 | 0.0549 |
| left Intergeniculate leaflet of the lateral geniculate complex | -0.0050 | -3.3266 | 1.31E-13 | 0.0549 |
| left Ventral medial nucleus of the thalamus | -0.0627 | -3.3256 | 1.32E-13 | 0.0549 |
| right Fields of Forel | -0.0174 | -3.3215 | 1.32E-13 | 0.0549 |
| left Submedial nucleus of the thalamus | -0.0200 | -3.3078 | 1.35E-13 | 0.0555 |
| left Cortical amygdalar area, anterior part | -0.0431 | -3.3014 | 1.38E-13 | 0.0555 |
| left Anterior hypothalamic nucleus | -0.0240 | -3.2874 | 1.34E-13 | 0.0555 |
| right Nucleus of the posterior commissure | -0.0141 | -3.2827 | 1.41E-13 | 0.0555 |
| Hypoglossal nucleus | -0.0282 | -3.2800 | 1.40E-13 | 0.0555 |
| left Perirhinal area | -0.0415 | -3.2746 | 1.37E-13 | 0.0555 |
| left Entorhinal area, medial part, dorsal zone | -0.3040 | -3.2746 | 1.36E-13 | 0.0555 |
| left optic radiation | -0.0783 | -3.2652 | 1.42E-13 | 0.0559 |
| left Prelimbic area | -0.1062 | -3.2273 | 1.43E-13 | 0.0587 |
| left Fields of Forel | -0.0158 | -3.2268 | 1.44E-13 | 0.0587 |
| right Prelimbic area | -0.1243 | -3.2044 | 1.45E-13 | 0.0605 |
| right Primary somatosensory area, unassigned | -0.0362 | -3.1929 | 1.46E-13 | 0.0612 |
| right Entorhinal area, lateral part | -0.2817 | -3.1875 | 1.47E-13 | 0.0612 |
| left Primary somatosensory area, upper limb | -0.1472 | -3.1774 | 1.48E-13 | 0.0618 |
| right lateral olfactory tract, body | -0.0541 | -3.1478 | 1.49E-13 | 0.0643 |

| | | | | |
|---|---|---|---|---|
| right Agranular insular area, dorsal part | -0.1797 | -3.1449 | 1.50E-13 | 0.0643 |
| left Secondary motor area | -0.4275 | -3.1288 | 1.53E-13 | 0.0645 |
| left nigrostriatal tract | -0.0058 | -3.1270 | 1.52E-13 | 0.0645 |
| right Gracile nucleus | -0.0096 | -3.1267 | 1.55E-13 | 0.0645 |
| left auditory radiation | -0.0160 | -3.1234 | 1.54E-13 | 0.0645 |
| left Nucleus of the brachium of the inferior colliculus | -0.0033 | -3.1106 | 1.56E-13 | 0.0654 |
| right Parafascicular nucleus | -0.0335 | -3.1012 | 1.59E-13 | 0.0655 |
| right Superior central nucleus raphe | -0.0240 | -3.1001 | 1.57E-13 | 0.0655 |
| left Zona incerta | -0.0784 | -3.0748 | 1.60E-13 | 0.0671 |
| right Basolateral amygdalar nucleus, anterior part | -0.0365 | -3.0735 | 1.61E-13 | 0.0671 |
| right Intercalated amygdalar nucleus | -0.0097 | -3.0721 | 1.62E-13 | 0.0671 |
| left Primary motor area | -0.3958 | -3.0605 | 1.64E-13 | 0.0680 |
| right Substantia nigra, compact part | -0.0136 | -3.0430 | 1.65E-13 | 0.0696 |
| right Subparafascicular nucleus | -0.0123 | -3.0309 | 1.66E-13 | 0.0705 |
| left Magnocellular nucleus | -0.0157 | -3.0187 | 1.67E-13 | 0.0712 |
| right Retrosplenial area, dorsal part | -0.1466 | -3.0143 | 1.69E-13 | 0.0712 |
| right Anteromedial nucleus, dorsal part | -0.0103 | -3.0123 | 1.70E-13 | 0.0712 |
| left Ventral tegmental area | -0.0258 | -3.0049 | 1.71E-13 | 0.0716 |
| left fasciculus retroflexus | -0.0089 | -2.9914 | 1.73E-13 | 0.0728 |
| left Piriform-amygdalar area | -0.0624 | -2.9798 | 1.77E-13 | 0.0732 |
| right Midbrain reticular nucleus | -0.3028 | -2.9794 | 1.78E-13 | 0.0732 |
| left Substantia innominata | -0.1218 | -2.9659 | 1.74E-13 | 0.0732 |
| right Cortical amygdalar area, posterior part, lateral zone | -0.0553 | -2.9651 | 1.80E-13 | 0.0732 |
| left Field CA1 | -0.3711 | -2.9636 | 1.75E-13 | 0.0732 |
| right Medullary reticular nucleus, ventral part | -0.0612 | -2.9627 | 1.81E-13 | 0.0732 |
| right Posterior limiting nucleus of the thalamus | -0.0117 | -2.9349 | 1.82E-13 | 0.0763 |
| right Magnocellular reticular nucleus | -0.0282 | -2.9285 | 1.84E-13 | 0.0767 |
| Edinger-Westphal nucleus | -0.0042 | -2.9239 | 1.85E-13 | 0.0767 |

| | | | | |
|---|---|---|---|---|
| right Primary somatosensory area, trunk | -0.0617 | -2.9059 | 1.87E-13 | 0.0787 |
| right posteromedial visual area | -0.0500 | -2.8985 | 1.88E-13 | 0.0792 |
| left Agranular insular area, posterior part | -0.1084 | -2.8929 | 1.90E-13 | 0.0794 |
| right Gustatory areas | -0.0740 | -2.8816 | 1.93E-13 | 0.0799 |
| left Posterior intralaminar thalamic nucleus | -0.0077 | -2.8791 | 1.91E-13 | 0.0799 |
| right Crus 2 | -0.3291 | -2.8777 | 1.94E-13 | 0.0799 |
| right Midbrain reticular nucleus, retrorubral area | -0.0087 | -2.8677 | 1.96E-13 | 0.0807 |
| right Pontine central gray | -0.0196 | -2.8464 | 1.97E-13 | 0.0833 |
| right Interstitial nucleus of Cajal | -0.0047 | -2.8382 | 1.99E-13 | 0.0840 |
| left Ectorhinal area | -0.0483 | -2.8248 | 2.01E-13 | 0.0854 |
| right Hippocampo-amygdalar transition area | -0.0200 | -2.8103 | 2.07E-13 | 0.0863 |
| left Anteroventral preoptic nucleus | -0.0039 | -2.8100 | 2.04E-13 | 0.0863 |
| left Anterior cingulate area, ventral part | -0.0888 | -2.8059 | 2.06E-13 | 0.0863 |
| left Arcuate hypothalamic nucleus | -0.0133 | -2.8038 | 2.02E-13 | 0.0863 |
| Periaqueductal gray | -0.2667 | -2.7944 | 2.12E-13 | 0.0870 |
| left Subparafascicular nucleus | -0.0115 | -2.7911 | 2.09E-13 | 0.0870 |
| left Medial preoptic nucleus | -0.0122 | -2.7884 | 2.11E-13 | 0.0870 |
| right Primary somatosensory area, lower limb | -0.0677 | -2.7850 | 2.14E-13 | 0.0870 |
| right Posterior amygdalar nucleus | -0.0457 | -2.7773 | 2.17E-13 | 0.0876 |
| left Spinal nucleus of the trigeminal, caudal part | -0.0995 | -2.7739 | 2.16E-13 | 0.0876 |
| left Anteromedial nucleus, dorsal part | -0.0103 | -2.7489 | 2.21E-13 | 0.0904 |
| left Postpiriform transition area | -0.0514 | -2.7486 | 2.19E-13 | 0.0904 |
| right Ventral part of the lateral geniculate complex | -0.0189 | -2.7430 | 2.23E-13 | 0.0904 |
| right Pontine gray | -0.0597 | -2.7426 | 2.25E-13 | 0.0904 |
| right Primary somatosensory area, upper limb | -0.0864 | -2.7280 | 2.27E-13 | 0.0918 |
| right mammillothalamic tract | -0.0077 | -2.7271 | 2.28E-13 | 0.0918 |
| right Entorhinal area, medial part, dorsal zone | -0.2488 | -2.7154 | 2.59E-13 | 0.0926 |
| left Orbital area, medial part | -0.0659 | -2.7042 | 2.38E-13 | 0.0926 |

| | | | | |
|---|---|---|---|---|
| left Ventral premammillary nucleus | -0.0111 | -2.7027 | 2.40E-13 | 0.0926 |
| left Paraventricular hypothalamic nucleus | -0.0092 | -2.7010 | 2.30E-13 | 0.0926 |
| left Linear nucleus of the medulla | -0.0046 | -2.6984 | 2.34E-13 | 0.0926 |
| right Anteroventral preoptic nucleus | -0.0038 | -2.6981 | 2.52E-13 | 0.0926 |
| right Ectorhinal area | -0.0533 | -2.6972 | 2.63E-13 | 0.0926 |
| left Posterior triangular thalamic nucleus | -0.0113 | -2.6817 | 2.48E-13 | 0.0926 |
| left Pedunculopontine nucleus | -0.0452 | -2.6776 | 2.44E-13 | 0.0926 |
| right Septohippocampal nucleus | -0.0027 | -2.6753 | 2.55E-13 | 0.0926 |
| left Primary somatosensory area, trunk | -0.0481 | -2.6752 | 2.42E-13 | 0.0926 |
| left Medial mammillary nucleus | -0.0350 | -2.6735 | 2.50E-13 | 0.0926 |
| right columns of the fornix | -0.0147 | -2.6734 | 2.57E-13 | 0.0926 |
| left Inferior olivary complex | -0.0331 | -2.6733 | 2.32E-13 | 0.0926 |
| left Medullary reticular nucleus, dorsal part | -0.0991 | -2.6697 | 2.46E-13 | 0.0926 |
| left Midbrain reticular nucleus, retrorubral area | -0.0109 | -2.6644 | 2.36E-13 | 0.0926 |
| right stria medullaris | -0.0119 | -2.6640 | 2.61E-13 | 0.0926 |
| right optic radiation | -0.0500 | -2.6640 | 2.65E-13 | 0.0926 |
| left brachium of the superior colliculus | -0.0113 | -2.6580 | 2.68E-13 | 0.0931 |
| left Dorsal nucleus raphe | -0.0067 | -2.6532 | 2.70E-13 | 0.0934 |
| right Paracentral nucleus | -0.0153 | -2.6450 | 2.72E-13 | 0.0942 |
| right rubrospinal tract | -0.0190 | -2.6378 | 2.79E-13 | 0.0946 |
| left internal capsule | -0.0696 | -2.6368 | 2.75E-13 | 0.0946 |
| right Substantia nigra, reticular part | -0.0935 | -2.6340 | 2.77E-13 | 0.0946 |
| left Primary somatosensory area, nose | -0.1022 | -2.6253 | 2.82E-13 | 0.0955 |
| left Dorsal premammillary nucleus | -0.0072 | -2.6196 | 2.84E-13 | 0.0960 |
| right Koelliker-Fuse subnucleus | -0.0094 | -2.6149 | 2.87E-13 | 0.0963 |
| right Basolateral amygdalar nucleus, ventral part | -0.0159 | -2.6009 | 2.97E-13 | 0.0978 |
| right Nucleus accumbens | -0.2116 | -2.5970 | 2.94E-13 | 0.0978 |
| left Midbrain reticular nucleus | -0.2580 | -2.5959 | 2.89E-13 | 0.0978 |
| left Medial amygdalar nucleus | -0.0815 | -2.5949 | 2.92E-13 | 0.0978 |
| corpus callosum | -0.4339 | -2.5908 | 2.99E-13 | 0.0980 |

| | | | | |
|---|---|---|---|---|
| left Dorsal part of the lateral geniculate complex | -0.0452 | -2.5852 | 3.02E-13 | 0.0985 |
| left Basolateral amygdalar nucleus, anterior part | -0.0373 | -2.5817 | 3.05E-13 | 0.0986 |
| left Fundus of striatum | -0.0185 | -2.5728 | 3.30E-13 | 0.0992 |
| left Temporal association areas | -0.0916 | -2.5697 | 3.13E-13 | 0.0992 |
| left Cortical subplate | -0.0121 | -2.5669 | 3.27E-13 | 0.0992 |
| left Nucleus of the lateral olfactory tract | -0.0154 | -2.5540 | 3.33E-13 | 0.0992 |
| left Supratrigeminal nucleus | -0.0126 | -2.5534 | 3.18E-13 | 0.0992 |
| right Anterior cingulate area, ventral part | -0.0952 | -2.5529 | 3.39E-13 | 0.0992 |
| Rostral linear nucleus raphe | -0.0081 | -2.5528 | 3.36E-13 | 0.0992 |
| left Oculomotor nucleus | -0.0029 | -2.5503 | 3.10E-13 | 0.0992 |
| right Ventral premammillary nucleus | -0.0086 | -2.5482 | 3.45E-13 | 0.0992 |
| left Nucleus of Darkschewitsch | -0.0055 | -2.5470 | 3.21E-13 | 0.0992 |
| left Superior central nucleus raphe | -0.0206 | -2.5424 | 3.24E-13 | 0.0992 |
| left Medial preoptic area | -0.0186 | -2.5415 | 3.15E-13 | 0.0992 |
| left Parataenial nucleus | -0.0127 | -2.5402 | 3.07E-13 | 0.0992 |
| right Retrosplenial area, lateral agranular part | -0.0954 | -2.5399 | 3.42E-13 | 0.0992 |
| left Pontine reticular nucleus, caudal part | -0.0794 | -2.5346 | 3.81E-13 | 0.0992 |
| left Lateral reticular nucleus, parvicellular part | -0.0046 | -2.5309 | 3.74E-13 | 0.0992 |
| right Anterior cingulate area | -0.1454 | -2.5288 | 4.10E-13 | 0.0992 |
| left Dorsal motor nucleus of the vagus nerve | -0.0153 | -2.5277 | 3.64E-13 | 0.0992 |
| right Lateral posterior nucleus of the thalamus | -0.0666 | -2.5170 | 3.98E-13 | 0.0992 |
| left Anterior cingulate area | -0.1064 | -2.5131 | 3.67E-13 | 0.0992 |
| posterior commissure | -0.0045 | -2.5111 | 3.84E-13 | 0.0992 |
| left Lateral habenula | -0.0138 | -2.5109 | 3.48E-13 | 0.0992 |
| left Dorsal auditory area | -0.0439 | -2.5092 | 3.54E-13 | 0.0992 |
| left posteromedial visual area | -0.0473 | -2.5084 | 3.60E-13 | 0.0992 |
| right Interposed nucleus | -0.0438 | -2.5083 | 3.91E-13 | 0.0992 |
| right Copula pyramidis | -0.1336 | -2.5057 | 4.13E-13 | 0.0992 |
| right Cortical amygdalar area, anterior part | -0.0444 | -2.5015 | 4.06E-13 | 0.0992 |

| | | | | |
|---|---|---|---|---|
| right Pontine reticular nucleus | -0.1012 | -2.4997 | 3.95E-13 | 0.0992 |
| left Piriform area | -0.4505 | -2.4997 | 3.70E-13 | 0.0992 |
| left Red nucleus | -0.0510 | -2.4982 | 3.51E-13 | 0.0992 |
| left mammillothalamic tract | -0.0057 | -2.4945 | 3.57E-13 | 0.0992 |
| Nucleus raphe obscurus | -0.0079 | -2.4929 | 3.88E-13 | 0.0992 |
| right Midbrain trigeminal nucleus | -0.0009 | -2.4906 | 4.02E-13 | 0.0992 |
| left Posterior complex of the thalamus | -0.0640 | -2.4890 | 3.77E-13 | 0.0992 |
| right Red nucleus | -0.0539 | -2.4826 | 4.17E-13 | 0.0999 |
| left medial longitudinal fascicle | -0.0069 | -2.4722 | 4.21E-13 | 0.1003 |
| right Ethmoid nucleus of the thalamus | -0.0114 | -2.4719 | 4.33E-13 | 0.1003 |
| right Postpiriform transition area | -0.0565 | -2.4714 | 4.25E-13 | 0.1003 |
| right Paramedian lobule | -0.2623 | -2.4709 | 4.29E-13 | 0.1003 |
| left Substantia nigra, compact part | -0.0105 | -2.4676 | 4.37E-13 | 0.1004 |
| right Medial terminal nucleus of the accessory optic tract | -0.0029 | -2.4647 | 4.41E-13 | 0.1004 |
| right alveus | -0.0332 | -2.4626 | 4.50E-13 | 0.1004 |
| right Nucleus sagulum | -0.0030 | -2.4609 | 4.45E-13 | 0.1004 |
| left Retrochiasmatic area | -0.0073 | -2.4536 | 4.54E-13 | 0.1010 |
| left Subgeniculate nucleus | -0.0013 | -2.4530 | 4.58E-13 | 0.1010 |
| right Subgeniculate nucleus | -0.0016 | -2.4492 | 4.63E-13 | 0.1012 |
| left Dorsal tegmental nucleus | -0.0042 | -2.4430 | 4.71E-13 | 0.1018 |
| right Superior colliculus | -0.2495 | -2.4388 | 4.76E-13 | 0.1018 |
| left Parabrachial nucleus | -0.0455 | -2.4370 | 4.67E-13 | 0.1018 |
| right dorsal limb | -0.0116 | -2.4368 | 4.80E-13 | 0.1018 |
| right Primary visual area | -0.2216 | -2.4303 | 4.85E-13 | 0.1024 |
| right Medial mammillary nucleus | -0.0315 | -2.4292 | 4.90E-13 | 0.1024 |
| Induseum griseum | -0.0127 | -2.4243 | 5.04E-13 | 0.1025 |
| left Medial geniculate complex, medial part | -0.0100 | -2.4218 | 4.99E-13 | 0.1025 |
| left Medial vestibular nucleus | -0.0537 | -2.4217 | 4.94E-13 | 0.1025 |
| right Parapyramidal nucleus | -0.0038 | -2.4090 | 5.09E-13 | 0.1044 |
| left Posterior auditory area | -0.0197 | -2.4055 | 5.14E-13 | 0.1047 |
| left Pontine central gray | -0.0181 | -2.4030 | 5.19E-13 | 0.1047 |

| | | | | |
|---|---|---|---|---|
| right cerebal peduncle | -0.0455 | -2.3987 | 5.29E-13 | 0.1049 |
| left Primary visual area | -0.2197 | -2.3978 | 5.24E-13 | 0.1049 |
| right Suprageniculate nucleus | -0.0077 | -2.3943 | 5.34E-13 | 0.1049 |
| right Posterior auditory area | -0.0192 | -2.3933 | 5.39E-13 | 0.1049 |
| left Preparasubthalamic nucleus | -0.0011 | -2.3881 | 5.45E-13 | 0.1054 |
| right Globus pallidus, external segment | -0.0431 | -2.3822 | 5.61E-13 | 0.1054 |
| right Medullary reticular nucleus, dorsal part | -0.0510 | -2.3804 | 5.67E-13 | 0.1054 |
| ventral tegmental decussation | -0.0045 | -2.3803 | 5.50E-13 | 0.1054 |
| right Anterior tegmental nucleus | -0.0020 | -2.3799 | 5.56E-13 | 0.1054 |
| right Perirhinal area | -0.0238 | -2.3752 | 5.72E-13 | 0.1059 |
| left Dentate gyrus | -0.2059 | -2.3581 | 5.78E-13 | 0.1087 |
| right Parataenial nucleus | -0.0118 | -2.3559 | 5.84E-13 | 0.1087 |
| right principal mammillary tract | -0.0031 | -2.3474 | 6.07E-13 | 0.1094 |
| right Submedial nucleus of the thalamus | -0.0196 | -2.3467 | 6.01E-13 | 0.1094 |
| left Primary auditory area | -0.0719 | -2.3447 | 5.95E-13 | 0.1094 |
| left Motor nucleus of trigeminal | -0.0142 | -2.3439 | 5.89E-13 | 0.1094 |
| right Paranigral nucleus | -0.0021 | -2.3415 | 6.13E-13 | 0.1095 |
| left Medial geniculate complex, ventral part | -0.0093 | -2.3371 | 6.26E-13 | 0.1097 |
| Rhomboid nucleus | -0.0093 | -2.3355 | 6.39E-13 | 0.1097 |
| fourth ventricle | -0.0515 | -2.3332 | 6.32E-13 | 0.1097 |
| left Nucleus accumbens | -0.1866 | -2.3321 | 6.20E-13 | 0.1097 |
| right fasciculus retroflexus | -0.0075 | -2.3257 | 6.52E-13 | 0.1102 |
| right Bed nuclei of the stria terminalis | -0.0491 | -2.3252 | 6.45E-13 | 0.1102 |
| right Supramammillary nucleus | -0.0098 | -2.3228 | 6.58E-13 | 0.1103 |
| right Mediodorsal nucleus of thalamus | -0.0413 | -2.3169 | 6.79E-13 | 0.1109 |
| left Lateral visual area | -0.0495 | -2.3153 | 6.65E-13 | 0.1109 |
| left Tuberal nucleus | -0.0208 | -2.3140 | 6.72E-13 | 0.1109 |
| left Nucleus x | -0.0039 | -2.3116 | 6.86E-13 | 0.1109 |
| right Dentate nucleus | -0.0121 | -2.3075 | 6.93E-13 | 0.1113 |
| left Supramammillary nucleus | -0.0137 | -2.3041 | 7.00E-13 | 0.1116 |

| | | | | |
|---|---|---|---|---|
| left Primary somatosensory area, lower limb | -0.0598 | -2.3007 | 7.15E-13 | 0.1119 |
| left motor root of the trigeminal nerve | -0.0031 | -2.2953 | 7.07E-13 | 0.1119 |
| right Frontal pole | -0.0339 | -2.2941 | 7.30E-13 | 0.1119 |
| cerebral aqueduct | -0.1024 | -2.2926 | 7.22E-13 | 0.1119 |
| right Lateral visual area | -0.0393 | -2.2916 | 7.37E-13 | 0.1119 |
| right Dorsomedial nucleus of the hypothalamus | -0.0179 | -2.2910 | 7.45E-13 | 0.1119 |
| left Basolateral amygdalar nucleus, posterior part | -0.0229 | -2.2833 | 7.53E-13 | 0.1128 |
| right Medial preoptic nucleus | -0.0127 | -2.2823 | 7.61E-13 | 0.1128 |
| right Posterior intralaminar thalamic nucleus | -0.0081 | -2.2777 | 7.85E-13 | 0.1130 |
| right Peripeduncular nucleus | -0.0029 | -2.2777 | 7.77E-13 | 0.1130 |
| right superior cerebelar peduncles | -0.0235 | -2.2758 | 7.69E-13 | 0.1130 |
| left Anterior pretectal nucleus | -0.0477 | -2.2690 | 8.02E-13 | 0.1137 |
| right Lateral amygdalar nucleus | -0.0268 | -2.2656 | 8.19E-13 | 0.1137 |
| left Pontine reticular nucleus | -0.0954 | -2.2650 | 7.93E-13 | 0.1137 |
| left Intercalated amygdalar nucleus | -0.0064 | -2.2646 | 8.10E-13 | 0.1137 |
| right Motor nucleus of trigeminal | -0.0154 | -2.2514 | 8.36E-13 | 0.1157 |
| left Subthalamic nucleus | -0.0086 | -2.2506 | 8.28E-13 | 0.1157 |
| left mammillary peduncle | -0.0025 | -2.2451 | 8.54E-13 | 0.1160 |
| left Supraoculomotor periaqueductal gray | -0.0017 | -2.2422 | 8.82E-13 | 0.1160 |
| right Laterointermediate area | -0.0152 | -2.2414 | 9.63E-13 | 0.1160 |
| right Crus 1 | -0.3787 | -2.2382 | 9.42E-13 | 0.1160 |
| left Postrhinal area | -0.0752 | -2.2362 | 8.73E-13 | 0.1160 |
| right Inferior colliculus | -0.1524 | -2.2362 | 9.21E-13 | 0.1160 |
| Nucleus raphe magnus | -0.0171 | -2.2356 | 8.92E-13 | 0.1160 |
| left Paracentral nucleus | -0.0132 | -2.2307 | 8.64E-13 | 0.1160 |
| right Parastrial nucleus | -0.0034 | -2.2306 | 9.52E-13 | 0.1160 |
| left Field CA3 | -0.1412 | -2.2302 | 8.45E-13 | 0.1160 |
| Uvula (IX) | -0.3034 | -2.2289 | 9.02E-13 | 0.1160 |
| right Taenia tecta, dorsal part | -0.0304 | -2.2269 | 9.12E-13 | 0.1160 |
| right brachium of the superior colliculus | -0.0078 | -2.2255 | 9.32E-13 | 0.1160 |

| | | | | |
|---|---|---|---|---|
| right Posterior triangular thalamic nucleus | -0.0115 | -2.2219 | 9.73E-13 | 0.1163 |
| right Medial habenula | -0.0106 | -2.2156 | 9.84E-13 | 0.1173 |
| left Anteromedial visual area | -0.0456 | -2.2041 | 9.95E-13 | 0.1193 |
| Pyramus (VIII) | -0.1544 | -2.1987 | 1.01E-12 | 0.1199 |
| right Agranular insular area, ventral part | -0.0775 | -2.1973 | 1.02E-12 | 0.1199 |
| right Basomedial amygdalar nucleus, anterior part | -0.0278 | -2.1945 | 1.04E-12 | 0.1201 |
| right Lateral dorsal nucleus of thalamus | -0.0320 | -2.1929 | 1.03E-12 | 0.1201 |
| left crossed tectospinal pathway | -0.0195 | -2.1905 | 1.05E-12 | 0.1202 |
| Median eminence | -0.0042 | -2.1884 | 1.06E-12 | 0.1203 |
| right Medial accesory oculomotor nucleus | -0.0020 | -2.1864 | 1.08E-12 | 0.1203 |
| left Ventral auditory area | -0.0539 | -2.1804 | 1.09E-12 | 0.1212 |
| left Lateral dorsal nucleus of thalamus | -0.0348 | -2.1650 | 1.10E-12 | 0.1242 |
| left Anteromedial nucleus, ventral part | -0.0064 | -2.1533 | 1.11E-12 | 0.1263 |
| left Parafascicular nucleus | -0.0269 | -2.1460 | 1.13E-12 | 0.1276 |
| right Pontine reticular nucleus, caudal part | -0.0852 | -2.1412 | 1.14E-12 | 0.1283 |
| right Gigantocellular reticular nucleus | -0.0941 | -2.1354 | 1.19E-12 | 0.1288 |
| left Nucleus of the solitary tract | -0.0429 | -2.1341 | 1.16E-12 | 0.1288 |
| left Anterodorsal preoptic nucleus | -0.0028 | -2.1321 | 1.15E-12 | 0.1288 |
| fiber tracts | -0.0897 | -2.1318 | 1.18E-12 | 0.1288 |
| right Dorsal peduncular area | -0.0178 | -2.1259 | 1.22E-12 | 0.1297 |
| left Nucleus of reuniens | -0.0185 | -2.1245 | 1.20E-12 | 0.1297 |
| right motor root of the trigeminal nerve | -0.0019 | -2.1188 | 1.25E-12 | 0.1306 |
| left Olfactory tubercle | -0.1123 | -2.1157 | 1.23E-12 | 0.1306 |
| right Paraventricular nucleus of the thalamus | -0.0202 | -2.1145 | 1.26E-12 | 0.1306 |
| right dorsal spinocerebellar tract | -0.0068 | -2.1137 | 1.28E-12 | 0.1306 |
| right Piriform-amygdalar area | -0.0677 | -2.1082 | 1.29E-12 | 0.1314 |
| left Nucleus of the optic tract | -0.0044 | -2.1031 | 1.32E-12 | 0.1317 |
| left Pontine gray | -0.0549 | -2.1028 | 1.34E-12 | 0.1317 |
| left Anterior amygdalar area | -0.0133 | -2.1018 | 1.31E-12 | 0.1317 |

| | | | | |
|---|---|---|---|---|
| right Lateral habenula | -0.0141 | -2.0966 | 1.35E-12 | 0.1326 |
| left medial lemniscus | -0.0231 | -2.0946 | 1.37E-12 | 0.1326 |
| left superior cerebelar peduncles | -0.0196 | -2.0886 | 1.38E-12 | 0.1331 |
| left Nucleus of the posterior commissure | -0.0087 | -2.0879 | 1.40E-12 | 0.1331 |
| left Caudoputamen | -0.8220 | -2.0864 | 1.42E-12 | 0.1331 |
| right Intergeniculate leaflet of the lateral geniculate complex | -0.0033 | -2.0858 | 1.44E-12 | 0.1331 |
| right Ventral tegmental area | -0.0212 | -2.0820 | 1.45E-12 | 0.1336 |
| right Anterodorsal preoptic nucleus | -0.0041 | -2.0756 | 1.47E-12 | 0.1347 |
| right Anterior pretectal nucleus | -0.0391 | -2.0731 | 1.49E-12 | 0.1349 |
| right Nucleus of the lateral olfactory tract | -0.0146 | -2.0715 | 1.51E-12 | 0.1349 |
| right corticospinal tract | -0.0052 | -2.0673 | 1.52E-12 | 0.1355 |
| right Main olfactory bulb | -0.4727 | -2.0652 | 1.54E-12 | 0.1356 |
| left Interanterodorsal nucleus of the thalamus | -0.0056 | -2.0621 | 1.56E-12 | 0.1360 |
| right Principal sensory nucleus of the trigeminal | -0.0290 | -2.0581 | 1.58E-12 | 0.1365 |
| right Perifornical nucleus | -0.0085 | -2.0454 | 1.60E-12 | 0.1392 |
| right Dorsal tegmental nucleus | -0.0031 | -2.0413 | 1.62E-12 | 0.1398 |
| right Bed nucleus of the anterior commissure | -0.0009 | -2.0362 | 1.66E-12 | 0.1403 |
| left Paragigantocellular reticular nucleus, dorsal part | -0.0057 | -2.0359 | 1.64E-12 | 0.1403 |
| right Substantia innominata | -0.1053 | -2.0342 | 1.68E-12 | 0.1403 |
| right Supratrigeminal nucleus | -0.0103 | -2.0268 | 1.70E-12 | 0.1417 |
| right Cuneiform nucleus | -0.0232 | -2.0235 | 1.72E-12 | 0.1421 |
| Declive (VI) | -0.2147 | -2.0164 | 1.76E-12 | 0.1429 |
| right Dentate gyrus | -0.1457 | -2.0161 | 1.79E-12 | 0.1429 |
| right Ventromedial hypothalamic nucleus | -0.0163 | -2.0156 | 1.81E-12 | 0.1429 |
| left Substantia nigra, reticular part | -0.0653 | -2.0141 | 1.74E-12 | 0.1429 |
| right ventral spinocerebellar tract | -0.0051 | -2.0085 | 1.85E-12 | 0.1436 |
| left cerebal peduncle | -0.0429 | -2.0079 | 1.83E-12 | 0.1436 |
| left Bed nuclei of the stria terminalis | -0.0449 | -2.0046 | 1.88E-12 | 0.1438 |
| left Posterolateral visual area | -0.0418 | -2.0042 | 1.90E-12 | 0.1438 |

| | | | | |
|---|---|---|---|---|
| left Tegmental reticular nucleus | -0.0238 | -1.9968 | 1.95E-12 | 0.1451 |
| left Infracerebellar nucleus | -0.0056 | -1.9958 | 1.92E-12 | 0.1451 |
| right Medial amygdalar nucleus | -0.0799 | -1.9908 | 1.97E-12 | 0.1458 |
| right Tegmental reticular nucleus | -0.0267 | -1.9899 | 2.00E-12 | 0.1458 |
| left Peritrigeminal zone | -0.0110 | -1.9831 | 2.02E-12 | 0.1471 |
| left Supplemental somatosensory area | -0.1954 | -1.9798 | 2.05E-12 | 0.1475 |
| third ventricle | -0.0541 | -1.9727 | 2.08E-12 | 0.1490 |
| left Septohippocampal nucleus | -0.0021 | -1.9697 | 2.10E-12 | 0.1493 |
| left Mediodorsal nucleus of thalamus | -0.0384 | -1.9590 | 2.13E-12 | 0.1514 |
| left lateral olfactory tract, body | -0.0361 | -1.9586 | 2.16E-12 | 0.1514 |
| right Simple lobule | -0.2859 | -1.9562 | 2.19E-12 | 0.1516 |
| left Anteroventral nucleus of thalamus | -0.0163 | -1.9513 | 2.21E-12 | 0.1522 |
| right Subparaventricular zone | -0.0038 | -1.9509 | 2.24E-12 | 0.1522 |
| left Inferior colliculus | -0.1741 | -1.9464 | 2.27E-12 | 0.1530 |
| left Orbital area, ventrolateral part | -0.0605 | -1.9368 | 2.30E-12 | 0.1551 |
| left Spinal vestibular nucleus | -0.0268 | -1.9338 | 2.33E-12 | 0.1555 |
| left Gracile nucleus | -0.0076 | -1.9296 | 2.36E-12 | 0.1562 |
| left Medial terminal nucleus of the accessory optic tract | -0.0025 | -1.9261 | 2.40E-12 | 0.1567 |
| left Lateral posterior nucleus of the thalamus | -0.0501 | -1.9249 | 2.43E-12 | 0.1567 |
| left Periventricular hypothalamic nucleus | -0.0242 | -1.9225 | 2.46E-12 | 0.1569 |
| right Superior vestibular nucleus | -0.0169 | -1.9187 | 2.49E-12 | 0.1575 |
| right uncinate fascicle | -0.0039 | -1.9148 | 2.56E-12 | 0.1579 |
| left Cuneiform nucleus | -0.0221 | -1.9143 | 2.53E-12 | 0.1579 |
| right Subparafascicular area | -0.0057 | -1.9049 | 2.59E-12 | 0.1600 |
| left Crus 2 | -0.2290 | -1.8985 | 2.63E-12 | 0.1614 |
| right Nucleus of the lateral lemniscus | -0.0189 | -1.8950 | 2.66E-12 | 0.1619 |
| left Agranular insular area, dorsal part | -0.0903 | -1.8922 | 2.70E-12 | 0.1623 |
| left Laterodorsal tegmental nucleus | -0.0061 | -1.8904 | 2.74E-12 | 0.1624 |
| right Dorsal premammillary nucleus | -0.0047 | -1.8859 | 2.77E-12 | 0.1632 |
| left Retroparafascicular nucleus | -0.0026 | -1.8835 | 2.81E-12 | 0.1635 |

| | | | | |
|---|---|---|---|---|
| right Hippocampal formation | -0.0121 | -1.8788 | 2.89E-12 | 0.1643 |
| left columns of the fornix | -0.0116 | -1.8774 | 2.85E-12 | 0.1643 |
| right crossed tectospinal pathway | -0.0205 | -1.8748 | 2.93E-12 | 0.1647 |
| right Medial preoptic area | -0.0145 | -1.8716 | 2.97E-12 | 0.1652 |
| right nigrostriatal tract | -0.0045 | -1.8620 | 3.01E-12 | 0.1674 |
| left Gigantocellular reticular nucleus | -0.0754 | -1.8496 | 3.05E-12 | 0.1705 |
| right Endopiriform nucleus, dorsal part | -0.0640 | -1.8470 | 3.18E-12 | 0.1708 |
| Intermediodorsal nucleus of the thalamus | -0.0114 | -1.8440 | 3.10E-12 | 0.1708 |
| right Cuneate nucleus | -0.0181 | -1.8440 | 3.14E-12 | 0.1708 |
| right Medial geniculate complex, dorsal part | -0.0045 | -1.8429 | 3.23E-12 | 0.1708 |
| left Anterolateral visual area | -0.0239 | -1.8401 | 3.27E-12 | 0.1712 |
| right Piriform area | -0.4636 | -1.8383 | 3.32E-12 | 0.1713 |
| left Interstitial nucleus of Cajal | -0.0027 | -1.8339 | 3.37E-12 | 0.1722 |
| right subependymal zone | -0.0065 | -1.8252 | 3.46E-12 | 0.1738 |
| right Arcuate hypothalamic nucleus | -0.0079 | -1.8248 | 3.51E-12 | 0.1738 |
| right Periventricular hypothalamic nucleus | -0.0196 | -1.8240 | 3.41E-12 | 0.1738 |
| left Retrosplenial area, lateral agranular part | -0.0663 | -1.8217 | 3.56E-12 | 0.1740 |
| right cingulum bundle | -0.0322 | -1.8192 | 3.67E-12 | 0.1741 |
| right pyramid | -0.0108 | -1.8188 | 3.61E-12 | 0.1741 |
| right Anterolateral visual area | -0.0243 | -1.8138 | 3.72E-12 | 0.1751 |
| pyramidal decussation | -0.0134 | -1.8114 | 3.88E-12 | 0.1751 |
| left Suprachiasmatic nucleus | -0.0018 | -1.8097 | 3.77E-12 | 0.1751 |
| left Primary somatosensory area, mouth | -0.1359 | -1.8095 | 3.83E-12 | 0.1751 |
| left dorsal acoustic stria | -0.0009 | -1.8041 | 3.94E-12 | 0.1763 |
| right Tuberomammillary nucleus, dorsal part | -0.0018 | -1.7924 | 4.00E-12 | 0.1793 |
| right Taenia tecta, ventral part | -0.0179 | -1.7839 | 4.06E-12 | 0.1815 |
| right medial lemniscus | -0.0184 | -1.7627 | 4.12E-12 | 0.1876 |
| left brachium of the inferior colliculus | -0.0073 | -1.7609 | 4.18E-12 | 0.1877 |
| left Lateral reticular nucleus, magnocellular part | -0.0196 | -1.7590 | 4.24E-12 | 0.1878 |

| | | | | |
|---|---|---|---|---|
| right Endopiriform nucleus, ventral part | -0.0266 | -1.7576 | 4.30E-12 | 0.1878 |
| left Visceral area | -0.0533 | -1.7521 | 4.37E-12 | 0.1891 |
| left Ethmoid nucleus of the thalamus | -0.0108 | -1.7446 | 4.43E-12 | 0.1910 |
| left Orbital area, lateral part | -0.0750 | -1.7410 | 4.50E-12 | 0.1917 |
| right Parabigeminal nucleus | -0.0020 | -1.7356 | 4.57E-12 | 0.1930 |
| right Anterior amygdalar area | -0.0156 | -1.7292 | 4.63E-12 | 0.1947 |
| left external medullary lamina of the thalamus | -0.0024 | -1.7271 | 4.71E-12 | 0.1949 |
| left Claustrum | -0.0126 | -1.7255 | 4.78E-12 | 0.1949 |
| right Anterior olfactory nucleus | -0.1143 | -1.7160 | 4.85E-12 | 0.1976 |
| left Paraventricular nucleus of the thalamus | -0.0194 | -1.7094 | 4.92E-12 | 0.1993 |
| Vascular organ of the lamina terminalis | -0.0010 | -1.6920 | 5.08E-12 | 0.2041 |
| left Superior colliculus | -0.2001 | -1.6915 | 5.00E-12 | 0.2041 |
| right optic tract | -0.0217 | -1.6908 | 5.15E-12 | 0.2041 |
| right Central medial nucleus of the thalamus | -0.0069 | -1.6713 | 5.23E-12 | 0.2102 |
| left Agranular insular area, ventral part | -0.0491 | -1.6689 | 5.32E-12 | 0.2106 |
| left mammillotegmental tract | -0.0027 | -1.6626 | 5.40E-12 | 0.2123 |
| left Laterointermediate area | -0.0166 | -1.6421 | 5.48E-12 | 0.2186 |
| right Dorsal motor nucleus of the vagus nerve | -0.0123 | -1.6418 | 5.57E-12 | 0.2186 |
| right Anterior hypothalamic nucleus | -0.0182 | -1.6392 | 5.66E-12 | 0.2190 |
| right Vestibulocerebellar nucleus | -0.0045 | -1.6366 | 5.75E-12 | 0.2195 |
| left Koelliker-Fuse subnucleus | -0.0062 | -1.6331 | 5.84E-12 | 0.2202 |
| left Central lateral nucleus of the thalamus | -0.0148 | -1.6297 | 5.93E-12 | 0.2209 |
| right lateral lemniscus | -0.0232 | -1.6286 | 6.02E-12 | 0.2209 |
| right Infracerebellar nucleus | -0.0036 | -1.6246 | 6.12E-12 | 0.2218 |
| right oculomotor nerve | -0.0007 | -1.6228 | 6.22E-12 | 0.2220 |
| left Medial habenula | -0.0111 | -1.6204 | 6.32E-12 | 0.2224 |
| left Diagonal band nucleus | -0.0185 | -1.6158 | 6.42E-12 | 0.2236 |
| Medial septal nucleus | -0.0274 | -1.6089 | 6.52E-12 | 0.2256 |
| right Nucleus of the brachium of the inferior colliculus | -0.0031 | -1.6038 | 6.63E-12 | 0.2270 |

| | | | | |
|---|---|---|---|---|
| right Cortical subplate | -0.0070 | -1.6017 | 6.85E-12 | 0.2271 |
| left Principal sensory nucleus of the trigeminal | -0.0208 | -1.6008 | 6.74E-12 | 0.2271 |
| right Perireunensis nucleus | -0.0056 | -1.5917 | 6.96E-12 | 0.2299 |
| left stria medullaris | -0.0079 | -1.5758 | 7.07E-12 | 0.2354 |
| left Paratrigeminal nucleus | -0.0059 | -1.5732 | 7.19E-12 | 0.2358 |
| right mammillotegmental tract | -0.0032 | -1.5651 | 7.31E-12 | 0.2384 |
| left Perifornical nucleus | -0.0060 | -1.5608 | 7.43E-12 | 0.2396 |
| right Nucleus of reuniens | -0.0158 | -1.5595 | 7.55E-12 | 0.2396 |
| right brachium of the inferior colliculus | -0.0078 | -1.5541 | 7.68E-12 | 0.2411 |
| right Anteroventral nucleus of thalamus | -0.0155 | -1.5504 | 7.81E-12 | 0.2420 |
| right Nucleus of Roller | -0.0025 | -1.5487 | 8.08E-12 | 0.2422 |
| left Interposed nucleus | -0.0328 | -1.5474 | 7.94E-12 | 0.2422 |
| left Dorsal peduncular area | -0.0144 | -1.5423 | 8.21E-12 | 0.2436 |
| right Postrhinal area | -0.0407 | -1.5348 | 8.49E-12 | 0.2455 |
| Central linear nucleus raphe | -0.0070 | -1.5347 | 8.35E-12 | 0.2455 |
| right Nucleus ambiguus, dorsal division | -0.0012 | -1.5314 | 8.64E-12 | 0.2463 |
| left Dentate nucleus | -0.0097 | -1.5250 | 8.94E-12 | 0.2478 |
| anterior commissure | -0.0478 | -1.5218 | 9.09E-12 | 0.2483 |
| right Paragigantocellular reticular nucleus, lateral part | -0.0258 | -1.5212 | 9.25E-12 | 0.2483 |
| left supra-callosal cerebral white matter | -0.0193 | -1.5130 | 9.41E-12 | 0.2510 |
| left Medullary reticular nucleus, ventral part | -0.0514 | -1.5107 | 9.58E-12 | 0.2514 |
| left Subceruleus nucleus | -0.0009 | -1.5054 | 9.75E-12 | 0.2526 |
| left Central medial nucleus of the thalamus | -0.0068 | -1.5049 | 9.92E-12 | 0.2526 |
| right Interanterodorsal nucleus of the thalamus | -0.0038 | -1.4958 | 1.01E-11 | 0.2558 |
| left Retrosplenial area, dorsal part | -0.0714 | -1.4943 | 1.03E-11 | 0.2559 |
| right Anterodorsal nucleus | -0.0050 | -1.4889 | 1.08E-11 | 0.2564 |
| right Lingula (I) | -0.0058 | -1.4882 | 1.10E-11 | 0.2564 |
| Subcommissural organ | -0.0010 | -1.4881 | 1.06E-11 | 0.2564 |
| left Paramedian lobule | -0.1463 | -1.4877 | 1.05E-11 | 0.2564 |

| | | | | |
|---|---|---|---|---|
| right external medullary lamina of the thalamus | -0.0035 | -1.4821 | 1.14E-11 | 0.2580 |
| left Suprageniculate nucleus | -0.0049 | -1.4812 | 1.12E-11 | 0.2580 |
| right sensory root of the trigeminal nerve | -0.0149 | -1.4785 | 1.16E-11 | 0.2586 |
| Folium-tuber vermis (VII) | -0.1077 | -1.4751 | 1.18E-11 | 0.2595 |
| left Bed nucleus of the anterior commissure | -0.0004 | -1.4582 | 1.21E-11 | 0.2659 |
| left Crus 1 | -0.2404 | -1.4489 | 1.23E-11 | 0.2692 |
| doral tegmental decussation | -0.0009 | -1.4467 | 1.30E-11 | 0.2696 |
| left Superior vestibular nucleus | -0.0153 | -1.4453 | 1.25E-11 | 0.2696 |
| left Posterior limiting nucleus of the thalamus | -0.0064 | -1.4443 | 1.27E-11 | 0.2696 |
| right Diagonal band nucleus | -0.0174 | -1.4402 | 1.35E-11 | 0.2703 |
| left Parvicellular motor 5 nucleus | -0.0027 | -1.4399 | 1.32E-11 | 0.2703 |
| left Intermediate geniculate nucleus | -0.0010 | -1.4294 | 1.43E-11 | 0.2735 |
| right Area postrema | -0.0033 | -1.4289 | 1.45E-11 | 0.2735 |
| left Cuneate nucleus | -0.0161 | -1.4281 | 1.37E-11 | 0.2735 |
| right supra-callosal cerebral white matter | -0.0180 | -1.4271 | 1.48E-11 | 0.2735 |
| left corticospinal tract | -0.0039 | -1.4259 | 1.40E-11 | 0.2735 |
| right Superior olivary complex, periolivary region | -0.0064 | -1.4245 | 1.51E-11 | 0.2736 |
| right Inferior salivatory nucleus | -0.0007 | -1.4116 | 1.54E-11 | 0.2786 |
| right Central lateral nucleus of the thalamus | -0.0136 | -1.3965 | 1.63E-11 | 0.2841 |
| right stria terminalis | -0.0121 | -1.3961 | 1.60E-11 | 0.2841 |
| right Accessory olfactory bulb | -0.0196 | -1.3952 | 1.57E-11 | 0.2841 |
| right Retroparafascicular nucleus | -0.0021 | -1.3906 | 1.66E-11 | 0.2855 |
| right Anteroventral periventricular nucleus | -0.0040 | -1.3883 | 1.69E-11 | 0.2860 |
| left Supragenual nucleus | -0.0009 | -1.3869 | 1.73E-11 | 0.2860 |
| right Fundus of striatum | -0.0148 | -1.3802 | 1.79E-11 | 0.2883 |
| right Suprachiasmatic nucleus | -0.0011 | -1.3793 | 1.76E-11 | 0.2883 |
| right Lateral reticular nucleus, magnocellular part | -0.0149 | -1.3776 | 1.83E-11 | 0.2885 |
| left alveus | -0.0364 | -1.3754 | 1.87E-11 | 0.2889 |
| right Posterior pretectal nucleus | -0.0033 | -1.3491 | 1.90E-11 | 0.3001 |

| | | | | |
|---|---|---|---|---|
| right Olfactory tubercle | -0.1184 | -1.3437 | 1.98E-11 | 0.3017 |
| left Perireunensis nucleus | -0.0035 | -1.3432 | 1.94E-11 | 0.3017 |
| right Parasubiculum | -0.0274 | -1.3408 | 2.02E-11 | 0.3022 |
| right Preparasubthalamic nucleus | -0.0006 | -1.3360 | 2.06E-11 | 0.3039 |
| left Posterodorsal preoptic nucleus | -0.0004 | -1.3306 | 2.11E-11 | 0.3058 |
| Nodulus (X) | -0.1187 | -1.3284 | 2.15E-11 | 0.3062 |
| right Dorsal nucleus raphe | -0.0033 | -1.3224 | 2.19E-11 | 0.3084 |
| right Paraventricular hypothalamic nucleus | -0.0093 | -1.3147 | 2.24E-11 | 0.3114 |
| right Superior olivary complex, lateral part | -0.0111 | -1.3126 | 2.29E-11 | 0.3118 |
| Lobule II | -0.1895 | -1.3019 | 2.33E-11 | 0.3163 |
| left commissural branch of stria terminalis | -0.0009 | -1.2937 | 2.38E-11 | 0.3196 |
| left Lateral amygdalar nucleus | -0.0301 | -1.2899 | 2.43E-11 | 0.3204 |
| superior cerebellar peduncle decussation | -0.0019 | -1.2895 | 2.49E-11 | 0.3204 |
| right middle cerebellar peduncle | -0.0237 | -1.2862 | 2.54E-11 | 0.3214 |
| left Nucleus prepositus | -0.0075 | -1.2814 | 2.59E-11 | 0.3231 |
| root | -0.1784 | -1.2695 | 2.70E-11 | 0.3282 |
| left Intermediate reticular nucleus | -0.0611 | -1.2684 | 2.65E-11 | 0.3282 |
| left choroid plexus | -0.0450 | -1.2663 | 2.76E-11 | 0.3287 |
| right Retrosplenial area, ventral part | -0.0897 | -1.2641 | 2.82E-11 | 0.3291 |
| left Parasolitary nucleus | -0.0021 | -1.2574 | 2.95E-11 | 0.3314 |
| left Hippocampal formation | -0.0113 | -1.2569 | 3.01E-11 | 0.3314 |
| left stria terminalis | -0.0105 | -1.2546 | 2.89E-11 | 0.3314 |
| left Supraoptic nucleus | -0.0021 | -1.2503 | 3.15E-11 | 0.3329 |
| left Tuberomammillary nucleus, dorsal part | -0.0017 | -1.2358 | 3.22E-11 | 0.3388 |
| right Spinal nucleus of the trigeminal, caudal part | -0.0453 | -1.2358 | 3.29E-11 | 0.3388 |
| left Taenia tecta, ventral part | -0.0157 | -1.2289 | 3.36E-11 | 0.3417 |
| right Flocculus | -0.0328 | -1.2104 | 3.44E-11 | 0.3504 |
| left Area prostriata | -0.0092 | -1.2070 | 3.52E-11 | 0.3516 |
| left Locus ceruleus | -0.0004 | -1.1928 | 3.60E-11 | 0.3583 |
| right amygdalar capsule | -0.0039 | -1.1774 | 3.77E-11 | 0.3656 |

| | | | | |
|---|---|---|---|---|
| left Nucleus ambiguus, ventral division | -0.0019 | -1.1763 | 3.68E-11 | 0.3656 |
| right Xiphoid thalamic nucleus | -0.0041 | -1.1704 | 3.85E-11 | 0.3681 |
| right Presubiculum | -0.0203 | -1.1643 | 3.94E-11 | 0.3707 |
| left Parvicellular reticular nucleus | -0.0500 | -1.1616 | 4.04E-11 | 0.3715 |
| dorsal hippocampal commissure | -0.0621 | -1.1567 | 4.13E-11 | 0.3734 |
| right Nucleus of the optic tract | -0.0041 | -1.1532 | 4.23E-11 | 0.3746 |
| arbor vitae | -0.4318 | -1.1509 | 4.33E-11 | 0.3752 |
| left Lateral septal nucleus, caudal (caudodorsal) part | -0.0203 | -1.1443 | 4.43E-11 | 0.3781 |
| left Nucleus of Roller | -0.0014 | -1.1133 | 4.54E-11 | 0.3941 |
| right Medial vestibular nucleus | -0.0317 | -1.1128 | 4.76E-11 | 0.3941 |
| left Postsubiculum | -0.0186 | -1.1116 | 4.65E-11 | 0.3941 |
| left Lateral mammillary nucleus | -0.0042 | -1.1067 | 4.87E-11 | 0.3959 |
| left Paragigantocellular reticular nucleus, lateral part | -0.0161 | -1.1047 | 4.99E-11 | 0.3959 |
| left Posterior pretectal nucleus | -0.0032 | -1.1036 | 5.12E-11 | 0.3959 |
| right Supraoculomotor periaqueductal gray | -0.0010 | -1.1034 | 5.24E-11 | 0.3959 |
| right cuneate fascicle | -0.0015 | -1.1000 | 5.37E-11 | 0.3971 |
| right Area prostriata | -0.0079 | -1.0962 | 5.50E-11 | 0.3985 |
| left Vestibulocerebellar nucleus | -0.0034 | -1.0902 | 5.93E-11 | 0.4002 |
| left Field CA2 | -0.0135 | -1.0895 | 5.78E-11 | 0.4002 |
| right Medial geniculate complex, ventral part | -0.0068 | -1.0790 | 6.08E-11 | 0.4055 |
| left Lingula (I) | -0.0051 | -1.0771 | 6.24E-11 | 0.4058 |
| right Tuberal nucleus | -0.0130 | -1.0739 | 6.40E-11 | 0.4069 |
| right Claustrum | -0.0148 | -1.0691 | 6.73E-11 | 0.4087 |
| left trapezoid body | -0.0061 | -1.0683 | 6.56E-11 | 0.4087 |
| left Spinal nucleus of the trigeminal, oral part | -0.0199 | -1.0669 | 6.91E-11 | 0.4088 |
| left Globus pallidus, external segment | -0.0190 | -1.0480 | 7.09E-11 | 0.4190 |
| left fimbria | -0.0526 | -1.0414 | 7.27E-11 | 0.4221 |
| right lateral recess | -0.0119 | -1.0394 | 7.46E-11 | 0.4226 |
| left pyramid | -0.0099 | -1.0360 | 7.87E-11 | 0.4231 |
| left Sublaterodorsal nucleus | -0.0008 | -1.0325 | 8.30E-11 | 0.4240 |

| | | | | |
|---|---|---|---|---|
| left Lateral septal nucleus, ventral part | -0.0218 | -1.0320 | 8.08E-11 | 0.4240 |
| right Nucleus of the solitary tract | -0.0223 | -1.0273 | 8.52E-11 | 0.4260 |
| right Precommissural nucleus | -0.0058 | -1.0229 | 8.76E-11 | 0.4279 |
| right Nucleus ambiguus, ventral division | -0.0009 | -1.0206 | 9.00E-11 | 0.4285 |
| left spinal tract of the trigeminal nerve | -0.0248 | -1.0175 | 9.25E-11 | 0.4292 |
| right Globus pallidus, internal segment | -0.0085 | -1.0169 | 9.50E-11 | 0.4292 |
| right Intermediate reticular nucleus | -0.0635 | -1.0148 | 9.77E-11 | 0.4297 |
| right Medial pretectal area | -0.0012 | -1.0133 | 1.00E-10 | 0.4299 |
| right Lateral reticular nucleus, parvicellular part | -0.0015 | -0.9990 | 1.03E-10 | 0.4377 |
| left facial nerve | -0.0014 | -0.9962 | 1.06E-10 | 0.4380 |
| cerebellar commissure | -0.0051 | -0.9954 | 1.12E-10 | 0.4380 |
| left Medial accesory oculomotor nucleus | -0.0009 | -0.9947 | 1.09E-10 | 0.4380 |
| right Field CA3 | -0.0698 | -0.9887 | 1.16E-10 | 0.4409 |
| left Xiphoid thalamic nucleus | -0.0030 | -0.9796 | 1.19E-10 | 0.4456 |
| right Ventral tegmental nucleus | -0.0006 | -0.9731 | 1.22E-10 | 0.4488 |
| left Lateral vestibular nucleus | -0.0065 | -0.9711 | 1.26E-10 | 0.4493 |
| left Presubiculum | -0.0208 | -0.9691 | 1.30E-10 | 0.4497 |
| right Fasciola cinerea | -0.0011 | -0.9653 | 1.34E-10 | 0.4513 |
| left Subparafascicular area | -0.0035 | -0.9614 | 1.38E-10 | 0.4529 |
| left lateral ventricle | -0.1085 | -0.9535 | 1.42E-10 | 0.4570 |
| left Globus pallidus, internal segment | -0.0063 | -0.9421 | 1.46E-10 | 0.4632 |
| right Trochlear nucleus | -0.0003 | -0.9397 | 1.51E-10 | 0.4640 |
| left ventral spinocerebellar tract | -0.0039 | -0.9304 | 1.55E-10 | 0.4690 |
| right Accessory trigeminal nucleus | -0.0006 | -0.9285 | 1.60E-10 | 0.4694 |
| right Spinal nucleus of the trigeminal, oral part | -0.0191 | -0.9165 | 1.87E-10 | 0.4754 |
| left olfactory nerve layer of main olfactory bulb | -0.0478 | -0.9159 | 1.81E-10 | 0.4754 |
| left Ventral tegmental nucleus | -0.0008 | -0.9134 | 1.75E-10 | 0.4754 |
| left optic tract | -0.0105 | -0.9131 | 1.70E-10 | 0.4754 |
| left Frontal pole | -0.0182 | -0.9128 | 1.65E-10 | 0.4754 |

| | | | | |
|---|---|---|---|---|
| left Bed nucleus of the accessory olfactory tract | -0.0008 | -0.9069 | 1.93E-10 | 0.4783 |
| left dorsal spinocerebellar tract | -0.0031 | -0.9019 | 1.99E-10 | 0.4807 |
| right Linear nucleus of the medulla | -0.0019 | -0.8916 | 2.12E-10 | 0.4856 |
| left cingulum bundle | -0.0132 | -0.8822 | 2.26E-10 | 0.4908 |
| left Parabigeminal nucleus | -0.0008 | -0.8811 | 2.19E-10 | 0.4908 |
| right spinal tract of the trigeminal nerve | -0.0292 | -0.8791 | 2.34E-10 | 0.4912 |
| right Tuberomammillary nucleus, ventral part | -0.0028 | -0.8674 | 2.42E-10 | 0.4980 |
| left Precommissural nucleus | -0.0048 | -0.8655 | 2.50E-10 | 0.4984 |
| left lateral lemniscus | -0.0123 | -0.8626 | 2.58E-10 | 0.4995 |
| right Inferior olivary complex | -0.0096 | -0.8545 | 2.76E-10 | 0.5035 |
| Median preoptic nucleus | -0.0015 | -0.8540 | 2.67E-10 | 0.5035 |
| right mammillary peduncle | -0.0010 | -0.8467 | 3.40E-10 | 0.5054 |
| right Oculomotor nucleus | -0.0009 | -0.8458 | 3.17E-10 | 0.5054 |
| left Taenia tecta, dorsal part | -0.0124 | -0.8455 | 2.96E-10 | 0.5054 |
| left Copula pyramidis | -0.0451 | -0.8448 | 3.06E-10 | 0.5054 |
| left Tuberomammillary nucleus, ventral part | -0.0021 | -0.8447 | 2.86E-10 | 0.5054 |
| right Dorsal terminal nucleus of the accessory optic tract | -0.0003 | -0.8436 | 3.29E-10 | 0.5054 |
| right choroid plexus | -0.0331 | -0.8416 | 3.53E-10 | 0.5059 |
| left Ventrolateral preoptic nucleus | -0.0014 | -0.8396 | 3.66E-10 | 0.5064 |
| right Posterodorsal preoptic nucleus | -0.0003 | -0.8260 | 3.93E-10 | 0.5140 |
| right Dorsal cochlear nucleus | -0.0155 | -0.8255 | 3.79E-10 | 0.5140 |
| right Nucleus of Darkschewitsch | -0.0014 | -0.8223 | 4.08E-10 | 0.5152 |
| right Nucleus x | -0.0010 | -0.8197 | 4.23E-10 | 0.5162 |
| left Nucleus incertus | -0.0020 | -0.8085 | 4.39E-10 | 0.5225 |
| left Paratrochlear nucleus | -0.0006 | -0.8069 | 4.56E-10 | 0.5225 |
| Subfornical organ | -0.0009 | -0.8051 | 4.73E-10 | 0.5225 |
| right Lateral septal nucleus, ventral part | -0.0179 | -0.8050 | 4.92E-10 | 0.5225 |
| right Fastigial nucleus | -0.0115 | -0.8036 | 5.11E-10 | 0.5227 |
| right Lateral vestibular nucleus | -0.0048 | -0.7965 | 5.31E-10 | 0.5266 |
| right Ventrolateral preoptic nucleus | -0.0015 | -0.7934 | 5.52E-10 | 0.5278 |

| | | | | |
|---|---|---|---|---|
| left Parastrial nucleus | -0.0010 | -0.7801 | 5.97E-10 | 0.5356 |
| left Medial pretectal area | -0.0007 | -0.7792 | 5.74E-10 | 0.5356 |
| right Magnocellular nucleus | -0.0069 | -0.7755 | 6.22E-10 | 0.5373 |
| left Fastigial nucleus | -0.0117 | -0.7705 | 6.74E-10 | 0.5391 |
| left Main olfactory bulb | -0.1435 | -0.7697 | 6.47E-10 | 0.5391 |
| left Posterodorsal tegmental nucleus | -0.0009 | -0.7691 | 7.02E-10 | 0.5391 |
| right Spinal vestibular nucleus | -0.0117 | -0.7615 | 7.95E-10 | 0.5427 |
| left Superior olivary complex, medial part | -0.0034 | -0.7605 | 7.31E-10 | 0.5427 |
| Interpeduncular nucleus | -0.0223 | -0.7600 | 7.62E-10 | 0.5427 |
| habenular commissure | -0.0013 | -0.7475 | 8.29E-10 | 0.5503 |
| right Nucleus of the trapezoid body | -0.0022 | -0.7369 | 8.65E-10 | 0.5567 |
| Lobule III | -0.1856 | -0.7271 | 9.85E-10 | 0.5623 |
| left Spinal nucleus of the trigeminal, interpolar part | -0.0406 | -0.7250 | 9.43E-10 | 0.5623 |
| left Accessory olfactory bulb | -0.0093 | -0.7249 | 9.03E-10 | 0.5623 |
| right Posterolateral visual area | -0.0151 | -0.7165 | 1.03E-09 | 0.5672 |
| right Nucleus prepositus | -0.0030 | -0.7104 | 1.08E-09 | 0.5706 |
| right Nucleus y | -0.0008 | -0.7014 | 1.18E-09 | 0.5750 |
| left Anterodorsal nucleus | -0.0024 | -0.6983 | 1.23E-09 | 0.5764 |
| left subependymal zone | -0.0032 | -0.6907 | 1.29E-09 | 0.5801 |
| right Postsubiculum | -0.0130 | -0.6904 | 1.35E-09 | 0.5801 |
| left amygdalar capsule | -0.0022 | -0.6866 | 1.42E-09 | 0.5810 |
| right Paratrigeminal nucleus | -0.0027 | -0.6866 | 1.49E-09 | 0.5810 |
| right Retrochiasmatic area | -0.0024 | -0.6842 | 1.56E-09 | 0.5818 |
| right vomeronasal nerve | -0.0006 | -0.6813 | 1.64E-09 | 0.5827 |
| right Paraflocculus | -0.0418 | -0.6804 | 1.72E-09 | 0.5827 |
| right fimbria | -0.0353 | -0.6769 | 1.80E-09 | 0.5842 |
| right Superior olivary complex, medial part | -0.0023 | -0.6720 | 1.99E-09 | 0.5865 |
| right Lateral mammillary nucleus | -0.0028 | -0.6690 | 2.10E-09 | 0.5865 |
| left Area postrema | -0.0015 | -0.6687 | 1.90E-09 | 0.5865 |
| left Dorsal terminal nucleus of the accessory optic tract | -0.0003 | -0.6569 | 2.45E-09 | 0.5930 |
| right inferior cerebellar peduncle | -0.0105 | -0.6555 | 2.58E-09 | 0.5931 |

| | | | | |
|---|---|---|---|---|
| right Facial motor nucleus | -0.0090 | -0.6343 | 2.88E-09 | 0.6062 |
| left External cuneate nucleus | -0.0055 | -0.6293 | 3.04E-09 | 0.6088 |
| Lobules IV-V | -0.2503 | -0.6233 | 3.21E-09 | 0.6122 |
| left Parasubiculum | -0.0149 | -0.6188 | 3.39E-09 | 0.6145 |
| right Paragigantocellular reticular nucleus, dorsal part | -0.0033 | -0.6115 | 3.80E-09 | 0.6181 |
| right lateral ventricle | -0.0738 | -0.6112 | 3.59E-09 | 0.6181 |
| right Supragenual nucleus | -0.0003 | -0.6019 | 4.53E-09 | 0.6229 |
| left Gustatory areas | -0.0135 | -0.5988 | 4.81E-09 | 0.6234 |
| right Parvicellular reticular nucleus | -0.0267 | -0.5871 | 5.11E-09 | 0.6308 |
| right Parasolitary nucleus | -0.0012 | -0.5827 | 5.78E-09 | 0.6326 |
| left Nucleus of the lateral lemniscus | -0.0076 | -0.5820 | 5.43E-09 | 0.6326 |
| left Parapyramidal nucleus | -0.0010 | -0.5749 | 6.16E-09 | 0.6369 |
| right olfactory nerve layer of main olfactory bulb | -0.0335 | -0.5531 | 7.00E-09 | 0.6508 |
| Triangular nucleus of septum | -0.0077 | -0.5329 | 7.98E-09 | 0.6637 |
| right Bed nucleus of the accessory olfactory tract | -0.0004 | -0.5284 | 8.54E-09 | 0.6661 |
| left Nucleus ambiguus, dorsal division | -0.0004 | -0.5262 | 9.14E-09 | 0.6667 |
| right Subceruleus nucleus | -0.0003 | -0.5216 | 9.79E-09 | 0.6691 |
| left optic nerve | -0.0009 | -0.5178 | 1.05E-08 | 0.6710 |
| right Lateral septal nucleus, caudal (caudodorsal) part | -0.0084 | -0.5128 | 1.13E-08 | 0.6737 |
| right Intermediate geniculate nucleus | -0.0003 | -0.4918 | 1.21E-08 | 0.6883 |
| right facial nerve | -0.0011 | -0.4900 | 1.30E-08 | 0.6887 |
| right supraoptic commissures | -0.0003 | -0.4834 | 1.40E-08 | 0.6926 |
| left Dorsal cochlear nucleus | -0.0081 | -0.4626 | 1.76E-08 | 0.7052 |
| Septofimbrial nucleus | -0.0106 | -0.4420 | 1.91E-08 | 0.7197 |
| right medial longitudinal fascicle | -0.0017 | -0.4386 | 2.07E-08 | 0.7213 |
| right Barrington's nucleus | -0.0003 | -0.4317 | 2.24E-08 | 0.7248 |
| right Parvicellular motor 5 nucleus | -0.0005 | -0.4313 | 2.43E-08 | 0.7248 |
| left Lateral septal nucleus, rostral (rostroventral) part | -0.0330 | -0.4289 | 2.65E-08 | 0.7256 |
| left trochlear nerve | -0.0002 | -0.4012 | 3.14E-08 | 0.7446 |
| right Posterodorsal tegmental nucleus | -0.0004 | -0.3976 | 3.43E-08 | 0.7463 |

| | | | | |
|---|---|---|---|---|
| left uncinate fascicle | -0.0009 | -0.3886 | 3.76E-08 | 0.7522 |
| left Simple lobule | -0.0668 | -0.3837 | 4.12E-08 | 0.7549 |
| left Anterior tegmental nucleus | -0.0003 | -0.3750 | 4.52E-08 | 0.7605 |
| right Sublaterodorsal nucleus | -0.0003 | -0.3562 | 5.48E-08 | 0.7729 |
| right trapezoid body | -0.0012 | -0.3468 | 6.05E-08 | 0.7791 |
| left Accessory trigeminal nucleus | -0.0002 | -0.3449 | 6.69E-08 | 0.7795 |
| left medial corticohypothalamic tract | -0.0002 | -0.3371 | 8.24E-08 | 0.7834 |
| right Ventral cochlear nucleus | -0.0066 | -0.3273 | 1.02E-07 | 0.7887 |
| left lateral recess | -0.0033 | -0.2959 | 1.28E-07 | 0.8110 |
| right medial corticohypothalamic tract | -0.0002 | -0.2943 | 1.44E-07 | 0.8111 |
| left Anterior olfactory nucleus | -0.0164 | -0.2837 | 1.63E-07 | 0.8183 |
| left Retrosplenial area, ventral part | -0.0220 | -0.2727 | 1.84E-07 | 0.8258 |
| right Lateral septal nucleus, rostral (rostroventral) part | -0.0160 | -0.2459 | 2.09E-07 | 0.8456 |
| right Ventromedial preoptic nucleus | -0.0003 | -0.2354 | 2.38E-07 | 0.8528 |
| right Paratrochlear nucleus | -0.0002 | -0.2273 | 2.72E-07 | 0.8580 |
| right Intertrigeminal nucleus | -0.0003 | -0.2101 | 3.11E-07 | 0.8704 |
| Interfascicular nucleus raphe | -0.0007 | -0.2076 | 3.58E-07 | 0.8712 |
| right Spinal nucleus of the trigeminal, interpolar part | -0.0113 | -0.1967 | 4.80E-07 | 0.8775 |
| optic chiasm | -0.0029 | -0.1807 | 5.60E-07 | 0.8889 |
| right Lateral terminal nucleus of the accessory optic tract | -0.0001 | -0.1450 | 1.09E-06 | 0.9132 |
| left vomeronasal nerve | -0.0001 | -0.1439 | 9.14E-07 | 0.9132 |
| left Ventromedial preoptic nucleus | -0.0002 | -0.1362 | 1.30E-06 | 0.9170 |
| superior colliculus commissure | -0.0003 | -0.1361 | 1.57E-06 | 0.9170 |
| left vestibular nerve | -0.0010 | -0.1317 | 1.91E-06 | 0.9187 |
| left Barrington's nucleus | -0.0001 | -0.1220 | 2.89E-06 | 0.9228 |
| left Abducens nucleus | -0.0001 | -0.1209 | 3.60E-06 | 0.9228 |
| right Olivary pretectal nucleus | -0.0002 | -0.1209 | 4.54E-06 | 0.9228 |
| left sensory root of the trigeminal nerve | -0.0008 | -0.1167 | 5.78E-06 | 0.9250 |
| right commissural branch of stria terminalis | -0.0001 | -0.1143 | 7.44E-06 | 0.9256 |
| inferior colliculus commissure | -0.0001 | -0.1106 | 9.72E-06 | 0.9272 |

| | | | | |
|---|---|---|---|---|
| left Facial motor nucleus | -0.0015 | -0.1079 | 1.29E-05 | 0.9281 |
| Interanteromedial nucleus of the thalamus | -0.0002 | -0.0724 | 1.08E-04 | 0.9485 |
| right Nucleus incertus | -0.0001 | -0.0487 | 1.71E-04 | 0.9658 |
| right optic nerve | -0.0001 | -0.0409 | 2.83E-04 | 0.9706 |
| left middle cerebellar peduncle | 0.0000 | -0.0002 | 9.17E-04 | 0.9999 |
| ventral hippocampal commissure | 0.0001 | 0.0305 | 4.93E-04 | 0.9775 |
| right Field CA2 | 0.0008 | 0.0763 | 7.09E-05 | 0.9468 |
| left Nucleus y | 0.0001 | 0.0855 | 4.80E-05 | 0.9408 |
| right vestibular nerve | 0.0006 | 0.0907 | 3.34E-05 | 0.9380 |
| left Fasciola cinerea | 0.0002 | 0.0929 | 2.38E-05 | 0.9374 |
| left Paranigral nucleus | 0.0001 | 0.1023 | 1.74E-05 | 0.9313 |
| left Paraflocculus | 0.0068 | 0.1308 | 2.34E-06 | 0.9187 |
| left dorsal limb | 0.0006 | 0.1469 | 7.72E-07 | 0.9132 |
| left Olivary pretectal nucleus | 0.0003 | 0.1719 | 6.56E-07 | 0.8947 |
| left Lateral terminal nucleus of the accessory optic tract | 0.0002 | 0.2018 | 4.14E-07 | 0.8746 |
| left Anteroventral periventricular nucleus | 0.0010 | 0.3124 | 1.14E-07 | 0.7992 |
| dorsal fornix | 0.0005 | 0.3286 | 9.17E-08 | 0.7887 |
| right trochlear nerve | 0.0001 | 0.3421 | 7.41E-08 | 0.7806 |
| left Nucleus of the trapezoid body | 0.0009 | 0.3593 | 4.97E-08 | 0.7716 |
| left Superior olivary complex, lateral part | 0.0027 | 0.4220 | 2.88E-08 | 0.7298 |
| left Midbrain trigeminal nucleus | 0.0003 | 0.4769 | 1.63E-08 | 0.6955 |
| left supraoptic commissures | 0.0003 | 0.4813 | 1.51E-08 | 0.6932 |
| left inferior cerebellar peduncle | 0.0075 | 0.5418 | 7.47E-09 | 0.6581 |
| left Superior olivary complex, periolivary region | 0.0030 | 0.5569 | 6.56E-09 | 0.6490 |
| left Flocculus | 0.0159 | 0.6008 | 4.27E-09 | 0.6229 |
| right Locus ceruleus | 0.0004 | 0.6034 | 4.03E-09 | 0.6228 |
| right Accessory supraoptic group | 0.0002 | 0.6525 | 2.72E-09 | 0.5943 |
| right Abducens nucleus | 0.0007 | 0.6675 | 2.21E-09 | 0.5865 |
| right External cuneate nucleus | 0.0047 | 0.6697 | 2.33E-09 | 0.5865 |
| left Trochlear nucleus | 0.0002 | 0.7070 | 1.13E-09 | 0.5720 |
| right dorsal acoustic stria | 0.0006 | 0.8969 | 2.05E-10 | 0.4831 |

| | | | | |
|---|---|---|---|---|
| left Accessory supraoptic group | 0.0003 | 1.0373 | 7.66E-11 | 0.4230 |
| left Inferior salivatory nucleus | 0.0006 | 1.0928 | 5.64E-11 | 0.3998 |
| right Supraoptic nucleus | 0.0024 | 1.2547 | 3.08E-11 | 0.3314 |
| left Ventral cochlear nucleus | 0.0299 | 1.5294 | 8.79E-12 | 0.2466 |

**Supplementary Table 2. IHC results.** The table describes differences in staining in terms of number of DAB positive markers (cells for GFAP, Iba-1, and TH; pixels for pS129Syn) normalised by tissue area in regions for four stains of interest (GFAP (astrocyte marker), Iba-1 (microglial marker), pS129Syn (phosphorylated aSyn marker), and TH (dopaminergic marker) at FDR 5% threshold (such that q<0.05). Yellow highlights represent regions identified in the structural covariance pattern, blue highlights represent regions where significant whole structure volume differences were observed, and green highlights represent regions in both MRI analyses. Injection site is denoted by the red border. All significant tests were in one direction; higher staining in PFF- compared to PBS-injected mice. Abbreviations: GFAP, glial fibrillary acidic protein; Iba-1, ionised calcium-binding adapter molecule 1; pS129Syn, phosphorylated Serine129 alpha-synuclein; TH, tyrosine hydroxylase; 1MC, primary motor cortex;1SS, primary somatosensory cortex; MRN, midbrain reticular nucleus; NAc, nucleus accumbens; PAG, periaqueductal grey; PVN, paraventricular hypothalamic nucleus; SN, substantia nigra; ns, non-significant; NA, not applicable.

| hemisphere | region | GFAP | IBA1 | pS129Syn | TH |
|---|---|---|---|---|---|
| left | 1MC | ns | ns | 0.030 | NA |
| right | 1MC | 0.008 | 0.027 | 0.007 | NA |
| left | 1SS (anterior) | ns | ns | ns | NA |
| right | 1SS (anterior) | 0.044 | ns | ns | NA |
| left | 1SS (posterior) | ns | ns | 0.050 | NA |
| right | 1SS (posterior) | 0.033 | ns | ns | NA |
| left | hippocampus | ns | ns | ns | NA |
| right | hippocampus | ns | ns | ns | NA |
| left | hypothalamus | ns | ns | ns | NA |
| right | hypothalamus | ns | ns | 0.010 | NA |
| medial | medulla (superior) | 0.019 | 0.045 | 0.023 | NA |
| medial | medulla (inferior) | 0.028 | 0.018 | 0.017 | NA |
| left | MRN | 0.014 | 0.041 | 0.027 | NA |
| right | MRN | 0.017 | 0.014 | ns | NA |
| left | NAc | 0.050 | ns | 0.033 | ns |
| right | NAc | 0.036 | ns | 0.040 | ns |
| medial | PAG | 0.006 | 0.005 | 0.003 | ns |
| left | pons | 0.042 | 0.036 | ns | ns |
| right | pons | ns | ns | 0.037 | ns |
| medial | PVN (posterior) | ns | 0.023 | ns | NA |
| left | SN | 0.022 | ns | ns | ns |
| right | SN | 0.011 | 0.009 | ns | ns |
| left | striatum | 0.025 | ns | ns | ns |
| right | striatum (inj. site) | 0.003 | 0.050 | 0.047 | ns |
| left | thalamus (anterior) | 0.039 | ns | 0.013 | NA |
| right | thalamus (anterior) | ns | ns | 0.043 | NA |
| left | thalamus (posterior) | 0.031 | ns | 0.020 | NA |
| right | thalamus (posterior) | ns | ns | ns | NA |
